# Deep Learning of Brain-Behavior Dimensions Identifies Transdiagnostic Biotypes in Youth with ADHD and Anxiety Disorders

**DOI:** 10.1101/2025.10.13.682243

**Authors:** Yong Jiao, Xiaoyu Tong, Gregory A. Fonzo, Ian H. Gotlib, Kilian M. Pohl, Theodore D. Satterthwaite, Jing Jiang, Yu Zhang

## Abstract

Attention-deficit/hyperactivity disorder and anxiety disorders are highly prevalent in youth and are characterized by substantial heterogeneity and frequent co-occurrence. This transdiagnostic complexity challenges conventional diagnostic frameworks that rely on symptom-based categories, which often obscure underlying dimensional and neurobiological mechanisms and offer limited neurobiological specificity. To address these issues, we developed a deep learning-based brain–behavior modeling framework that integrates clinically salient functional connectivity with cognitive and behavioral measures to identify interpretable dimensions and biologically grounded subtypes (biotypes). We applied our model to the Adolescent Brain Cognitive Development (ABCD) dataset comprising 3,508 children aged 9–11 years and revealed two reproducible brain–behavior dimensions that captured variation in cognitive control and emotion–attention regulation. These dimensions further yielded three distinct biotypes, each exhibiting unique symptom profiles and distinct brain development. We tested the robustness and generalizability of the dimensions and corresponding biotypes in an independent cohort of 224 age-matched participants from the Healthy Brain Network (HBN) and documented their early expression before symptom onset during adolescence. These findings highlight the utility of brain–behavior dimensions for elucidating heterogeneous psychiatric presentations and advance a biologically grounded framework for early classification and potential clinical translation in youth mental health.

## Introduction

Attention-deficit/hyperactivity disorder (ADHD) and anxiety disorders (AXD) are highly prevalent in youth^1^. Furthermore, both exhibit pronounced clinical and neurobiological heterogeneity in symptom presentation, developmental symptom trajectories, and treatment outcomes. Beyond the diversity of each disorder, ADHD and AXD frequently co-occur, blurring diagnostic boundaries and complicating treatment selection. Disruptions in cognitive control and emotional regulation are central to both disorders. This combination of within-disorder heterogeneity and cross-disorder comorbidity is difficult to capture with categorical diagnostic systems (e.g., DSM-5^2^), which impose thresholded categories defined by symptom counts and duration criteria, and rely on clinician interpretation, potentially obscuring shared mechanisms and contributing to inconsistent identification of individualized treatment targets across clinical and research contexts^3,4^. Given these limitations, it remains unclear whether such transdiagnostic heterogeneity and comorbidity reflect underlying neurobiological patterns that can be systematically characterized. Self-report questionnaires, while clinically informative, often show limited correspondence with brain abnormalities, yielding categories that show only poor alignment with neuroimaging measures. This results in insufficient neurobiological specificity for precise characterization and stratification. These difficulties highlight the need for dimensional representations as compact, continuous axes with individual scores learned from multivariate data that integrate cognitive and clinical information and map directly onto underlying neurobiological variation.

To link brain to behavior, functional connectivity (FC) quantifies interregional coordination and captures reliable individual differences in brain function. In youth, prior studies have reported group-level alterations in FC and potential biomarkers linked to cognition and psychiatric symptoms^5–8^. Resting-state fMRI (rs-fMRI) provides a task-free, scalable approach to estimate FC and relate brain to behavior. However, disentangling clinically meaningful subtypes remains challenging given the heterogeneity of ADHD and AXD. To address this challenge, recent efforts have shifted toward data-driven approaches using rs-fMRI to uncover neurobiologically informed brain–behavior associations that transcend conventional diagnostic categories, yielding a more precise and biologically grounded understanding of psychiatric conditions^9,10^. Among these approaches, multivariate methods such as canonical correlation analysis (CCA) and partial least squares (PLS) have shown promise in linking FC to cognitive and behavioral measures^11–13^, moving beyond group-level comparisons by capturing latent transdiagnostic factors. However, the effectiveness of these multivariate methods in identifying behavior-aligned dimensions remains constrained by the high dimensionality of neuroimaging data and the shared variance between patients and controls. Conventional dimensionality reduction techniques, such as univariate feature selection or principal component analysis (PCA), have limited ability to isolate clinically relevant effects, motivating integrative methods that learn compact, behavior-aligned representations.

To address these limitations, contrastive learning has emerged as a principled framework for isolating disorder-specific signals from normative variability^14^. Sequential approaches, such as contrastive PCA followed by sparse CCA, have demonstrated feasibility for extracting clinically relevant features and strengthening associations with symptom severity^15,16^. Beyond linear approaches, contrastive variational autoencoders (cVAEs)^17^ highlight the potential of nonlinear representation learning for disorder-specific feature extraction in neuroimaging^18^. Building on these advances, we propose a **De**ep learning framework for **Co**ntrastive **D**imensional **E**mbedding (DeCoDE), which unifies contrastive feature learning with brain–behavior CCA to identify transdiagnostic dimensions, yielding compact FC representations (Fig. 1). By disentangling ADHD/AXD-specific patterns of FC from normative variation via cVAEs while simultaneously aligning these representations with cognitive and behavioral profiles via Deep Generalized Canonical Correlation Analysis (DGCCA)^19^, DeCoDE enables precise mapping of brain–behavior relations. Applied to 3,508 ABCD youth with ADHD or AXD and validated in 224 age-matched HBN participants, DeCoDE uncovered reproducible transdiagnostic dimensions capturing symptom heterogeneity. Projecting individuals into this low-dimensional neurobiological space further revealed neurophysiological subtypes (biotypes) that transcend diagnostic boundaries, establishing a data-driven foundation for biologically grounded subtyping in youth psychiatry^20^.

**Figure 1.**
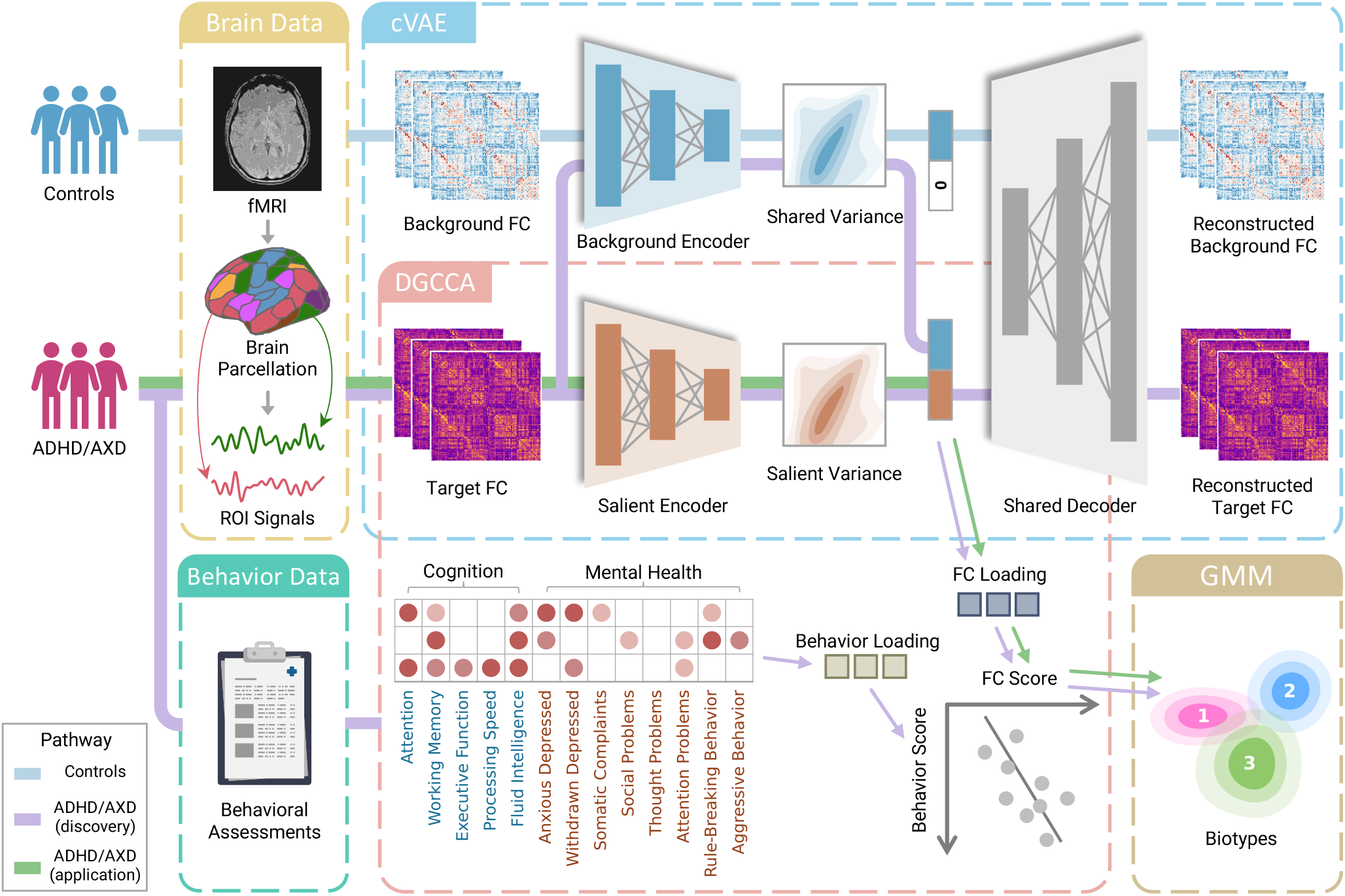
Framework for identifying transdiagnostic brain–behavior dimensions and biotypes in youth with ADHD and AXD. *Discovery phase:* Resting-state fMRI from ADHD/AXD and controls is parcellated into regional time series to compute functional connectivity (FC) for all participants. Behavioral assessments (cognitive tasks and clinical scales) are obtained for ADHD/AXD participants. Within DeCoDE, a contrastive variational autoencoder (cVAE) disentangles ADHD/AXD-specific latent representations from shared variability. Deep generalized CCA (DGCCA) aligns cVAE-derived FC features with ADHD/AXD behavioral measures in a shared latent space, yielding canonical FC and behavior scores with interpretable loadings. The cVAE salient encoder also serves as the feature extractor for DGCCA. Gaussian mixture modeling (GMM) on FC scores delineates neurophysiologically informed biotypes. *Application phase:* For new ADHD/AXD cases, FC alone is processed with the learned salient encoder, FC loadings, and GMM from the discovery phase to assign biotypes without additional behavioral assessments.

## Results

We identified two brain–behavior dimensions from ADHD/AXD-specific functional connectivity, which delineated three transdiagnostic biotypes with distinct neural and clinical profiles. We validated these findings in an independent cohort and benchmarked them against diagnostic groupings to assess neurobiological specificity.

### Contrastive FC defines robust, transdiagnostic brain–behavior dimensions

We applied DeCoDE to a transdiagnostic cohort of 3,508 youth with ADHD or AXD, contrasted with 1,947 controls from the ABCD dataset (Method 1; demographic and clinical characteristics are summarized in Supplementary Table S1). The extracted ADHD/AXD-specific FC representations served as the brain view in DGCCA, and the behavioral view comprised four NIH Toolbox subtests^21^, fluid intelligence from the WISC-V^22^, and eight parent-reported CBCL subscales^23^, covering broad domains of cognition and psychopathology (Method 2).

DeCoDE identified two robust and generalizable brain–behavior dimensions through 10-fold cross-validation (Method 8). The first dimension (cross-validated canonical correlation *r*_cv_ = 0.412, *p <* 0.0001, Cohen’s *d* = 0.905, Fig. 2 a) represented a Cognitive–Behavioral Dysfunction axis, reflecting a co-aggregation of cognitive capacity and behavioral regulation, with strong positive loadings on fluid intelligence, working memory, and executive function, and negative loadings on rule-breaking behavior. Individuals with lower scores on this dimension had higher levels of externalizing symptoms and poorer cognitive performance (Fig. 2 c). The second dimension (*r*_cv_ = 0.212, *p <* 0.0001, Cohen’s *d* = 0.435, Fig. 2 b) captured an Emotion–Attention Dysfunction axis, with positive loadings on anxious/depressed symptoms and negative loadings on attention and thought problems (Fig. 2 d). Higher scores corresponded to more severe internalizing symptoms; conversely, lower scores to greater attention problems. The two dimensions were mutually uncorrelated in both FC and behavior score spaces (Supplementary Fig. S1). No sex differences were observed (Supplementary Fig. S2a). We assessed reliability using the intraclass correlation (ICC) of canonical loadings across folds. Both FC and behavior loadings demonstrated excellent stability, confirming robust brain–behavior mappings (Cognitive-Behavioral Dysfunction: FC ICC = 0.89, behavior ICC = 0.99; Emotion–Attention Dysfunction: FC ICC = 0.91, behavior ICC = 0.99; Supplementary Fig. S3). Ablations showed that removing the integrative modeling, contrastive/variational components, or the nonlinear multiview embedding reduced correlations or impaired cross-modal alignment (Supplementary Figs.S4–S8). An overview across methods is provided in Supplementary Fig. S9.

**Figure 2.**
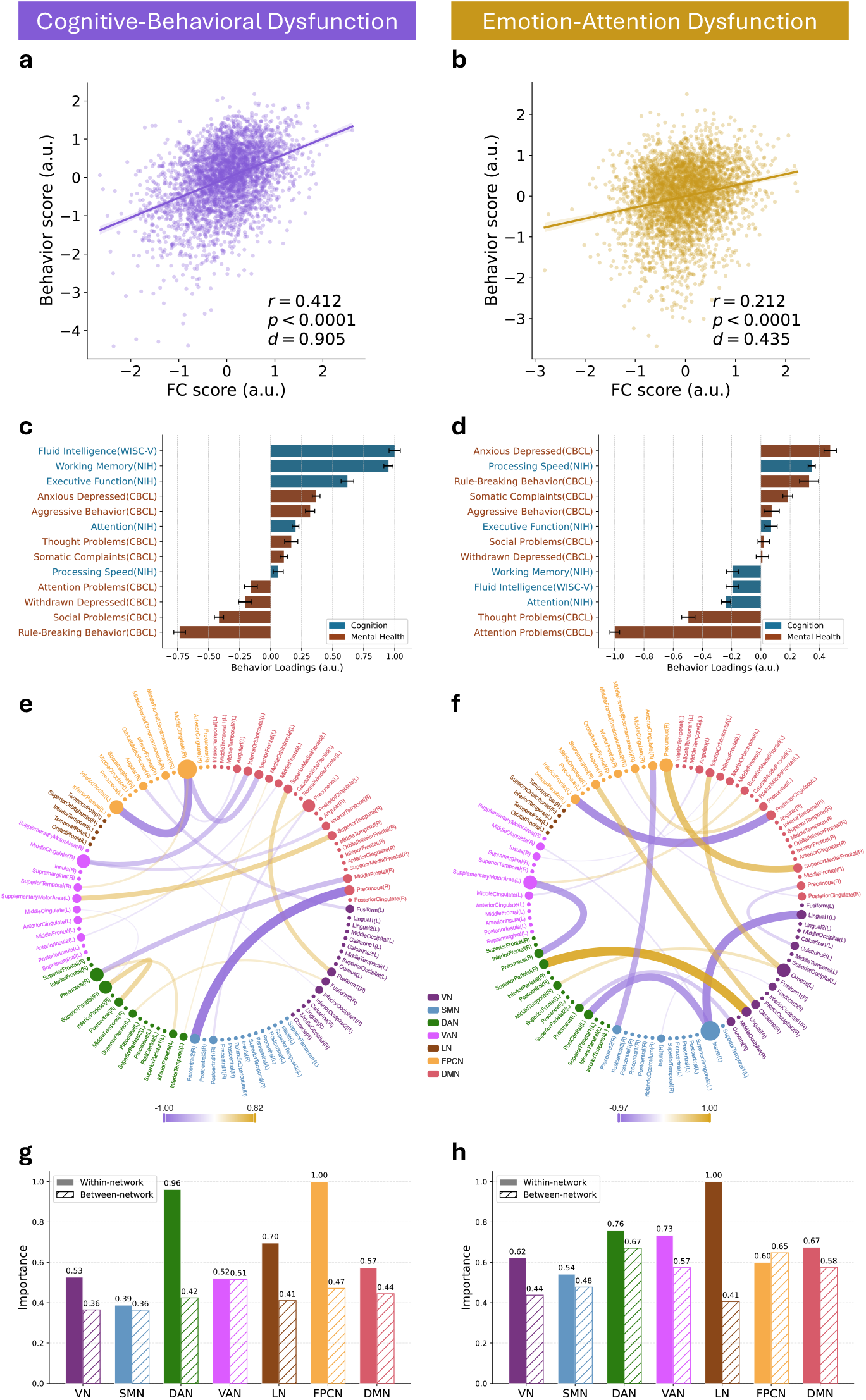
Canonical dimensions linking functional connectivity and behavior measures in youth with ADHD and AXD. **a,b**, Correlation between FC scores and behavior scores for the Cognitive–Behavioral Dysfunction (**a**) and Emotion–Attention Dysfunction (**b**) dimensions across 3,508 youth with 10-fold cross-validation. **c,d**, Behavior loadings of cognitive tasks and CBCL subscales; error bars indicate standard deviation across folds. The Cognitive–Behavioral Dysfunction dimension (**c**) shows positive loadings on fluid intelligence, working memory, and executive function, and negative loadings on rule-breaking behavior. The Emotion–Attention Dysfunction (**d**) shows positive loadings on anxious/depressed and negative loadings on attention problems. **e,f**, Top 20 FC loadings estimated with OMP at a sparsity of 500. Node size reflects ROI strength, defined as the mean absolute weight of selected connections incident on each ROI. **g,h**, Within and between functional network importance for the (**g**) Cognitive–Behavioral Dysfunction and (**h**) Emotion–Attention Dysfunction dimensions, computed from absolute OMP-derived FC loadings (all 500 connections) and normalized to 0–1 within each dimension.

DGCCA learns canonical directions for the FC view in latent space, which are not directly interpretable in the native connectome. We therefore approximated FC loadings with orthogonal matching pursuit (OMP^24^; Method 9). A sparsity level of 500 connections was selected based on correlation trends (Supplementary Fig. S10). The top 20 connections by absolute weight, guided by the sharp drop and gap peaks (Supplementary Fig. S11a), are shown in Fig. 2 e–f; all 500 are provided in Supplementary Fig. S11b. Using these 500 connections, we quantified within- and between-network importance (Fig. 2 g–h) and pairwise inter-network importance (Supplementary Fig. S11c) (Method 10). The Cognitive–Behavioral Dysfunction dimension was dominated by within-network connections in the FPCN and DAN, and the right precuneus–precentral connection carried the largest weight. The Emotion–Attention Dysfunction dimension showed prominent within-network contributions from the LN, DAN, and VAN, together with strong DAN between-network effects; the right superior parietal–lingual and left insula–lingual connections were most prominent, with opposite weight signs. ROI strength, defined as the mean absolute weight of selected connections incident on each ROI, highlighted the right middle cingulate for the Cognitive–Behavioral Dysfunction and the left insula for the Emotion–Attention Dysfunction. Together, these findings delineate two robust, transdiagnostic, and behaviorally interpretable brain–behavior dimensions that provide mechanistic insight into the neural organization underlying symptom heterogeneity in ADHD and AXD.

### Contrastive FC dimensions reveal distinct transdiagnostic biotypes

Building on the identified brain–behavior dimensions, we next investigated whether these continuous symptom dimensions in individuals with ADHD or AXD could be translated into clinically meaningful subgroups. We clustered FC scores from the DeCoDE model using a Gaussian mixture model (GMM)^25^ (Method 11). Model selection based on the Bayesian information criterion (BIC), Akaike information criterion (AIC), and cluster stability consistently supported a three-cluster solution (Supplementary Fig. S12), yielding three biotypes among 3,508 youth, displayed in a ternary plot of posterior probabilities (Fig. 3 a) and across DSM categories (Fig. 3 b). A chi-square test further revealed significant distributional differences of DSM categories (ADHD, AXD, comorbidity) across the three biotypes (*χ*^2^ = 48.99, *p* = 5.8 *×* 10^−10^). Proportionally, ADHD and comorbidity were higher in ED and MD, whereas AXD was higher in ID. Inspection of CBCL item-level symptom profiles indicated distinct behavioral signatures (detailed in the next section), on which we based the labels Externalizing–Disinhibited (ED), Internalizing–Distressed (ID), and Mixed Dysregulation (MD). Sociodemographic and clinical characteristics (Supplementary Table S2) showed that ED (12.5% of the sample) was linked to socioeconomic disadvantage, advanced pubertal development, and elevated BMI; ID (35.8%) showed demographic and clinical features most similar to controls; and MD (51.7%) included fewer female participants.

**Figure 3.**
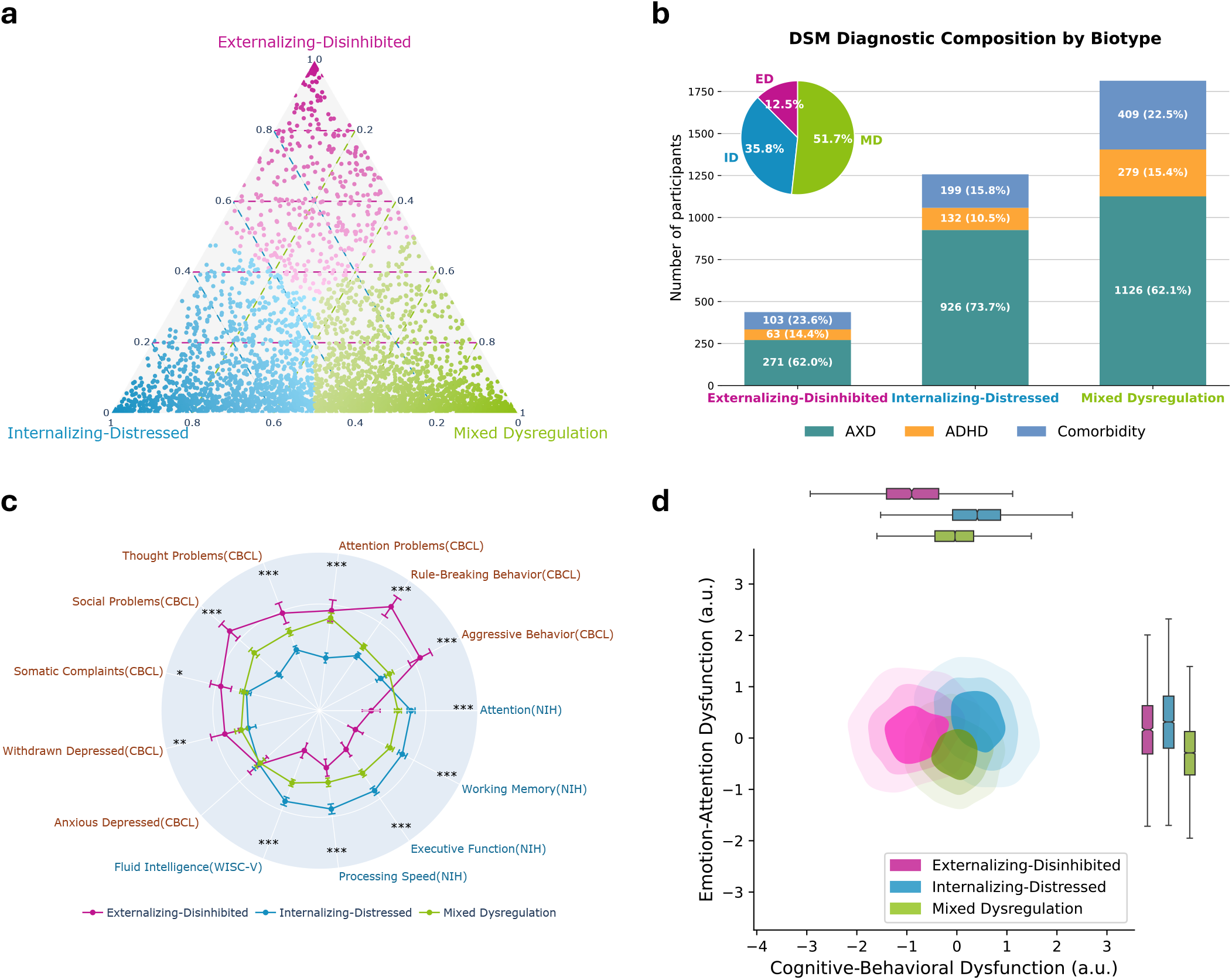
Biotypes derived from canonical FC dimensions in adolescents with ADHD and AXD. **a**, Posterior membership probabilities for 3,508 youth assigned to ED, ID, and MD biotypes via Gaussian mixture modeling of FC scores. Each point represents an individual, with position reflecting their probabilistic assignment to the three biotypes. **b**, DSM diagnostic composition (ADHD, AXD, comorbidity) within each biotype. **c**, Clinical profiles across the five cognitive and eight mental health scales used in model training. Cognitive measures were approximately normally distributed and thus underwent outlier removal using a 3-standard-deviation threshold, whereas mental health scales were non-normally distributed and processed using the Median Absolute Deviation method with a threshold of 4. Group comparisons were conducted using the Kruskal–Wallis test, followed by FDR correction (*q <* 0.05) across scales. Radial positions represent normalized group means; error bars indicate 95% confidence intervals. Asterisks denote scales with significant group differences. ED showed the highest externalizing symptoms and lowest cognitive performance; ID combined high cognitive scores with prominent internalizing symptoms; MD exhibited intermediate severity with marked attentional deficits. **d**, Kernel density estimation plots of biotypes across the two identified brain–behavior dimensions. Density contours (25%, 50%, 75%) illustrate spatial distribution, and marginal boxplots on the top and right summarize score distributions along each axis. ED clustered at low Cognitive–Behavioral Dysfunction scores, ID at high scores on both axes, and MD at low emotion–attention scores.

### Biotypes exhibit distinct cognitive, symptom, and personality profiles

To assess clinical relevance, we examined CBCL item-level profiles in detail (Supplementary Fig. S13). The ED biotype showed the greatest overall severity, dominated by externalizing behaviors such as hostility, defiance, and impulsive tendencies. The ID biotype was characterized by internalizing symptoms, including guilt, tension, and perfectionism, indicative of chronic psychological distress, while the MD biotype displayed heterogeneous features spanning attentional dysregulation, somatic complaints, irritability, and mild self-injury. Group comparisons across the 13 training scales confirmed distinct cognitive–symptom profiles (Fig. 3 c; Supplementary Fig. S14): ED had the lowest cognitive performance and highest levels of social problems, rule-breaking, and aggression; ID showed higher cognition but elevated anxious–depressed symptoms; and MD was intermediate with pronounced attention problems. Extended analyses across additional cognitive, mental health, personality, and contextual measures provide complementary detail (Supplementary Fig. S15; see Supplementary Table S3 for scoring directions). Notably, ID retained cognitive performance comparable to controls despite elevated internalizing symptoms. Personality comparisons revealed that ED was characterized by heightened reward responsiveness, drive, fun seeking (BIS/BAS), and increased urgency (UPPS-P), reflecting emotion-driven impulsivity, while all three biotypes showed impaired perseverance, suggesting shared deficits in self-regulation.

We next examined whether these behavioral distinctions were reflected in the FC score space defined by the first two canonical dimensions (Fig. 3 d). Each biotype occupied a distinct region in this low-dimensional space, consistent with their behavioral signatures: ED scored lowest on the Cognitive–Behavioral Dysfunction dimension, aligning with reduced executive function and elevated externalizing tendencies; ID scored highest on both dimensions, reflecting strong cognition alongside heightened internal distress; and MD showed particularly low scores on the emotion–attention dimension, indicating severe attentional impairments and affective instability. Together, these patterns highlight a coherent alignment between connectivity-derived clustering and symptom-based interpretation.

### Biotype-specific associations with environmental and familial risk factors

Beyond clinical profiles, contextual and familial factors also contributed to symptom variation across biotypes. Relative to controls, all three biotypes showed greater sleep disturbances, with the ED biotype particularly affected by shortened sleep and higher family conflict (Supplementary Fig. S15d). We examined partial correlations between these external factors and behavioral outcomes, controlling for age, sex, parental education, family income, and marital status (Supplementary Fig. S16). Across biotypes, sleep disturbances correlated more strongly with mental health symptoms than in controls. In the ID biotype, higher sleep duration scores (a sleep-disturbance subscale; see Supplementary Table S3 for scoring directions) were more strongly associated with aggression (*r* = 0.19, *p*_FDR_ *<* 0.001) and rule-breaking behavior (*r* = 0.18, *p*_FDR_ *<* 0.001). Family conflict was broadly linked to aggression and rule-breaking, with the strongest associations in ED (*r* = 0.23 to 0.38, *p*_FDR_ *<* 0.001). In the ID biotype, lower parental acceptance related to greater attention problems (*r* = 0.13, *p*_FDR_ *<* 0.001) and withdrawn/depressed symptoms (*r* = 0.13, *p*_FDR_ *<* 0.001). Regarding familial psychiatric history, in ED, parental drug use and antisocial behavior were linked to aggression and rule-breaking (*r* = 0.26 to 0.29, *p*_FDR_ *<* 0.001), with additional effects of parental mania (*r* = 0.14 to 0.20, *p*_FDR_ *<* 0.05). In the MD biotype, parental depression and psychiatric hospitalization correlated with overall symptom burden. Together, these results indicate that biotype-specific symptoms reflect distinct environmental and familial risk patterns.

### Biotypes show distinct patterns of functional network organization

We next assessed whether biotypes exhibited distinct FC patterns in original FC space by comparing each biotype to controls at both the ROI and network levels (Fig. 4), with network-level FC computed as mean connectivity within and between Yeo’s seven functional systems^26^. Group-mean original FC matrices are shown in Supplementary Fig. S17, and pairwise comparisons across biotypes are provided in Supplementary Fig. S18. All three biotypes showed distinct large-scale connectivity patterns. The ED biotype exhibited the most pronounced alterations (Cohen’s *d*), with increased within-network FC in SMN, decreased FC in VN, hyperconnectivity between DAN and LN, and hypoconnectivity between SMN and FPCN, suggesting disrupted motor–executive integration. The ID biotype showed elevated FC within VN and DMN, hyperconnectivity between DMN and LN, and hypoconnectivity between DMN and DAN, consistent with inefficient coordination between emotional salience and attentional control. The MD biotype showed reduced within-network FC in SMN, hyperconnectivity between DMN and DAN/VAN, hypoconnectivity between DMN and LN, and widespread hyperconnectivity between FPCN and other networks. These results indicate that biotype symptom profiles are supported by distinct, atypical network organization.

**Figure 4.**
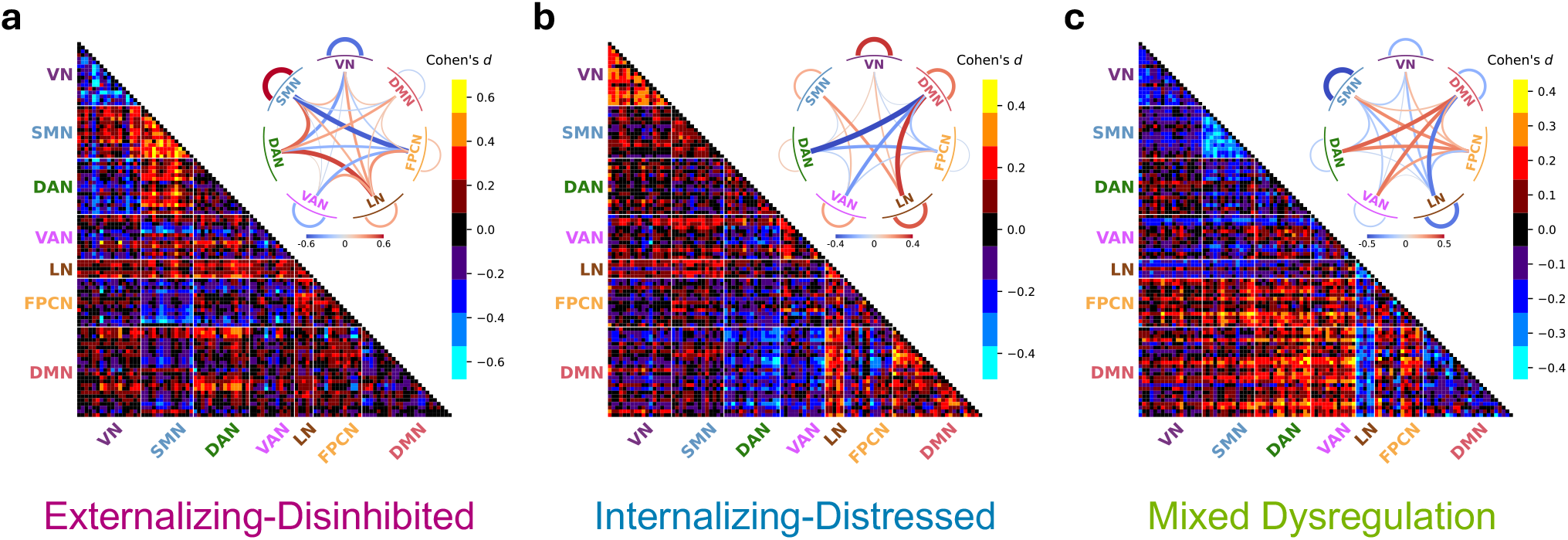
Atypical functional connectivity patterns in each biotype compared with controls. Biotype-specific FC deviations were quantified at both ROI and functional network levels. For each biotype, FC were compared with controls using two-sample *t*-tests, followed by FDR correction (*q <* 0.05). Colorbars indicate effect size (Cohen’s *d*). Warm colors indicate hyperconnectivity (biotype *>* control), and cool colors indicate hypoconnectivity (biotype *<* control). **Bottom left of each panel:** ROI-level FC differences. **Top right of each panel:** Network-level FC differences. Network-level connectivity was computed for each fMRI run by averaging FC within and between functional networks. Edge thickness and opacity encode the effect magnitude. **a**, ED showed increased within-network FC in SMN, reduced FC in VN, hyperconnectivity between DAN and LN, and hypoconnectivity between SMN and FPCN. **b**, ID exhibited elevated within-network FC in VN and DMN, hyperconnectivity between DMN and LN, and hypoconnectivity between DMN and DAN. **c**, MD showed reduced within-network FC in SMN, hyperconnectivity between DMN and DAN/VAN, hypoconnectivity between DMN and LN, and widespread hyperconnectivity from FPCN to other networks.

### Biotypes are characterized by distinct structural brain alterations

Building on the identified FC patterns across biotypes, we examined whether these functional differences were accompanied by structural alterations, using independent structural MRI (sMRI) and diffusion tensor imaging (DTI) measures not used to define biotypes. Analyzing cortical volume, thickness, surface area, subcortical volume, fractional anisotropy, mean diffusivity, longitudinal diffusivity (axial diffusivity), and transverse diffusivity (radial diffusivity) revealed distinct patterns across biotypes (Fig. 5; longitudinal trajectories in Supplementary Fig. S19). Compared with controls, the ED biotype showed significantly reduced cortical volume (*p*_FDR_ *<* 0.001, Cohen’s *d* = 0.26), cortical thickness (*p*_FDR_ *<* 0.01, Cohen’s *d* = 0.13), and cortical surface area (*p*_FDR_ *<* 0.01, Cohen’s *d* = 0.17); the ID biotype exhibited increased cortical volume (*p*_FDR_ *<* 0.05, Cohen’s *d* = 0.10), cortical thickness (*p*_FDR_ *<* 0.001, Cohen’s *d* = 0.17), and subcortical volume (*p*_FDR_ *<* 0.05, Cohen’s *d* = 0.09), along with elevated fractional anisotropy (*p*_FDR_ *<* 0.001, Cohen’s *d* = 0.27), mean diffusivity (*p*_FDR_ *<* 0.01, Cohen’s *d* = 0.12), and longitudinal diffusivity (*p*_FDR_ *<* 0.001, Cohen’s *d* = 0.25); the MD biotype did not show significant global sMRI alterations but exhibited reduced fractional anisotropy (*p*_FDR_ *<* 0.001, Cohen’s *d* = 0.13) and increased transverse diffusivity (*p*_FDR_ *<* 0.01, Cohen’s *d* = 0.12). ROI-level morphology and tract-level DTI results are in Supplementary Figs. S20, S21. These findings indicate neurobiologically distinct biotypes, with FC alterations mirrored by complementary, biotype-specific structural changes, yielding a more comprehensive account of symptom mechanisms.

**Figure 5.**
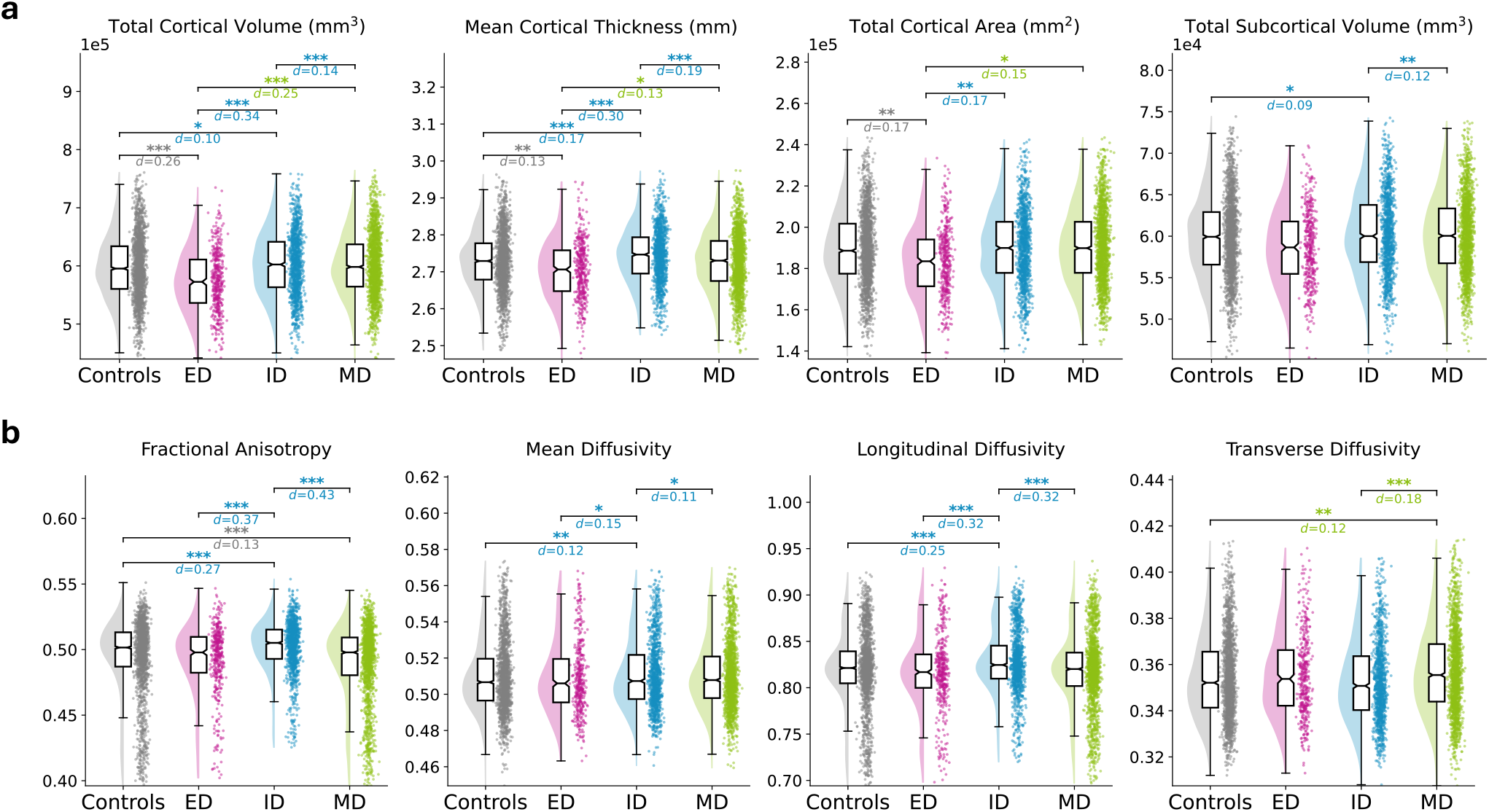
Biotype-specific structural brain differences across sMRI and DTI measures. Pairwise group comparisons for **a** four sMRI metrics (cortical volume, cortical thickness, cortical surface area, and subcortical volume) and **b** four DTI metrics (fractional anisotropy, mean diffusivity, longitudinal diffusivity, and transverse diffusivity). Outliers were excluded within each group using a 3-standard-deviation rule. Linear regression models controlled for age, sex, parental education, family income, marital status, and intracranial volume. Asterisks indicate FDR-corrected significance (*q <* 0.05), Cohen’s *d* effect sizes are shown, and the colors of significance markers and labels denote the group with the higher mean.

### Longitudinal Modeling of Symptom Trajectories

To assess whether symptom trajectories differed across biotypes, we fit linear mixed-effects models (LME) (Method 12) to CBCL subscales from baseline through the fourth follow-up year, using age as the time variable (Supplementary Fig. S22). Models included biotype, covariates (sex, parental education, family income, medication use), and their interactions with age. Significant interactions of biotype and age were obtained for Withdrawn/Depressed (*p*_FDR_ *<* 0.05), Attention Problems (*p*_FDR_ *<* 0.01), Rule-Breaking Behavior (*p*_FDR_ *<* 0.001), Aggressive Behavior (*p*_FDR_ *<* 0.001), Externalizing Problems (*p*_FDR_ *<* 0.01), and Total Problems (*p*_FDR_ *<* 0.01). Group differences were also found in DSM-oriented scales: ADHD (*p*_FDR_ *<* 0.05), Conduct Disorder (*p*_FDR_ *<* 0.001), and Oppositional Defiant Disorder (*p*_FDR_ *<* 0.01). The ED biotype started with the highest symptom levels and showed greater age-related attenuation of symptoms, yet remained highest on externalizing problems across the observed window. The ID biotype showed the flattest trajectories with a slight increase in Withdrawn/Depressed. The MD biotype ended up highest on Total Problems, driven in particular by Attention Problems and Thought Problems.

### External validation of brain–behavior dimensions and biotypes in an independent cohort

To evaluate the generalizability of our findings, we applied the discovery-derived framework (DeCoDE and GMM), without refitting, to an independent cohort of 224 age-matched children diagnosed with ADHD or AXD from the HBN dataset (Method 1). Using identical cognitive and behavioral measures, we obtained significant canonical correlations for both dimensions: Cognitive–Behavioral Dysfunction (*r* = 0.336, *p <* 0.0001, Cohen’s *d* = 0.714) and Emotion–Attention Dysfunction (*r* = 0.179, *p* = 7.39 *×* 10^−3^, Cohen’s *d* = 0.363) (Fig. 6 a). The resulting biotypes closely matched ABCD, with similarity (Method 13) of 0.98 for prevalence (Fig. 3 b vs. Fig. 6 c), 0.87 for clinical profiles (Fig. 3 c vs. Fig. 6 d), 0.86 for FC score distributions for the first two dimensions (Fig. 3 d vs. Fig. 6 e), and 0.63*/*0.57*/*0.72 for FC pattern similarity in ED, ID, and MD biotypes, respectively (Fig. 4 vs. Fig. 6 f). Statistical validation using 1,000 permutations confirmed these correspondences (Supplementary Figs. S23 and S24), all yielding *p*_permutation_ *<* 0.001. These findings demonstrate that brain–behavior dimensions and biotypes identified by DeCoDE are generalizable across independent samples.

**Figure 6.**
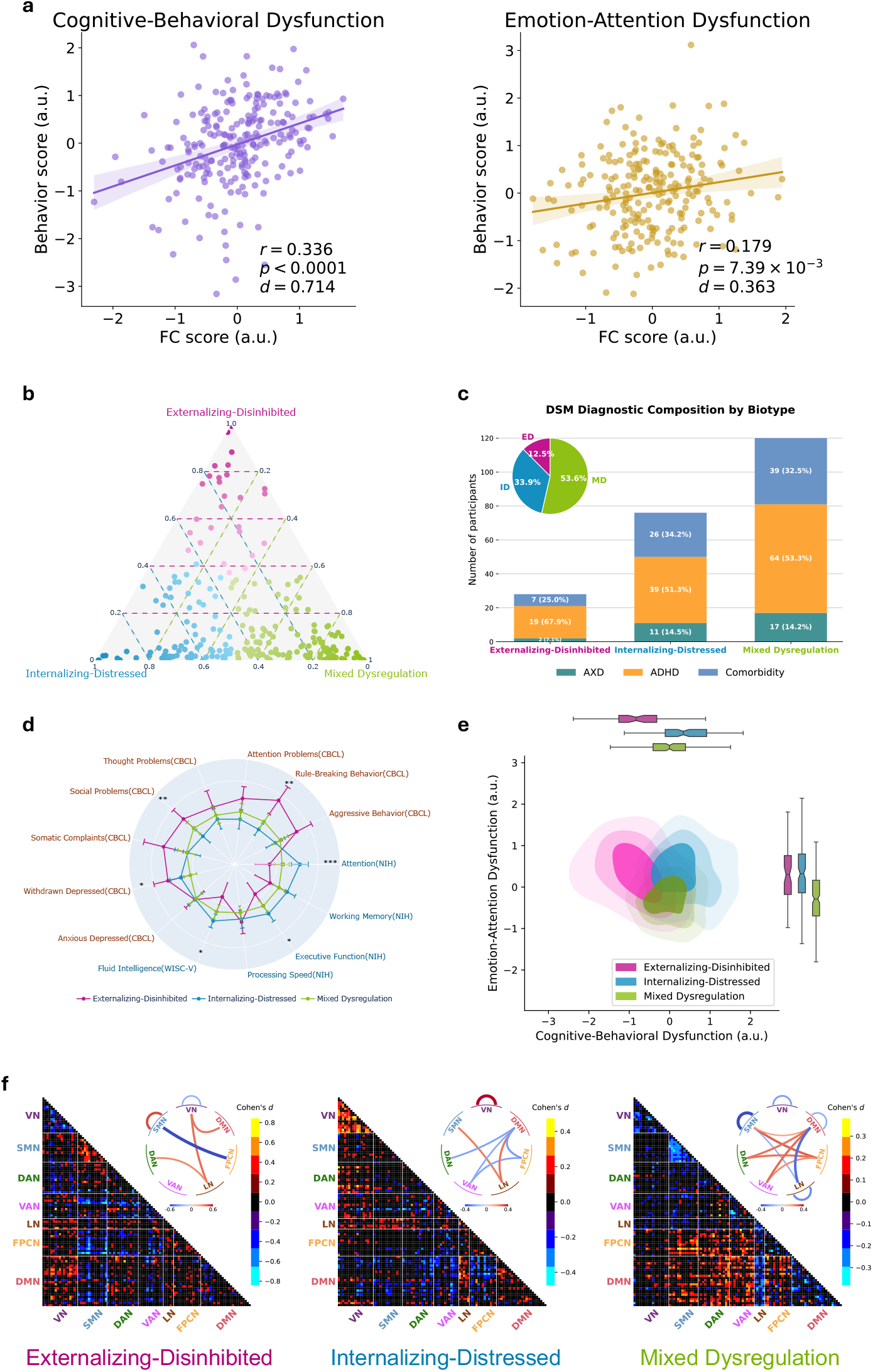
External validation of brain–behavior dimensions and neurofunctional biotypes in the HBN cohort. **a**, Canonical correlations between FC scores and behavior scores for the two identified dimensions: Cognitive–Behavioral Dysfunction and Emotion–Attention Dysfunction. **b**, Posterior membership probabilities for 244 individuals diagnosed with ADHD or AXD, obtained by applying GMM derived from the diagnosed cohort to FC scores. **c**, DSM diagnostic composition of each biotype, displaying the number of individuals diagnosed with ADHD or AXD. **d**, Group clinical profiles across five cognitive and eight mental health scales. Group differences were tested using the Kruskal–Wallis test with FDR correction (*q <* 0.05). **e**, Kernel density estimation plots of biotypes across the two identified brain–behavior dimensions. Contours (25%, 50%, 75%) depict density levels, and marginal boxplots on the top and right summarize score distributions along each dimension. **f**, Functional connectivity deviations from controls at both network and ROI levels, with colors indicating Cohen’s *d* effect sizes.

### Preclinical biotypic expression in late-onset ADHD and AXD

Findings from longitudinal neuroimaging studies indicate that brain alterations precede the onset of psychiatric symptoms, reflecting a vulnerability to these symptoms^27–29^. We therefore applied our framework to a late-onset (LO) group of 693 ABCD children who were free of ADHD or AXD at baseline but received a diagnosis during follow-up (Method 1; demographic and clinical characteristics are summarized in Supplementary Table S4). At baseline, the LO group showed a strong canonical correlation on the Cognitive–Behavioral Dysfunction dimension (*r* = 0.442, *p <* 0.0001, Cohen’s *d* = 0.985) and a weaker yet significant association on the Emotion–Attention Dysfunction dimension (*r* = 0.162, *p <* 0.0001, Cohen’s *d* = 0.328). Biotype assignment was obtained by applying the GMM derived from the diagnosed cohort to baseline FC scores (Supplementary Fig. S25). Despite low baseline symptom severity, LO biotypes had cognitive, personality, contextual, and demographic profiles that closely resembled those of their diagnosed counterparts (Supplementary Fig. S26, Supplementary Table S5), suggesting early, preclinical expression of the same brain–behavior dimensions. No statistically significant sMRI differences were observed after covariate adjustment, although LO-ED showed nominal cortical reductions paralleling those in the diagnosed ED group (Supplementary Fig. S27a). For DTI, LO-ID demonstrated significantly elevated fractional anisotropy (*p*_FDR_ *<* 0.001, Cohen’s *d* = 0.32), mean diffusivity (*p*_FDR_ *<* 0.05, Cohen’s *d* = 0.10), and longitudinal diffusivity (*p*_FDR_ *<* 0.001, Cohen’s *d* = 0.23), consistent with the diagnosed ID group (Supplementary Fig. S27b). Longitudinal CBCL trajectories aligned with their biotype’s defining features: LO-ED showed rising externalizing symptoms (rule-breaking, aggression), LO-ID exhibited a marked increase in internalizing symptoms peaking in the final year, and LO-MD showed persistent attention problems (Supplementary Fig. S28). Developmental trajectories of sMRI and DTI metrics in LO biotypes closely mirrored those of their diagnosed counterparts (cf. Supplementary Fig. S19 vs. Supplementary Fig. S29). Together, these findings suggest that brain–behavior–derived biotypic patterns are detectable before clinical manifestation of ADHD or AXD, highlighting potential for early prevention.

### Comparative validity of conventional DSM classifications and brain-behavior derived biotypes

As shown in Fig. 3 b, each biotype comprised a mix of ADHD, AXD, or comorbid cases, without dominance by any single DSM category, demonstrating substantial cross-diagnostic overlap. To further assess the neurobiological validity of DSM-based groups, we compared their imaging signatures with those of controls across original FC, sMRI, and DTI (Supplementary Fig. S30). Unlike the pronounced functional and structural alterations in biotypes, DSM groups showed limited differentiation from controls across modalities. These findings suggest that transdiagnostic biotypes derived from brain–behavior dimensions provide greater neurobiological specificity than conventional diagnostic labels, offering a more valid framework to parse heterogeneity in childhood psychiatric conditions.

Similarly, the HBN dataset included conventional ADHD subtypes (hyperactive/impulsive, inattentive, and combined), enabling direct comparison with our data-driven biotypes (Supplementary Fig. S31). Consistent with ABCD, HBN biotypes showed greater FC deviations from controls, whereas conventional subtypes showed minimal or no significant differences. This analysis of DSM ADHD subtypes excluded AXD cases, but the biotypes retained clinical profiles closely matching those from the transdiagnostic setting, with significant group differences across multiple cognitive and mental health measures. In contrast, no significant symptom differences were observed among DSM subtypes. These results further underscore the enhanced sensitivity and clinical relevance of data-driven biotyping in capturing biologically and behaviorally meaningful heterogeneity, compared to conventional symptom-based subtype frameworks.

### FC score–derived biotyping outperforms behavior–derived biotyping

All biotyping results above were based on FC scores. For comparison, we assessed behavior-based clustering, which is more accessible and less costly. Clustering the 13 standardized clinical scales yielded no clear optimal cluster number (Supplementary Fig. S32). We then applied biotyping to behavior scores projected via DeCoDE’s behavioral-side canonical loadings. Cluster-number metrics partly agreed with those from FC scores, but cross-metric concordance was weaker, with three clusters no longer the unequivocal choice (Supplementary Fig. S33). If the number of clusters was fixed to three, the distributions of behavior scores on the first two canonical dimensions exhibited large group overlap, and the resulting biotypes showed weak differentiation in atypical FC patterns and less robust external generalization to the HBN cohort, as indicated by marked differences in biotype prevalence (prevalence similarity: 0.62, Supplementary Fig. S34 and S35). These results support our use of FC scores rather than behavior scores for biotyping.

## Discussion

The present study was conducted to clarify diagnostic heterogeneity in developmental psychopathology using a data-driven approach. We identified two robust and reproducible brain–behavior dimensions in youth with ADHD and AXD, capturing variation in cognitive control and emotion–attention interaction. Derived from a large multi-site cohort, these dimensions showed cross-validation consistency and generalized to an independent cohort. These dimensions defined three biotypes that differed systematically in clinical profiles, demographics, and network organization. These findings establish a principled framework for characterizing heterogeneity in cognitive function and internalizing–externalizing symptoms in developmental psychopathology.

A central challenge in modeling transdiagnostic heterogeneity is disentangling disorder-specific brain features from shared variability while capturing nonlinear associations with behavior. Linear methods such as PCA or CCA are limited in this regard, as they assume linear relations and often fail to isolate subtle but clinically relevant variance in high-dimensional connectomes. To overcome these limitations, we developed DeCoDE, a unified framework that extracts latent FC representations maximally informative for distinguishing clinical populations and embeds them in a low-dimensional space preserving meaningful brain–behavior co-variation. By integrating contrastive variational autoencoding with deep multiview embedding, DeCoDE yields individualized connectivity signatures that reflect disorder-specific deviations and align with behavioral profiles. Ablation analyses confirmed the necessity of each component: removing the generative encoder or contrastive objective weakened brain–behavior coupling, particularly for the dimension reflecting subtle internalizing symptoms, and replacing DGCCA with linear GCCA disrupted optimization and cross-modal alignment. Thus, both variational regularization and contrastive supervision are critical for revealing clinically meaningful axes of variation, supporting a shift from categorical diagnoses to biologically grounded, continuous representations of psychopathology.

Clustering directly on clinical scales yielded no clear structure, reflecting the limited ability of symptoms alone to resolve heterogeneity. Although DeCoDE was trained with both brain and behavioral inputs, its design also allows biotyping from behavior scores alone, potentially reducing reliance on fMRI and facilitating clinical use. Yet this behavior-based approach was suboptimal, showing no definitive optimal cluster solutions, weak differentiation of atypical FC patterns, and reduced generalization to the external cohort. These shortcomings likely stem from the subjectivity and low dimensionality of behavioral measures. Thus, neuroimaging-based inference remains the more powerful strategy for identifying biotypes in new patients.

The first brain–behavior dimension, Cognitive–Behavioral Dysfunction, captures a co-occurrence of cognitive ability and behavioral control. Converging evidence links the implicated networks and regions to cognitive ability and externalizing behaviors: the FPCN serves as a flexible control hub for fluid reasoning and working memory, with stronger activity tracking better spatial working-memory performance and higher fluid intelligence^30–32^; the DAN subserves top–down attentional selection and goal maintenance^33,34^; the precuneus contributes to sustained attention and cognitive control^35–37^; and the middle cingulate supports goal-directed action sequencing and performance monitoring central to behavioral regulation^38^. Together, the evidence positions cognitive ability and rule-breaking behavior as opposing poles of a shared neurobiological axis, where disruption may compromise top-down regulation and increase vulnerability to impulsive actions.

The second brain–behavior dimension, Emotion–Attention Dysfunction, delineates a transdiagnostic axis of affective and attentional symptoms. The underpinning neural circuits have been linked to internalizing symptoms or attentional control: within the LN, circuitry supporting negative affect and stress reactivity is centrally implicated in internalizing psychopathology^39,40^; the VAN supports reorienting and is linked to emotion-driven attentional capture^41,42^; altered between-network connectivity of the DAN is associated with attentional difficulties in youth^43,44^; and the insula supports interoception and affect regulation and shows alterations linked to internalizing symptoms in youth^45–47^. Moreover, deliberate attentional deployment in emotional contexts can mitigate negative affect^48,49^. These observations suggest that Emotion–Attention Dysfunction reflects a shared neurobiological axis in which attentional control and emotional dysregulation are reciprocally linked, highlighting implicated circuits as candidate targets for mechanistic studies and interventions in youth.

Embedding individuals within this two-dimensional space revealed three biotypes with distinct behavioral and neurobiological signatures. The ED biotype, although least prevalent (12.5%), exhibited the most severe clinical impairments, marked by externalizing symptoms and reduced cognitive performance. Positioned at the low end of the Cognitive–Behavioral Dysfunction axis (Fig. 3 d), ED showed disrupted SMN organization, with heightened within-network connectivity and reduced coupling with the FPCN, consistent with impaired top-down modulation of sensorimotor processes^50–54^. VN hypoconnectivity further aligned with abnormal visual encoding of reward and threat^16,55^. Structurally, ED displayed widespread reductions in cortical volume, thickness, and surface area (Fig. 5), resembling prior ADHD subgroups characterized by gray matter reductions and impaired cognitive control^56–58^. These reductions co-occurred with accelerated pubertal development and align with evidence that lower socioeconomic status is associated with faster cortical thinning and maturation trajectories, consistent with compensatory acceleration accounts^59^.

In contrast, the ID biotype presented with prominent internalizing symptoms despite preserved cognitive performance, positioning at the high end of the Cognitive–Behavioral Dysfunction dimension^55,60^. FC abnormalities centered on the DMN, including hyperconnectivity within the network, stronger coupling with the limbic system, and reduced integration with the DAN. These alterations are consistent with excessive self-referential focus^61,62^, rumination, and impaired coordination with attention systems^43^, and mood-related symptoms linked to DMN–LN coupling^63^. Structurally, ID exhibited widespread increases in cortical thickness and prefrontal volume, indicating atypical cortical maturation^64,65^, a pattern that persisted longitudinally despite normative declines. White matter abnormalities, including elevated fractional anisotropy and longitudinal diffusivity, further suggested inefficient pruning^66^, with significant increases in the cingulum and uncinate fasciculus aligning with DMN–limbic coupling implicated in internalizing symptoms^67^. Together, these multimodal alterations delineate a distinct ID-specific trajectory that increases vulnerability to internalizing psychopathology.

The MD biotype, the largest subgroup (52%), was defined by pronounced attentional deficits and body-focused repetitive behaviors (BFRBs)^68^. It was positioned at the low end of the Emotion–Attention Dysfunction axis and showed FC deviations nearly opposite to those of the ID biotype, consistent with behavioral dissociation across the two dimensions. Their connectomic profile was marked by widespread external dysregulation of both DMN and FPCN, with increased connectivity to attention-related networks (DAN, VAN) and SMN. DMN also showed reduced integration with LN. This pattern may reflect instability between internal and external attention systems, impairing response inhibition and increasing susceptibility to environmental distraction^69,70^. Additionally, SMN hypoconnectivity may contribute to motor disinhibition and attentional lapses, consistent with prior reports linking motor overflow and reduced nodal efficiency to BFRBs and inattention^71,72^. Unlike ED and ID, the MD group did not exhibit notable structural MRI alterations but showed reduced fractional anisotropy and elevated transverse diffusivity, indicative of a myelination lag or delayed axonal development during adolescence^73,74^.

Despite overlapping diagnoses, the three biotypes followed dissociable neurodevelopmental trajectories. The ED biotype showed the most pronounced functional and structural deviations and was disproportionately linked to contextual adversity, including low socioeconomic status, non-intact family structure, family conflict, parental psychopathology, and health-related risks such as elevated BMI and sleep disturbance. Although ED youth exhibited greater age-related symptom remission, high rates of aggressive and destructive behaviors (e.g., firesetting, stealing, self-injury) underscore the need for early detection and targeted monitoring. The ID biotype resembled controls in cognitive performance and demographics, potentially obscuring early recognition, yet showed limited remission and rising depressive symptoms over time, suggesting heightened risk for adolescent depression. Elevated fractional anisotropy and longitudinal diffusivity in both diagnosed and late-onset ID cases delineate a distinctive white matter signature that may serve as an early biomarker, highlighting the need for adjunctive, mechanism-targeted interventions rather than standard psychotherapy alone. The MD biotype, the most prevalent, exhibited reduced fractional anisotropy, increased transverse diffusivity, and delayed pubertal maturation, pointing to a broader developmental lag. Together, these findings reveal distinct neurofunctional pathways underlying shared clinical features and support biotype-informed psychotherapy approaches, such as tailored emotion-regulation and impulse-control modules within cognitive behavioral therapy^75^, and attention and executive skills training^76^.

Our findings generalized robustly to the independent HBN cohort. Despite milder clinical profiles in ABCD under stricter diagnostic criteria^77^, the model performed well on HBN without retraining or fine-tuning. Both dimensions showed significant brain–behavior correlations that matched ABCD, and FC-derived biotypes in HBN closely resembled those in ABCD across functional and clinical profiles, suggesting that DGCCA captures transferable brain–behavior associations. Generalizability was also supported by the training scale (over 3,500 participants and 12,000 imaging runs), ensuring statistical power and stable canonical components for out-of-sample transfer^78,79^. Applying the model to the LO group in ABCD, despite the absence of baseline diagnoses, also revealed early biotypic patterns. LO youth showed significant brain–behavior associations and longitudinal symptom trajectories aligned with their biotype’s defining features. Notably, despite being derived solely from rs-fMRI FC, LO biotypes closely mirrored diagnosed biotypes across cognitive, personality, contextual, demographic, and structural measures. These observations highlight the richness of information embedded in rs-fMRI FC and the robustness of our framework.

Our results further highlight the limitations of DSM-based categories. Both atypical FC patterns within diagnostic labels and traditional ADHD subtypes (inattentive, hyperactive/impulsive, combined) showed weak differentiation from controls, underscoring the misalignment between symptom-based categories and underlying neurophysiological variation. In contrast, brain-derived biotypes demonstrated clearer functional and structural deviations, supporting their value as mechanistically informed tools for parsing heterogeneity in childhood psychiatric conditions.

Several limitations warrant consideration. First, although biotypic differences may reflect both genetic and environmental influences, the absence of genomic analyses in the present study limited our ability to disentangle these factors or their potential interactions. Incorporating polygenic risk scores could elucidate etiological pathways^80^, and how such genetic endowments may interact with environmental factors to produce phenotypic variation. Second, richer behavioral measures, when available, may refine or uncover additional canonical dimensions. Third, our analyses focused on baseline rs-fMRI, leaving developmental changes in connectivity and their relation to symptom progression unexplored. Finally, although DGCCA is inherently multimodal, the present implementation used only two modalities. Incorporating genomic data, structural MRI, or task-based fMRI in future research may yield deeper insights into the neurobiological organization of transdiagnostic subgroups.

In summary, we identified two robust, generalizable brain–behavior dimensions and three transdiagnostic biotypes in youth with ADHD/AXD. These dimensions provide a mechanistic account of how cognitive control, attention, and emotion regulation jointly shape symptom heterogeneity. By integrating transdiagnostic behaviors with neuroimaging features, our framework delineates distinct neurophysiological pathways underlying shared clinical presentations. This biotype-based stratification advances biologically informed classification and provides a basis for targeted, mechanism-based interventions, moving toward precision psychiatry guided by objective brain markers. Establishing these biotypes’ stability and clinical utility in prospective designs, and testing mechanism-matched interventions for differential benefit, will enable translation of biotype-based stratification into care pathways.

## Data availability

Data used in this study are available from established repositories under their standard access procedures. The Adolescent Brain Cognitive Development (ABCD) Study can be accessed via https://abcdstudy.org/. The Healthy Brain Network (HBN) dataset is available through the Child Mind Institute Biobank at https://fcon_1000.projects.nitrc.org/indi/cmi_healthy_brain_network.

## Code availability

The DeCoDE framework was implemented in Python (v3.12.11) and PyTorch (v2.5.1). Upon acceptance of the manuscript, the code will be released in a public GitHub repository. Functional connectivity was computed with Nilearn (v0.11.1). Gaussian mixture modeling (GMM), orthogonal matching pursuit (OMP), and linear regression models were performed with scikit-learn (v1.7.2). Statistical analyses used SciPy (v1.15.2), statsmodels (v0.14.4), and R (v4.5.0).

## Methods

### Method 1: Participants

This study used data from the Adolescent Brain Cognitive Development (ABCD)^81^ and the Healthy Brain Network (HBN) Release 11^82^ datasets. For ABCD, all tabulated measures were obtained from Release 5.1, while demographic information and resting-state fMRI data were obtained from the ABCD BIDS Community Collection (ABCC; Collection 3165)^83^. The ABCD study enrolled over 11,000 children aged 9–10 years from 21 sites across the United States, with extensive neuroimaging, cognitive, behavioral, developmental, and psychiatric assessments. Study protocols were approved by the Institutional Review Board (IRB) at each site^84^. Parents or guardians provided written informed consent, and children assented. After excluding participants with incomplete rs-fMRI, missing diagnostic or behavioral data at baseline, or failed MRI preprocessing and quality control, 6,148 participants remained. Of these, 3,508 youth diagnosed with ADHD or AXD at baseline constituted the discovery cohort, 1,947 children with no diagnosis throughout the study served as controls, and 693 children without a baseline diagnosis but who developed ADHD or AXD during follow-up formed the LO group. AXD included generalized anxiety disorder, social anxiety disorder (social phobia), specific phobia, separation anxiety disorder, panic disorder, agoraphobia, and selective mutism. Data filtering steps are shown in Supplementary Fig. S36, and sociodemographic and clinical characteristics of the groups are summarized in Supplementary Table S4.

The HBN dataset is a large-scale, community-based initiative designed to advance the understanding of neurodevelopmental disorders. It includes over 4,000 participants aged 5–21 years, primarily recruited from the New York area, who were scanned at four sites: Staten Island Flagship Research Center, Rutgers University Brain Imaging Center, CitiGroup Cornell Brain Imaging Center, and CUNY Advanced Science Research Center. The dataset provides multimodal neuroimaging, behavioral assessments, cognitive tests, and extensive clinical evaluations. Ethical approval for the HBN study was obtained from the Chesapeake Institutional Review Board, and participants provided written informed consent or assent, as appropriate, before data collection. Using the same data filtering procedure as the ABCD study, 224 age-matched participants diagnosed with ADHD or AXD were included as the external validation cohort. AXD included generalized anxiety disorder, social anxiety disorder (social phobia), specific phobia, separation anxiety disorder, and other specified anxiety disorder. Detailed data filtering procedures are provided in Supplementary Fig. S36.

### Method 2: Cognitive and behavioral assessment

Thirteen clinical scales commonly assessed in ABCD and HBN were used for model training in this study, including five cognitive scales and eight mental health scales. The cognitive battery consists of four tasks from the NIH Toolbox^21^: the Flanker Test, List Sort Working Memory Task, Dimensional Change Card Sort Task, and Pattern Comparison Processing Speed Task, which measure attention, working memory, executive function, and processing speed, respectively. Additionally, one matrix reasoning task from the Wechsler Intelligence Scale for Children (WISC-V)^22^ evaluates fluid intelligence. The eight mental health scales are all from the parent-reported Child Behavior Checklist (CBCL)^23^. The correlations among the thirteen clinical scales are presented in Supplementary Fig. S37. The subscales encompass the following domains: Anxious/Depressed, which evaluates symptoms associated with anxiety and depression; Withdrawn/Depressed, focusing on social withdrawal and depressive tendencies; Somatic Complaints, addressing physical symptoms lacking a clear medical explanation and often tied to emotional distress; Social Problems, assessing challenges in social interactions, including difficulties with peers; Thought Problems, identifying atypical thoughts or behaviors such as peculiar ideas or obsessions; Attention Problems, measuring inattention, impulsivity, and hyperactivity; Rule-Breaking Behavior, evaluating actions that violate societal norms or rules; and Aggressive Behavior, examining confrontational or hostile actions. To preserve the full scope of brain-behavior relationships and avoid potential information loss from overadjustment, we used uncorrected behavioral data in our analyses. Specifically, we employed uncorrected standard scores from the NIH Toolbox and unadjusted raw scores from CBCL and WISC during model training.

### Method 3: fMRI acquisition and preprocessing

MRI data from the ABCD study were collected across 21 sites in the United States using Siemens Prisma, General Electric 750, and Philips scanners, harmonized to ensure compatibility. Each participant completed 2–4 runs of rs-fMRI with a multiband echo-planar imaging sequence (2.4 mm isotropic voxels, TR = 800 ms, TE = 30 ms, flip angle = 52°, multiband acceleration = 6, 60 slices), yielding approximately 20 minutes of data per participant while viewing a fixation crosshair^81^. Only baseline rs-fMRI data were analyzed in this study. ABCD imaging data were obtained from the ABCD-BIDS Community Collection (ABCC; Collection 3165)^83^, which provides minimally preprocessed data using the DCAN Labs HCP-style surface-based pipeline^85^. This workflow includes distortion correction, motion correction, registration to structural images, and projection to cortical surfaces, yielding CIFTI time series harmonized across scanner platforms.

For HBN, rs-fMRI data were acquired on 3T scanners with site-specific multiband protocols (e.g., Staten Island: 2.5 mm isotropic voxels, TR = 1450 ms, TE = 40 ms; other sites: 2.4 mm isotropic voxels, TR = 800 ms, TE = 30 ms). Preprocessing was performed locally using fMRIPrep^86^, including intensity nonuniformity correction, skull stripping, tissue segmentation, susceptibility distortion correction, boundary-based co-registration to T1-weighted images, and spatial normalization to MNI152NLin2009cAsym space. Motion artifacts were further mitigated with ICA-AROMA, and data were spatially smoothed with a 6 mm FWHM Gaussian kernel. Runs with mean framewise displacement greater than 0.5 mm or fewer than 300 volumes were excluded in both datasets.

### Method 4: Functional connectivity calculation

Regional time series were calculated by averaging the preprocessed voxel-level BOLD signals across 100 regions of interest (ROIs) defined by the Schaefer parcellation^87^ (Supplementary Table S6). FC was then computed as the Pearson correlation coefficient between the time series of each ROI pair. The resulting connectivity values underwent Fisher’s *r*-to-*z* transformation to improve normality^88^, and were subsequently demeaned within each rs-fMRI run to remove global connectivity offsets. Considering the consistency of fMRI acquisition across ABCD’s multi-site design and the harmonization already performed, ComBat harmonization^89^ was applied to address variations in scanner settings within the HBN dataset. This procedure was conducted across the four collection sites of HBN, using the entire ABCD cohort as a reference site and incorporating sex and age as covariates. It is important to note that the ABCD cohort served solely as a reference for aligning HBN data; their original FC features were preserved for model training and subsequent statistical analyses.

### Method 5: Contrastive variational autoencoder

We applied a cVAE to extract disorder-specific latent features from functional connectivity data, enabling the model to disentangle clinically relevant neural variation from background population variability. Unlike standard variational autoencoders (VAEs)^90^, which learn a unified latent space, the cVAE introduces two distinct sets of latent variables: **s**, which encode salient, disorder-specific signals present only in the target data, and **z**, which capture shared variability present in both target and background datasets. This separation enhances interpretability and facilitates the identification of transdiagnostic brain–behavior associations.

The cVAE is trained on both target data **X** and background data **B**. The model architecture consists of two independent encoders, a salient encoder 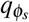 and a background encoder 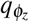, along with a single shared decoder *p*_*θ*_. For target data, both salient and shared background features contribute to reconstruction. The corresponding evidence lower bound (ELBO) for the target dataset is defined as:

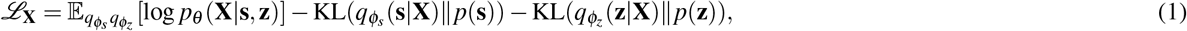

For background data, only the background encoder is used, and the salient latent variable is set to zero during decoding. Its ELBO is given by:

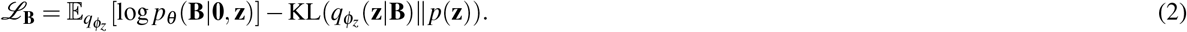

To promote independence between the salient and shared latent spaces, the model includes a total correlation loss, as introduced by Abid et al.^17^:

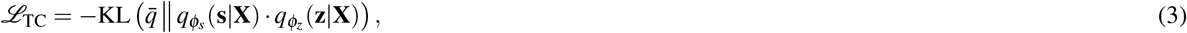

where 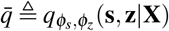 denotes the joint posterior over the latent variables. This term penalizes redundancy and encourages the two encoders to learn disentangled representations.

The final training objective combines the ELBOs for both datasets with the total correlation regularization:

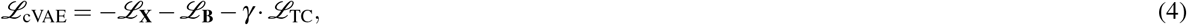

where *γ* is a weighting parameter controlling the strength of the regularization.

### Method 6: Deep generalized canonical correlation analysis

CCA is a classical statistical method that identifies maximally correlated projections between two datasets^91^. Generalized CCA (GCCA) extends this approach to multiple data views by finding a shared low-dimensional representation that minimizes the discrepancy between views across all modalities^92^. However, traditional GCCA is limited to linear mappings, which restricts its ability to capture nonlinear dependencies in high-dimensional data. DGCCA addresses these limitations by incorporating deep neural networks to model view-specific nonlinear transformations. Given *J* data modalities (or views), denoted as 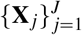, where each 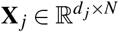 is a data matrix of *N* samples and *d* _*j*_ features, DGCCA learns nonlinear mappings *f* _*j*_(*·*), parameterized by neural networks, that embed each view into a latent space. Let 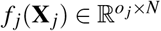 denote the transformed output of the *j*-th view. DGCCA then learns a shared latent representation **G** ∈R^*r×N*^ and linear projection matrices 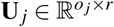(i.e., loading matrix) such that the following objective is minimized:

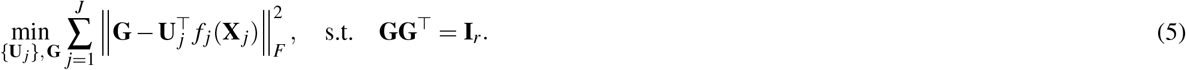

Here, ∥ · ∥_*F*_ denotes the Frobenius norm. The orthogonality constraint **GG**^*T*^ = **I**_*r*_ ensures that the learned shared representation has uncorrelated and unit-variance components across the *r*-dimensional latent space. In practice, this corresponds to minimizing the following DGCCA loss function with respect to all neural network parameters and projection matrices:

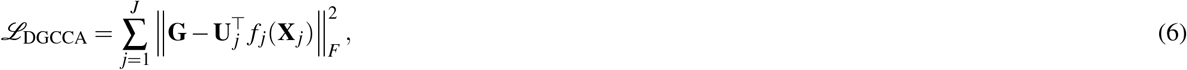

where **G** is updated at each iteration to satisfy the orthogonality constraint. DGCCA is trained end-to-end using backpropagation and facilitates the learning of shared representations that are robust to noise, flexible to modality-specific variation, and effective in high-dimensional settings.

### Method 7: Joint optimization of DeCoDE

To jointly eliminate irrelevant variation and maximize cross-view correlation, the cVAE and DGCCA are integrated into a unified end-to-end architecture. The cVAE acts as the contrastive learning module, isolating clinically relevant latent FC by contrasting patients with controls to attenuate normative variation; DGCCA is used as the brain–behavior linking module, projecting latent FC features and multivariate behavioral measures into a shared low-dimensional space. The cVAE’s salient encoder also serves as the feature extractor for DGCCA. This allows the salient latent representation to be optimized to maximize cross-view correlation with the auxiliary view within each training batch. The total loss combines the cVAE objectives with the DGCCA alignment term:

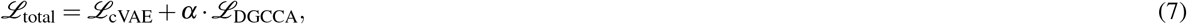

where *α* controls the contribution of the multiview correlation objective.

### Method 8: Implementation details of DeCoDE

The joint deep learning models were trained on the discovery ABCD dataset. We defined the target dataset **X** as resting-state FC vectors from 3,508 youth diagnosed with ADHD or AXD, and the background dataset **B** as 1,947 controls. Each FC vector contained 4,950 edges derived from the lower triangle of the pairwise correlation matrix among 100 brain regions. In addition to the FC vectors from patients, a second view comprising thirteen clinical scales, paired with each patient’s FC vector, was incorporated to capture cognitive and mental health profiles. Given the low dimensionality of this behavioral view, it was directly used as input to the DGCCA objective without an additional neural network.

Both encoders in the cVAE were implemented as multilayer perceptrons with architecture [4950, 256, 32], and the shared decoder adopted the reverse structure [64, 256, 4950]. The model was trained end-to-end using the Adam optimizer. The 3,508 participants were randomly partitioned into ten folds for cross-validation to determine training epochs and hyperparameters. To ensure dimensional correspondence across folds and datasets, we defined a *reference model*, trained on the entire set of 3,508 participants using the selected hyperparameters; components from each validation fold were reordered to this reference via the Hungarian algorithm^93^, ensuring a consistent dimension order for downstream analyses. All scans from the same participant were assigned either to the training or the validation set. Model selection and early stopping were guided by the average correlation across validation folds, aiming for stable convergence and maximal cross-view correlation (Supplementary Fig. S38). The final learning rate was 5 *×* 10^−6^, and the DGCCA and total correlation weights were fixed at *α* = 17 and *γ* = 5, respectively.

After tuning, the reference model was used to generate latent representations for downstream analyses and was subsequently applied to the independent HBN dataset and the ABCD late-onset group. For the final evaluation, *r*_cv_ was computed as the Pearson correlation between FC-view and behavioral-view canonical scores obtained in cross-validation, where within each fold training-set loadings were applied to the held-out set, FC scores from multiple runs were averaged within participant, and per-participant score pairs were concatenated across folds. The number of identified brain–behavior dimensions was determined based on cross-validated model performance. Although the third dimension reached nominal significance given the large sample size, its cross-validated correlation was low (*r*_cv_ = 0.070; Cohen’s *d* = 0.141), and its canonical loadings showed substantially reduced cross-validation agreement across folds compared with the first two dimensions (Supplementary Fig. S3). Therefore, subsequent analyses focused on the first two robust dimensions.

### Method 9: Interpretable loading approximation

Since the cVAE employs nonlinear deep encoders, the DGCCA canonical weight matrix **U** implements a linear projection from the latent representation **Z** to the canonical scores **S** = **ZU** but cannot be directly expressed in terms of the original FC feature space **X**. To enable interpretable analysis of brain–behavior dimensions in the original FC feature space, we therefore learn a sparse loading matrix **W** such that **XW** ≈**S**, yielding edge-level loadings for interpretation.

We estimate **W** with OMP, solving

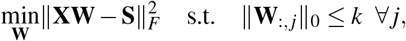

where *k* controls the number of nonzero FC features per dimension. This yields sparse loadings over FC edges and supports circuit-level interpretation.

The procedure approximately preserves the dimensional separation learned by DGCCA. The DGCCA objective enforces **GG**^*T*^ = **I**_*r*_ on the shared representation **G**, which implies orthonormal canonical scores **S** = **G**^*T*^ with **S**^*T*^**S** = **I**_*r*_. Using these columns as independent OMP targets promotes decorrelated sparse directions in **W**.

OMP also provides nested solutions: feature supports selected at lower sparsity are retained as *k* increases, enabling straightforward feature ranking and improving stability across sparsity levels. In contrast, Lasso-type decoders can add or drop features non-monotonically as regularization changes. Although **W** is an approximation rather than an exact inversion of the nonlinear encoder, it provides a practical and stable route to interpretability in large-scale analyses, yielding sparse loading patterns suitable for downstream biological inspection.

### Method 10: Network-level FC importance

Let *R* denote the number of regions of interest and *E* = {(*i, j*) : 1 ≤ *i < j* ≤ *R*} the set of undirected ROI pairs with |*E*| = *R*(*R* − 1)*/*2. Let **w**_OMP_ ∈ ℝ^|*E*|^ be the OMP-derived connectivity-loading vector, and define nonnegative edge weights *w*_*ij*_ = |**w**_OMP_(*i, j*)| for (*i,j) ∈ E*. Let 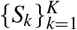 be the *K* functional networks with sizes *n*_*k*_ = |*S*_*k*_|. To obtain size-invariant summaries, we define three per-edge-averaged measures: within-network importance,

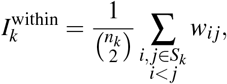

pairwise inter-network importance,

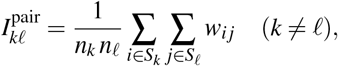

and between-network (aggregate inter-network) importance,

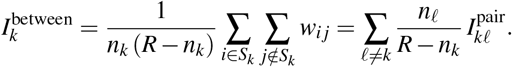

For visualization, quantities displayed together were rescaled to [0, 1] within each comparison set to place values on a common scale; this display normalization does not alter the definitions above.

### Method 11: Biotype identification via clustering

To identify stable and biologically meaningful biotypes, we extracted the posterior mean of salient latent features from the reference cVAE model. These latent vectors were projected onto DGCCA-derived FC loadings to obtain FC scores, which served as input to a GMM for unsupervised clustering. The GMM was fit on FC scores from 3,508 diagnosed youth in ABCD. The optimal number of clusters was determined using the Bayesian Information Criterion (BIC), Akaike Information Criterion (AIC), and clustering stability via repeated subsampling (Supplementary Fig. S12). The fitted model can then be directly applied to assign biotype labels in clinically similar populations.

### Method 12: Longitudinal mixed-effects modeling

Longitudinal effects were tested with linear mixed-effects models estimated by restricted maximum likelihood (REML) in R (version 4.5.0), with age at each visit modeled as the time variable. All covariates were defined at baseline and treated as time-invariant. A random intercept for each participant accounted for repeated measurements; random slopes were excluded to ensure convergence and interpretability. We first fit a full model with biotype and all covariates, each interacting with age:

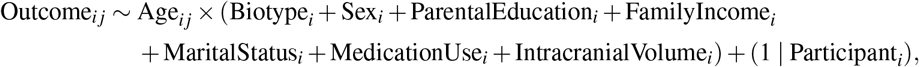

where *i* indexes participants and *j* indexes timepoints. Fixed-effect significance was assessed with Type III Wald *χ*^2^ tests; *p*-values were FDR-corrected for each target effect across outcomes. Only covariates that remained significant after correction were retained in the final model.

### Method 13: Similarity metrics for biotype validation

To assess the generalizability and external validity of the identified biotypes, we computed four complementary similarity metrics capturing distinct feature domains: (i) prevalence similarity, (ii) clinical profile similarity, (iii) FC score distributional similarity, and (iv) FC pattern similarity.

#### (i) Prevalence similarity

Similarity was quantified as 1 − JSD, where the Jensen–Shannon divergence (JSD) was computed from the biotype proportion vectors using base-2 logarithms. This yields values in [0, 1], with higher scores indicating more similar biotype distributions.

#### (ii) Clinical profile similarity

For each dataset, biotype-specific mean scores were computed across the five cognitive and eight mental health scales used in model training. Cognitive measures were approximately normally distributed and underwent outlier removal using a 3-standard-deviation threshold, whereas mental health scales were non-normally distributed and processed using the Median Absolute Deviation method with a threshold of 4. All scales were min–max normalized within scale, yielding a scale-by-biotype matrix. Scale-wise Pearson correlations across biotypes between datasets were then computed and averaged across scales to obtain the final similarity score (range [− 1, 1]).

#### (iii) FC score distributional similarity

A kernel density estimation (KDE)-based JSD metric was used to compare the distribution of FC scores between datasets. The first two CCA-derived FC score dimensions were extracted and stratified by biotype. For each biotype, a two-dimensional Gaussian KDE was fitted, evaluated over a shared grid, and normalized to a probability map. The base-2 JSD between maps was computed and transformed to similarity as 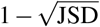. Biotype-level scores were aggregated via weighted averaging, with weights proportional to the mean biotype-specific sample size across datasets (range [0, 1]).

#### (iv) FC pattern similarity

For each biotype, a vector of *t*-statistics was obtained by comparing FC with the control group using Welch’s two-sample *t*-tests across features, setting non-significant values (*p*_FDR_ ≥ 0.05) to zero. Pearson correlations between corresponding *t*-statistic vectors across datasets yielded one similarity value per biotype (range [−1, 1]).

## Acknowledgements

This study was funded by grants from the US National Institutes of Health (R01MH129694, R21AG080425, and R21MH130956) and the Alzheimer’s Association grant (AARG-22-972541).

## Author contributions

Y.J. conceptualized and designed the study, developed the code, performed data analyses, interpreted results, and drafted and revised the manuscript. X.T. contributed to conceptualization, advised on result interpretation, and revised the manuscript. G.A.F., I.H.G., K.M.P., and T.D.S. contributed to conceptualization and revised the manuscript. J.J. and Y.Z. conceptualized and designed the study, oversaw data analyses and interpretation, and revised the manuscript.

