## Supplementary Material for "Deep Learning of Brain-Behavior Dimensions Identifies Transdiagnostic Biotypes in Youth with ADHD and Anxiety Disorders"

**Figure S1. Correlation of FC and behavior scores between the Cognitive–Behavioral Dysfunction and Emotion–Attention Dysfunction dimensions.** Analyses were performed on 3,508 participants diagnosed with ADHD or AXD in the ABCD dataset (10-fold cross-validated). No significant correlations were observed, confirming the independence of the two dimensions.

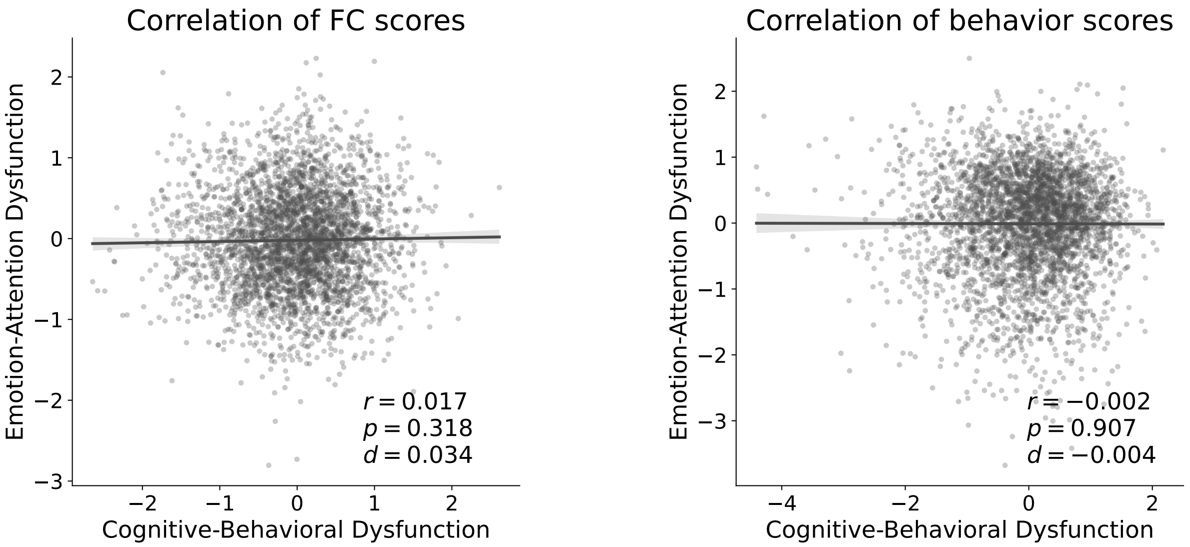

**Figure S2. Sex-stratified correlations between FC scores and behavior scores for the Cognitive–Behavioral Dysfunction and Emotion–Attention Dysfunction dimensions.** **a**, Participants diagnosed with ADHD or AXD in ABCD (N=3,508; 10-fold cross-validated); **b**, Age-matched HBN participants with ADHD or AXD (N=224); **c**, ABCD late-onset participants (N=693; no baseline diagnosis; later diagnosed with ADHD or AXD). Fisher Z tests detected no significant sex differences in correlation strength within datasets.

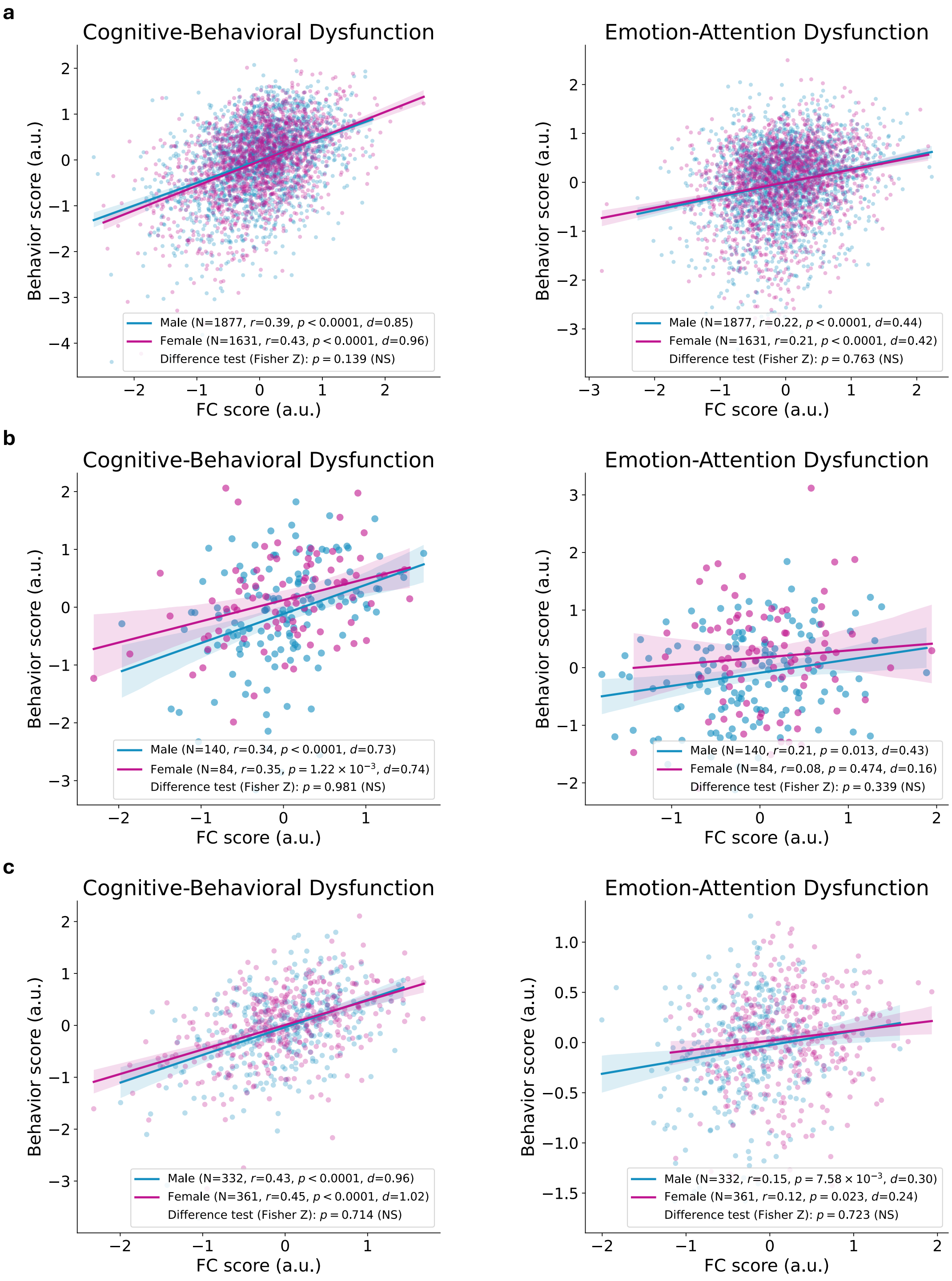

**Figure S3. Cross-validation agreement of canonical components across folds.** Distributions of Pearson correlation coefficients between reference and 10-fold cross-validation CCA loadings. Components in each fold were reordered based on their correlations with the reference behavior loadings. **a**, Correlation coefficients of FC loadings across ten folds. **b**, Correlation coefficients of behavior loadings across ten folds.

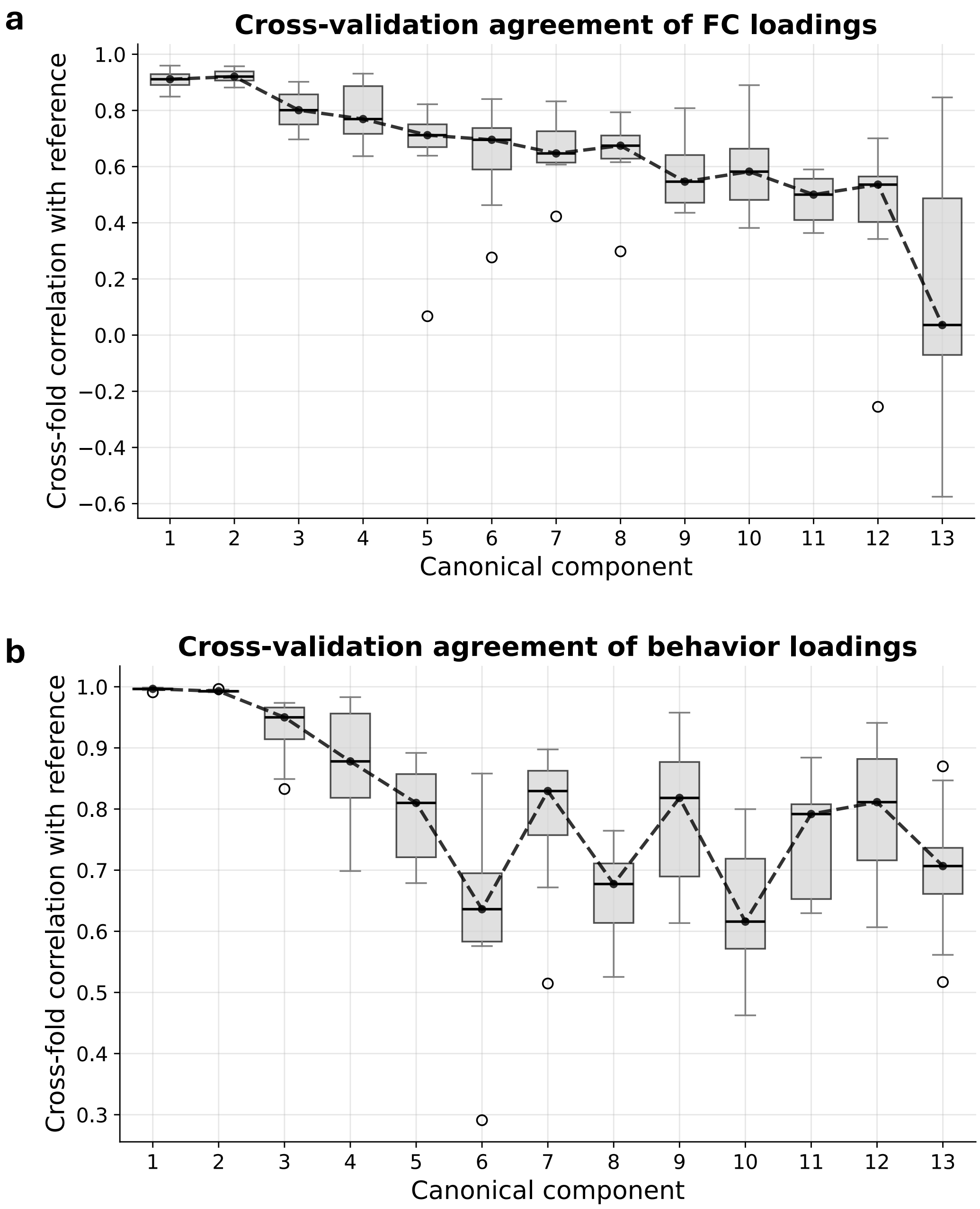

**Figure S4. Ablation analysis of the cVAE + GCCA model.** cVAE and GCCA trained separately without unified optimization. **a**, Cross-validated correlation curves for each dimension during training. Lines show fold-averaged values; shaded areas represent standard deviation across folds. **b**, Intraclass correlation (ICC) of canonical loadings across 10-fold cross-validation. **c**, Behavior loadings of cognitive measures and CBCL subscales on the first (left) and the second (right) canonical dimensions. Error bars indicate standard deviation across folds. **d**, Correlations between FC and behavior scores for the first two canonical dimensions in the ABCD cohort using 10-fold cross-validation. **e**, Correlations between FC and behavior scores for the two identified canonical dimensions in the HBN cohort.

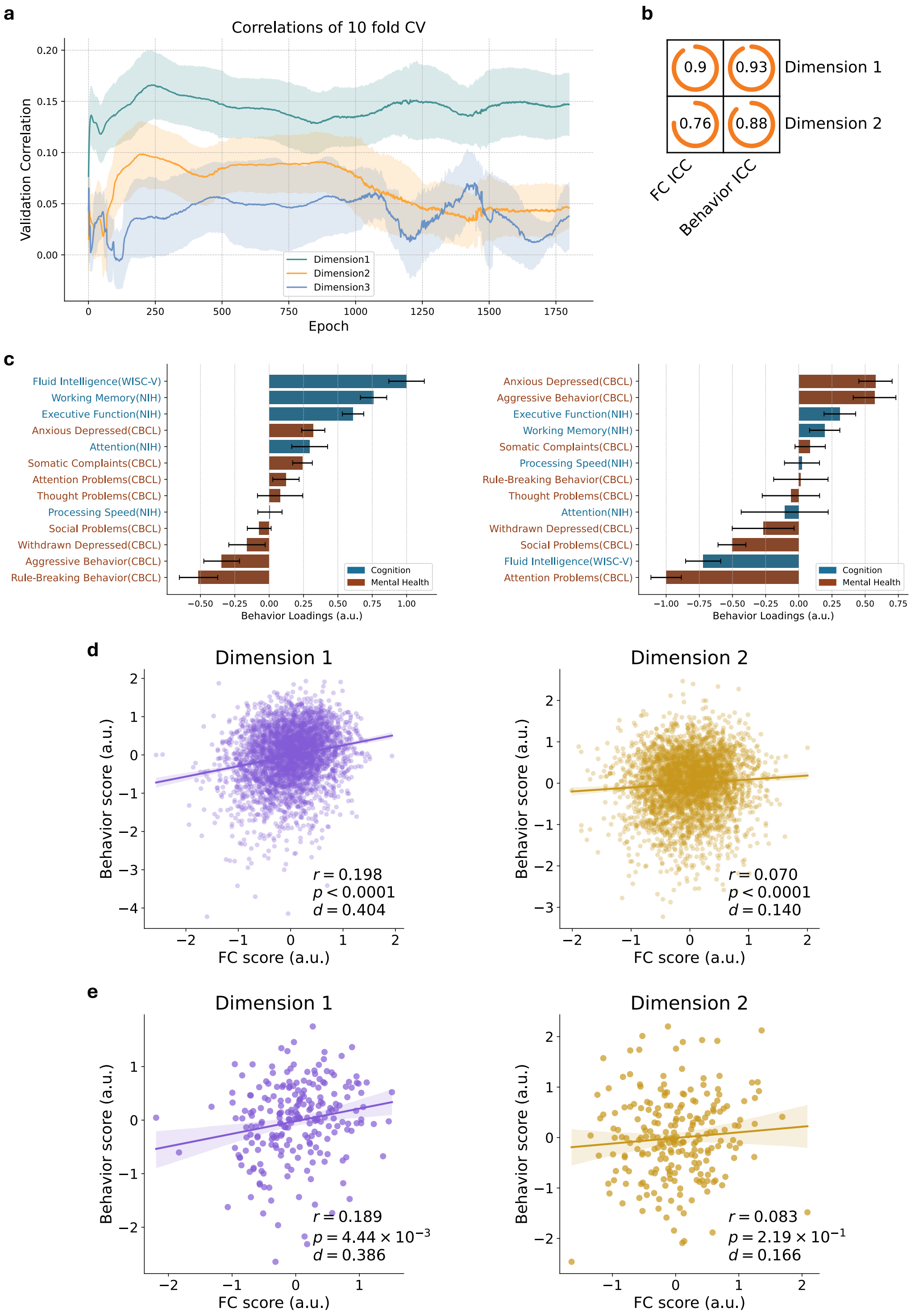

**Figure S5. Ablation analysis of the DGCCA model.** Standard DGCCA without autoencoder-based latent representation. **a**, Cross-validated correlation curves for each dimension during training. Lines show fold-averaged values; shaded areas represent standard deviation across folds. **b**, Intraclass correlation (ICC) of canonical loadings across 10-fold cross-validation. **c**, Behavior loadings of cognitive measures and CBCL subscales on the first (left) and the second (right) canonical dimensions. Error bars indicate standard deviation across folds. **d**, Correlations between FC and behavior scores for the first two canonical dimensions in the ABCD cohort using 10-fold cross-validation. **e**, Correlations between FC and behavior scores for the two identified canonical dimensions in the HBN cohort.

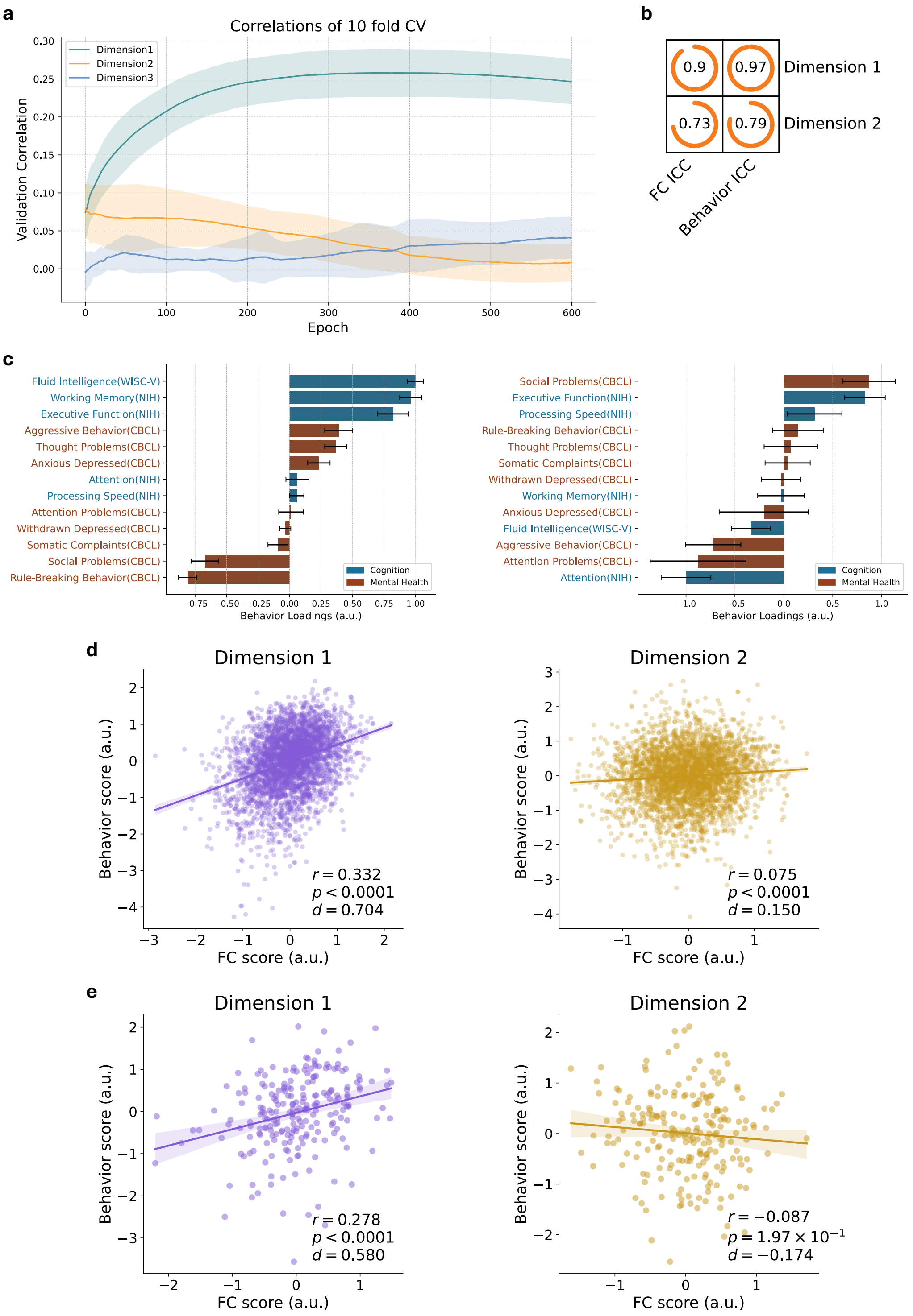

**Figure S6. Ablation analysis of the cAE + DGCCA model.** Contrastive autoencoder plus DGCCA without variational regularization. **a**, Cross-validated correlation curves for each dimension during training. Lines show fold-averaged values; shaded areas represent standard deviation across folds. **b**, Intraclass correlation (ICC) of canonical loadings across 10-fold cross-validation. **c**, Behavior loadings of cognitive measures and CBCL subscales on the first (left) and the second (right) canonical dimensions. Error bars indicate standard deviation across folds. **d**, Correlations between FC and behavior scores for the first two canonical dimensions in the ABCD cohort using 10-fold cross-validation. **e**, Correlations between FC and behavior scores for the two identified canonical dimensions in the HBN cohort.

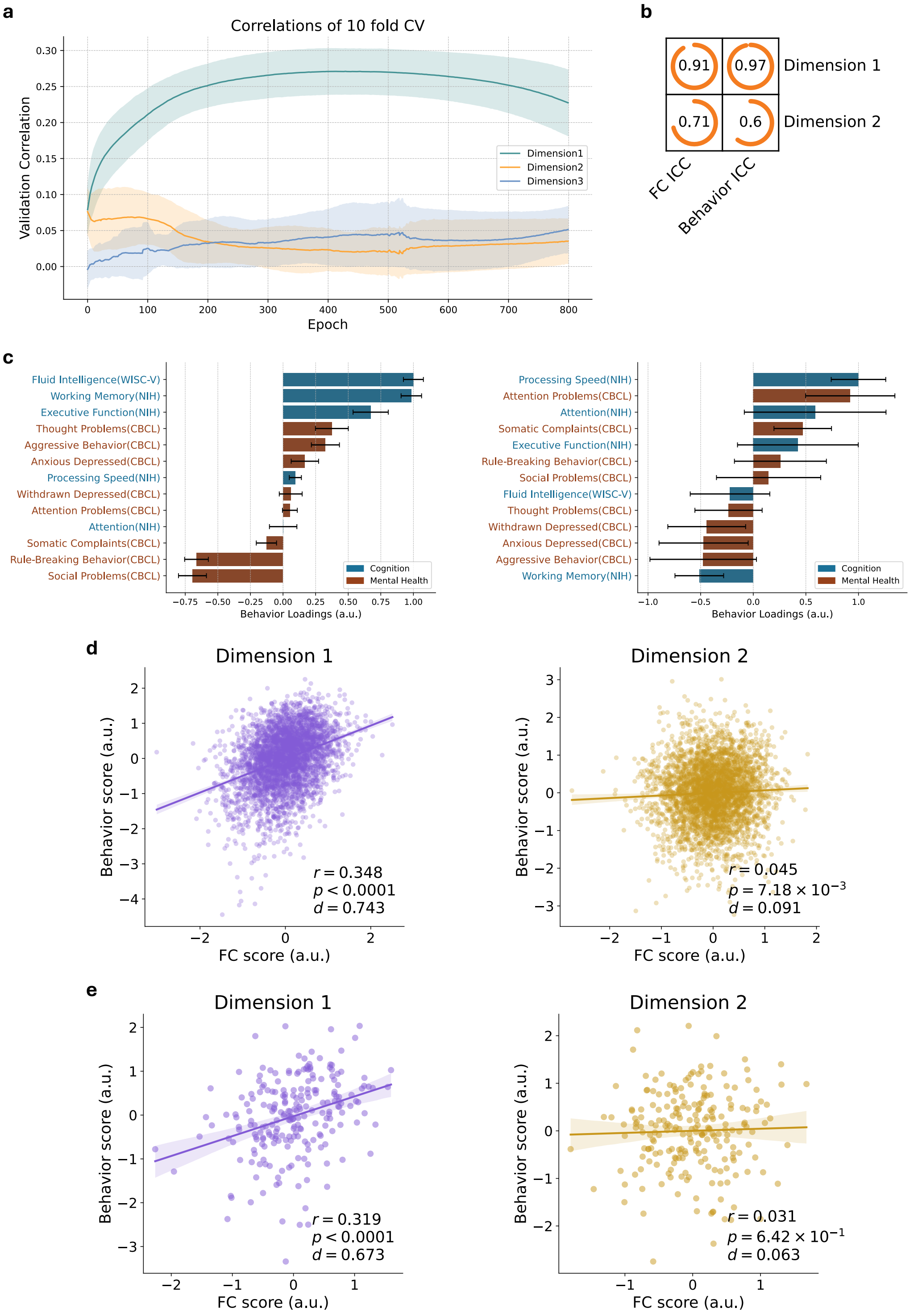

**Figure S7. Ablation analysis of the VAE + DGCCA model.** VAE plus DGCCA without contrastive mechanism. **a**, Cross-validated correlation curves for each dimension during training. Lines show fold-averaged values; shaded areas represent standard deviation across folds. **b**, Intraclass correlation (ICC) of canonical loadings across 10-fold cross-validation. **c**, Behavior loadings of cognitive measures and CBCL subscales on the first (left) and the second (right) canonical dimensions. Error bars indicate standard deviation across folds. **d**, Correlations between FC and behavior scores for the first two canonical dimensions in the ABCD cohort using 10-fold cross-validation. **e**, Correlations between FC and behavior scores for the two identified canonical dimensions in the HBN cohort.

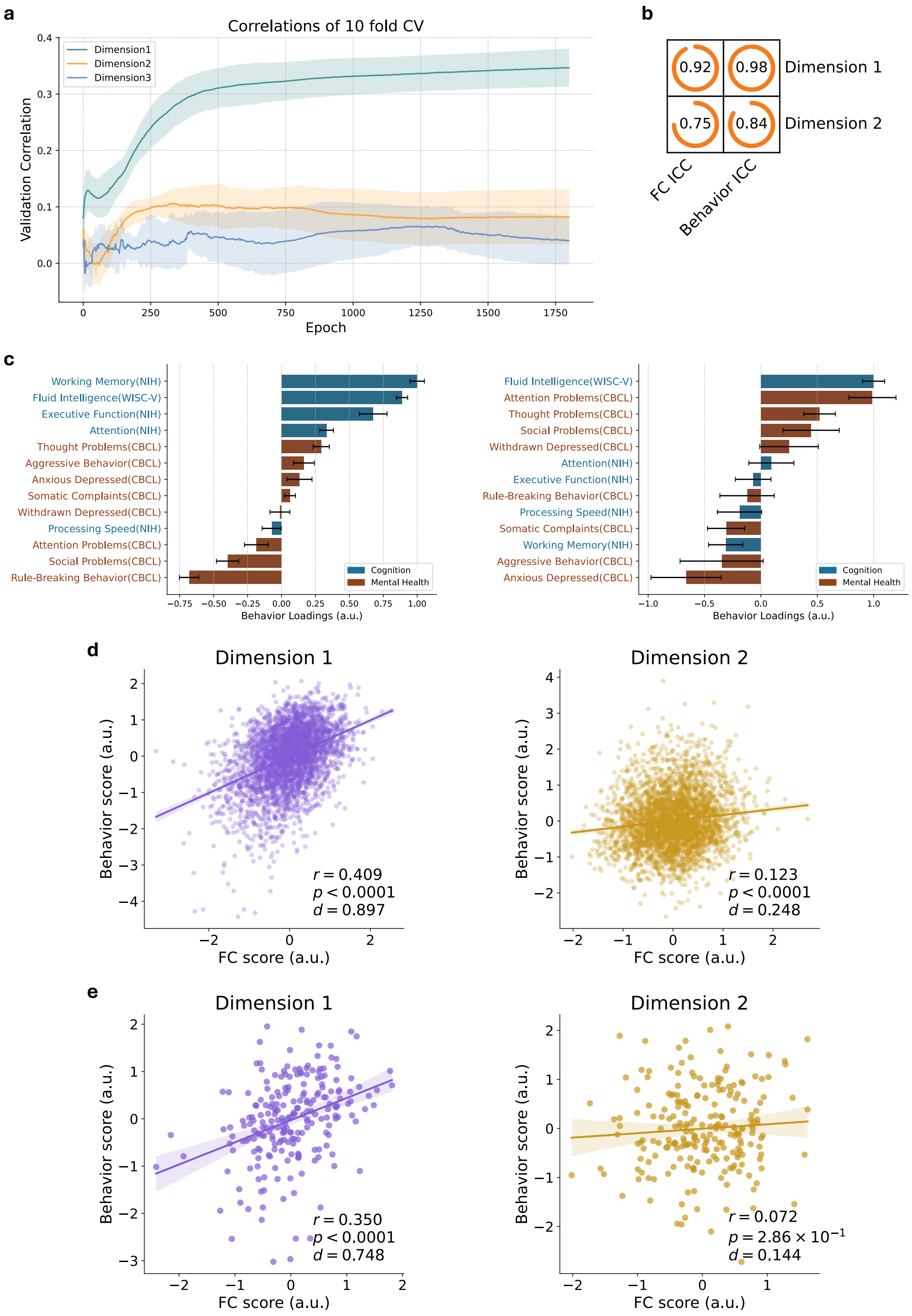

**Figure S8. Ablation analysis of the cPCA + sCCA model.** Contrastive PCA followed by sparse CCA[15,16], in the absence of nonlinear mapping and unified optimization. **a**, Intraclass correlation (ICC) of canonical loadings across 10-fold cross-validation. **b**, Behavior loadings of cognitive measures and CBCL subscales on the first (left) and the second (right) canonical dimensions. Error bars indicate standard deviation across folds. **c**, Correlations between FC and behavior scores for the first two canonical dimensions in the ABCD cohort using 10-fold cross-validation. **d**, Correlations between FC and behavior scores for the two identified canonical dimensions in the HBN cohort.

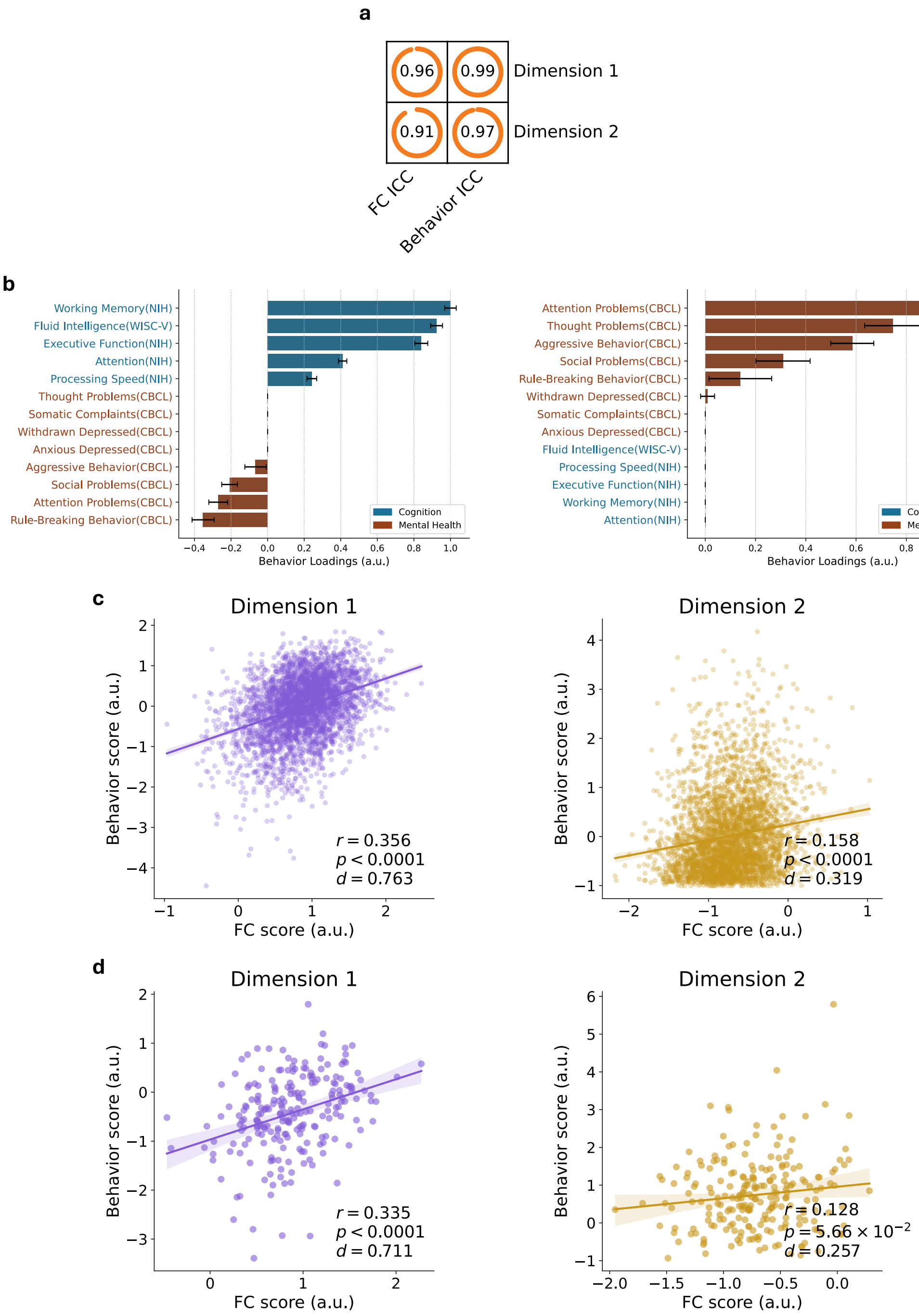

**Figure S9. Summary of stability and correlation ablation analyses.** Stability is shown as intraclass correlation (ICC) of canonical loadings across 10-fold cross-validation, and brain–behavior associations are shown as correlations between FC and behavior scores in the ABCD cohort ($r_{\text{CV\_ABCD}}$) and in the HBN cohort ($r_{\text{HBN}}$). Statistical differences between DeCoDE and baseline methods were evaluated using paired bootstrap resampling with 20,000 iterations and Fisher $z$ transformation, followed by FDR correction ($q < 0.05$). **a**, Dimension 1. **b**, Dimension 2.

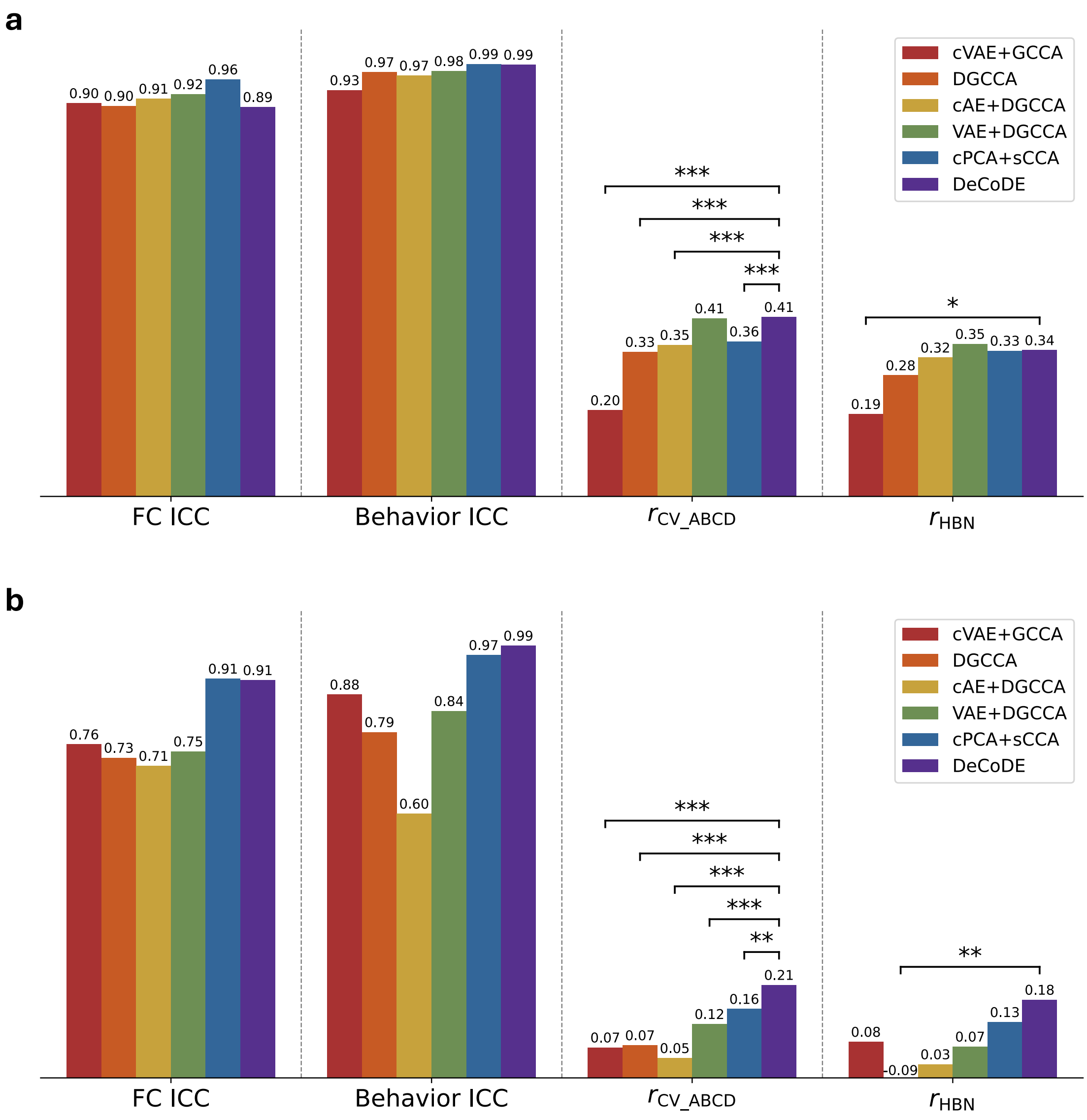

**Figure S10. OMP sparsity trends and approximated brain–behavior correlations.** **a**, Correlation trends across OMP sparsity levels (number of non-zero connections), fitted using DeCoDE-derived brain–behavior dimensions. Dashed lines indicate the DeCoDE correlation results as reference. A sparsity of $k = 500$ was adopted in this study based on observation. **b**, Correlations between FC scores obtained by projecting participant FC vectors onto OMP-estimated loading vectors ($k = 500$) and behavior scores in ABCD youth diagnosed with ADHD or AXD (N=3,508; 10-fold cross-validated). **c**, Correlations between FC scores in age-matched HBN youth with ADHD or AXD (N=224), obtained by applying the reference OMP projection ($k = 500$) estimated from the 3,508 diagnosed participants in ABCD.

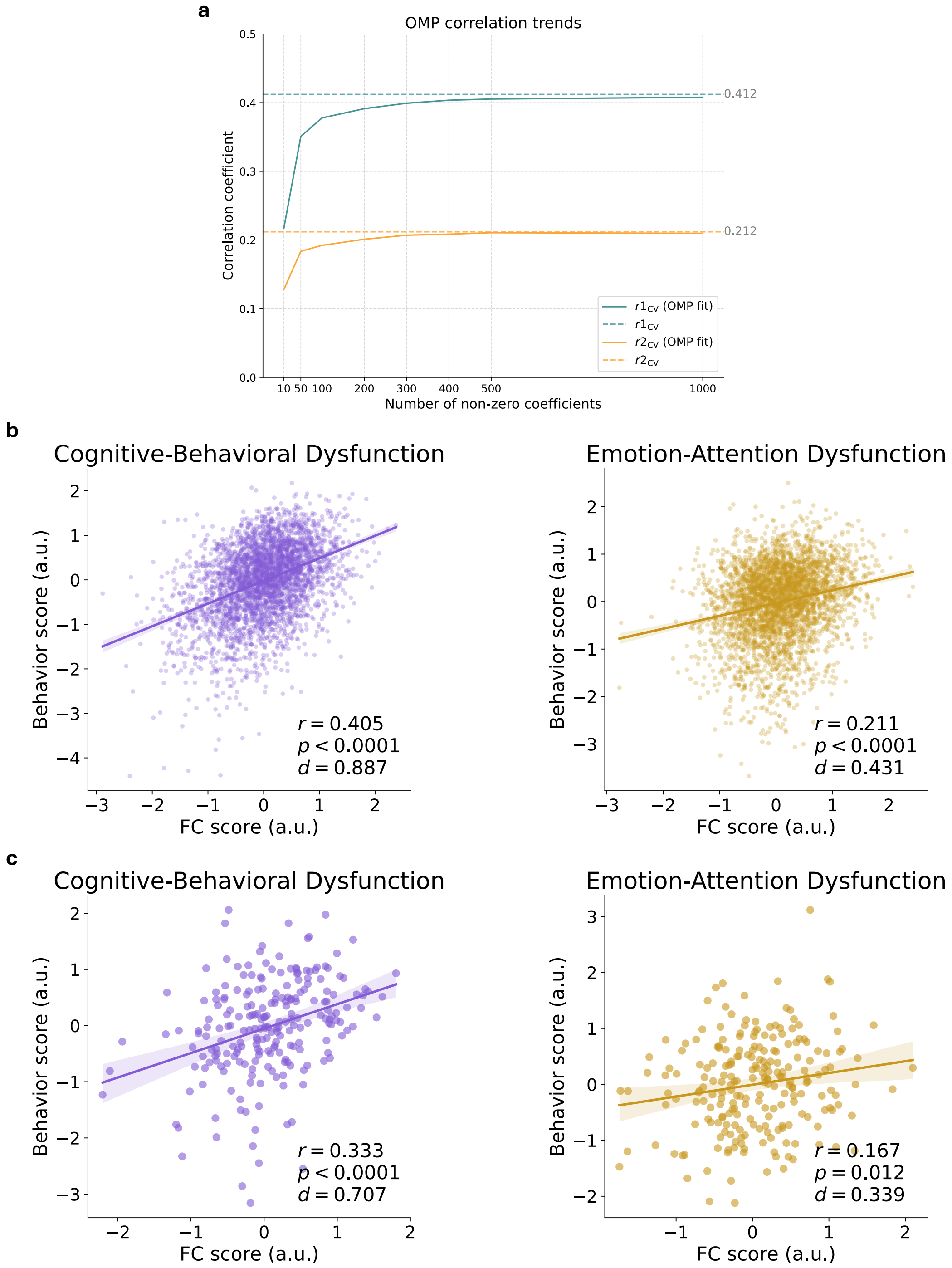

**Figure S11. OMP-derived FC loadings.** **a**, Sorted absolute OMP weights (top) and gap curves (bottom) at sparsity $k = 500$. Both dimensions show a sharp drop in weight magnitude within the top 20 ranks, with gap peaks around rank 20 separating the strongest weights from the remainder, supporting the choice of visualizing the top 20 connections in the main text (Fig. 2e–f). **b**, Full set of 500 nonzero FC loadings estimated with OMP. Node size reflects ROI strength, defined as the mean absolute weight of selected connections incident on each ROI. **c**, Network-level importance of OMP loadings, summarized as within-network and pairwise inter-network measures (see Method 10).

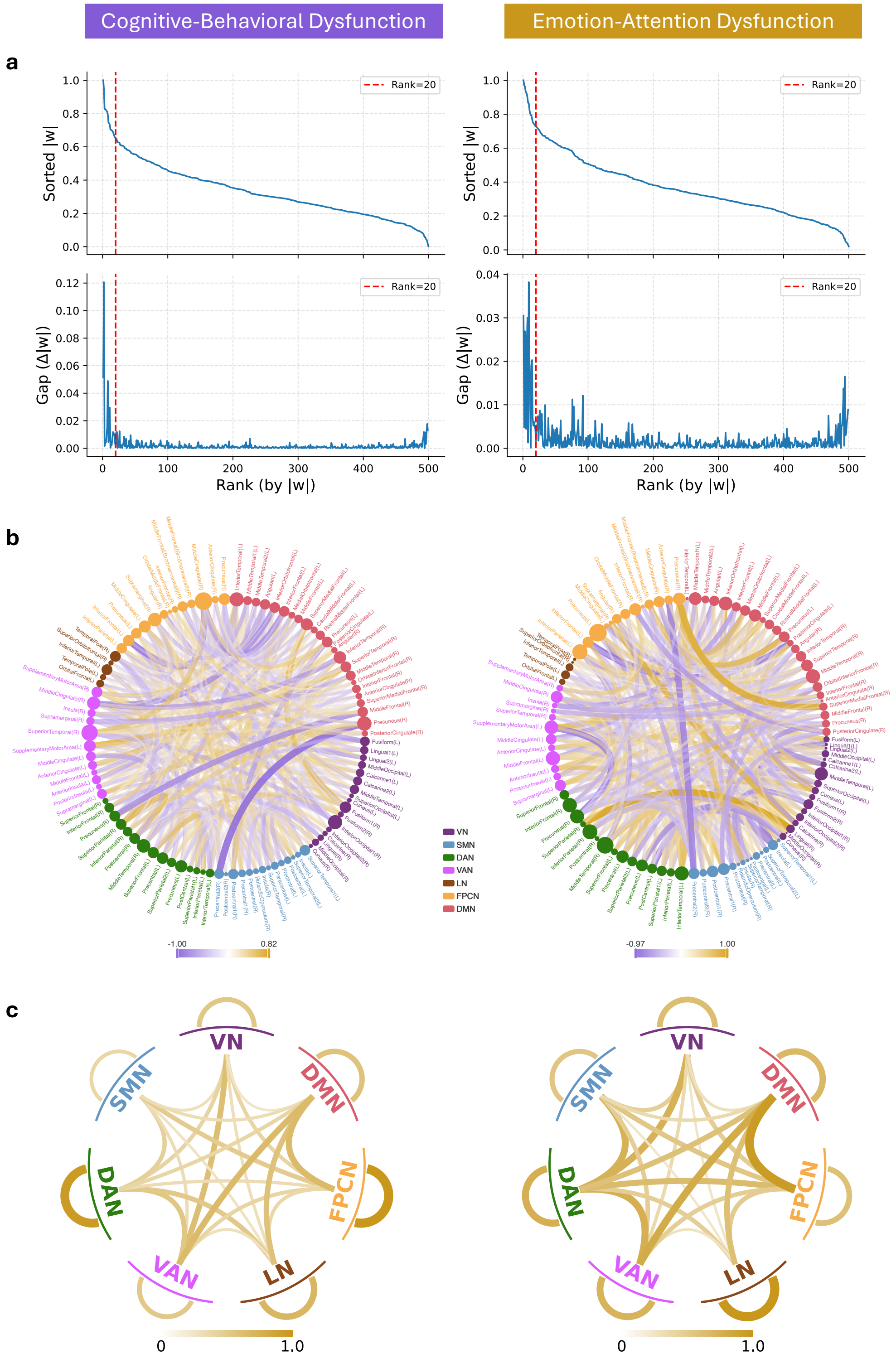

**Figure S12. Clustering model selection and stability evaluation.** **a**, Bayesian Information Criterion (BIC) and Akaike Information Criterion (AIC) values estimated from 100 repetitions of Gaussian mixture modeling on 80% randomly subsampled data. Lines represent the mean BIC/AIC across repetitions, and shaded areas indicate the range between minimum and maximum values. Lower values indicate better model fit. **b**, Clustering stability scores estimated using 100 repetitions of 5-fold cross-validation with 80% subsampled data. Left panel: Original clustering stability, measured by clustering accuracy (ACC), adjusted Rand index (ARI), and normalized mutual information (NMI), alongside chance-level stability computed by applying the same procedure to randomly shuffled data. Error bars indicate standard deviations across repetitions. Right panel: Adjusted stability scores were calculated as $(\text{Original} - \text{Chance})/(1 - \text{Chance})$ to account for baseline agreement due to chance.

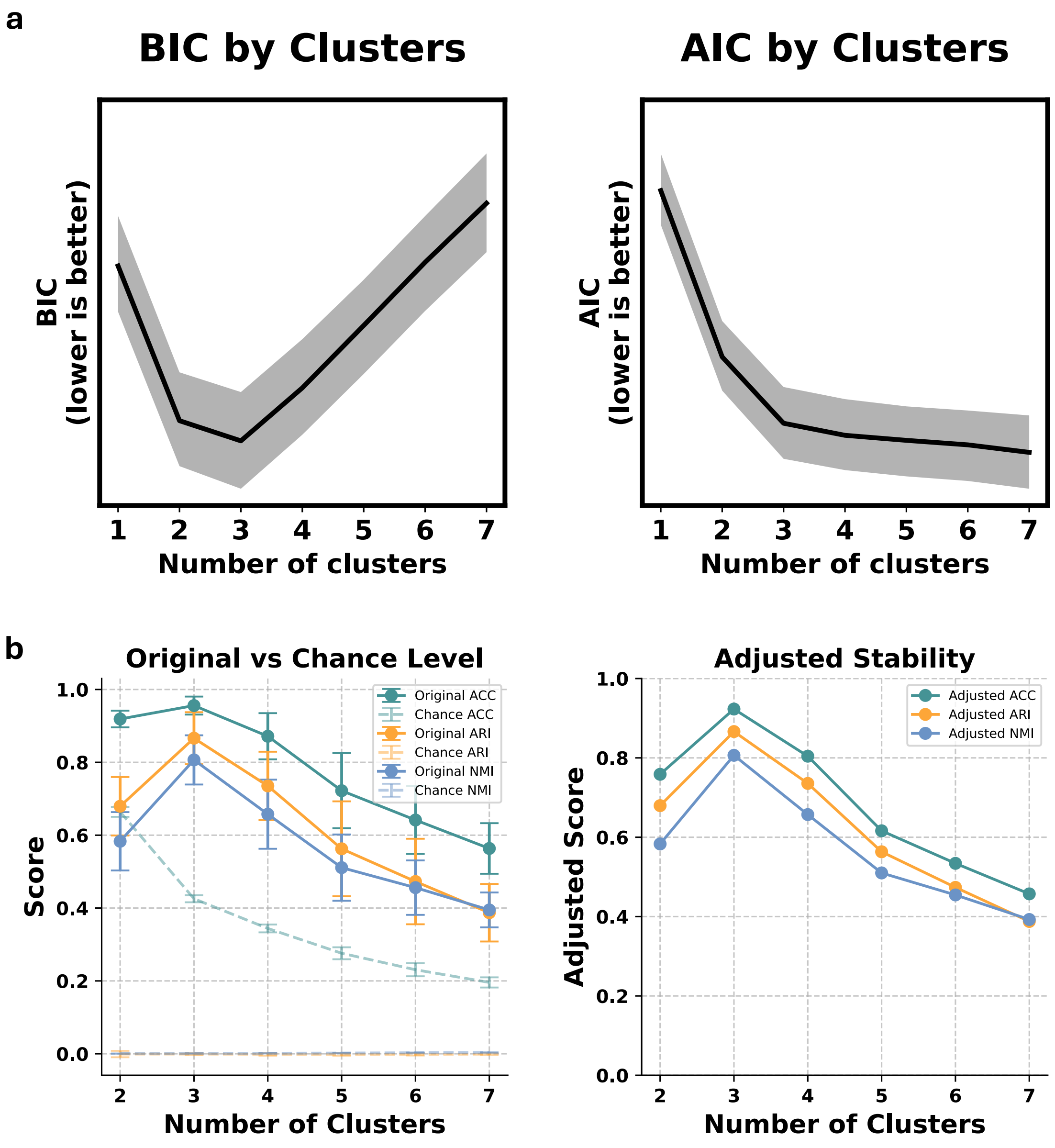

**Figure S13. CBCL symptom profiles across transdiagnostic biotypes.** CBCL subscale scores (mean ± 95% CI) are shown for three biotypes identified in: **a**, clinically diagnosed participants with ADHD or AXD from the ABCD dataset (N=3,508); **b**, age-matched participants with ADHD or AXD from the HBN dataset (N=224); **c**, ABCD late-onset individuals (N=693) assessed at baseline, who had not yet received a diagnosis but were later classified as ADHD or AXD during follow-up. Group differences were assessed using Kruskal–Wallis tests across the three biotypes, followed by FDR correction ($q < 0.05$) across subscales. Asterisks indicate subscales with significant group effects after correction. Subscale labels are color-coded to mark the biotype with the highest mean score.

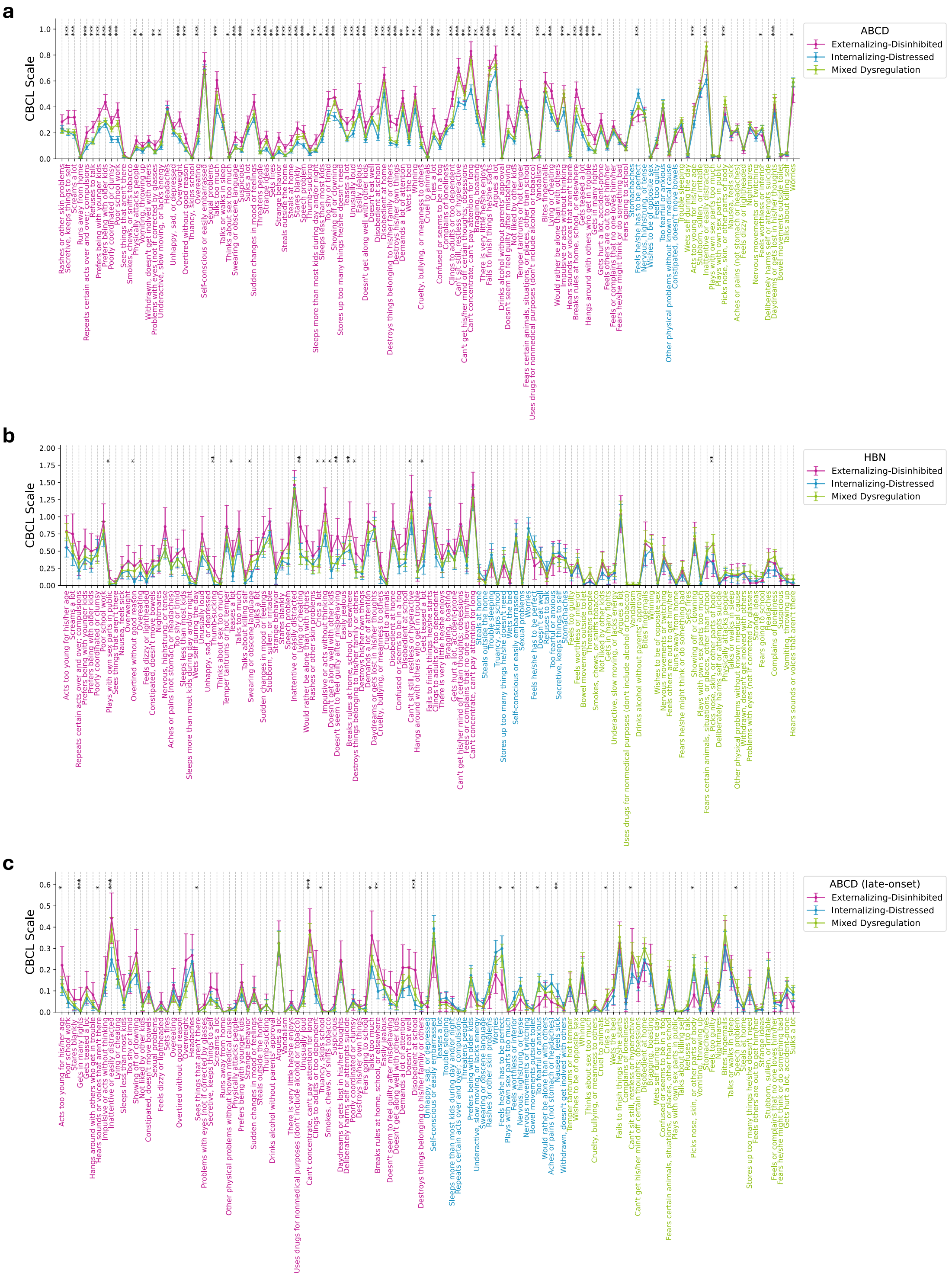

**Figure S14. Pairwise comparisons of cognitive and mental health scales across transdiagnostic biotypes.** **a**, Clinically diagnosed participants with ADHD or AXD from the ABCD dataset (N=3,508); **b**, Age-matched participants with ADHD or AXD from the HBN dataset (N=224). Cognitive scales were approximately normally distributed and underwent outlier removal using a 3-standard-deviation threshold; mental health scales were processed using median absolute deviation with a threshold of 4. Pairwise group comparisons were conducted using Mann–Whitney U tests, with FDR correction ($q < 0.05$) across all comparisons. Asterisks denote pairwise significant differences after correction.

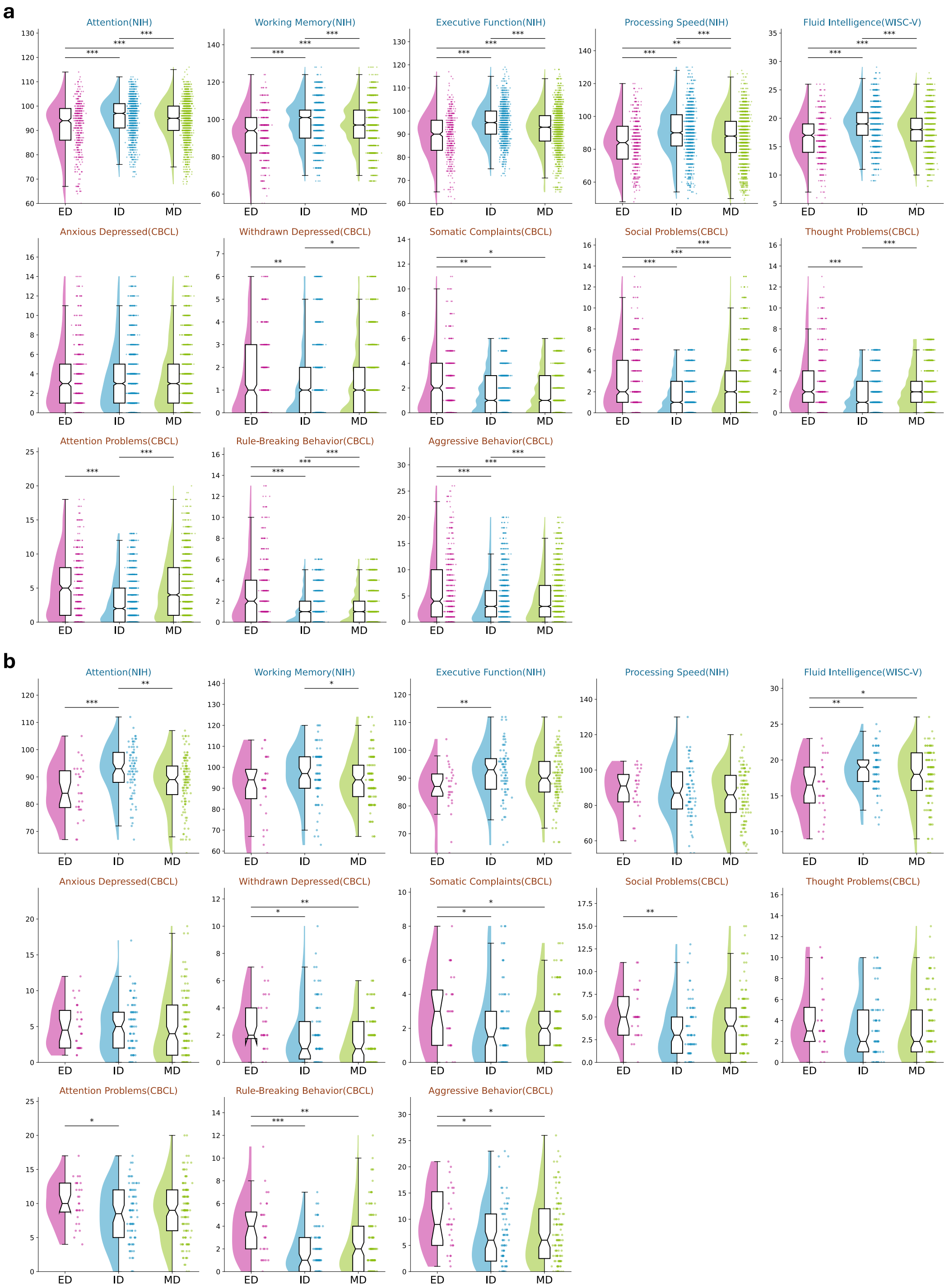

**Figure S15. Group-level comparisons across cognitive, mental health, personality, and contextual domains in diagnosed participants from the ABCD dataset.** Domain-specific profiles are shown for individuals diagnosed with ADHD or AXD at baseline (N=3,508), stratified into three neurobehavioral biotypes and compared to controls. **a**, Cognitive performance, including working memory, executive function, processing speed, and fluid intelligence; **b**, Mental health symptoms derived from CBCL subscales; **c**, Personality traits encompassing BIS/BAS and UPPS-P constructs; **d**, Contextual and environmental factors such as sleep patterns, family conflict, and parental psychopathology. Outliers were removed using a 3-standard-deviation threshold for approximately normally distributed variables and the median absolute deviation method with a threshold of 4 for others. Group differences across biotypes were assessed using the Kruskal–Wallis test, followed by FDR correction ($q < 0.05$) within each domain. Asterisks denote scales with significant biotype-level differences after correction. Radar positions indicate normalized group means; error bars represent 95% confidence intervals.

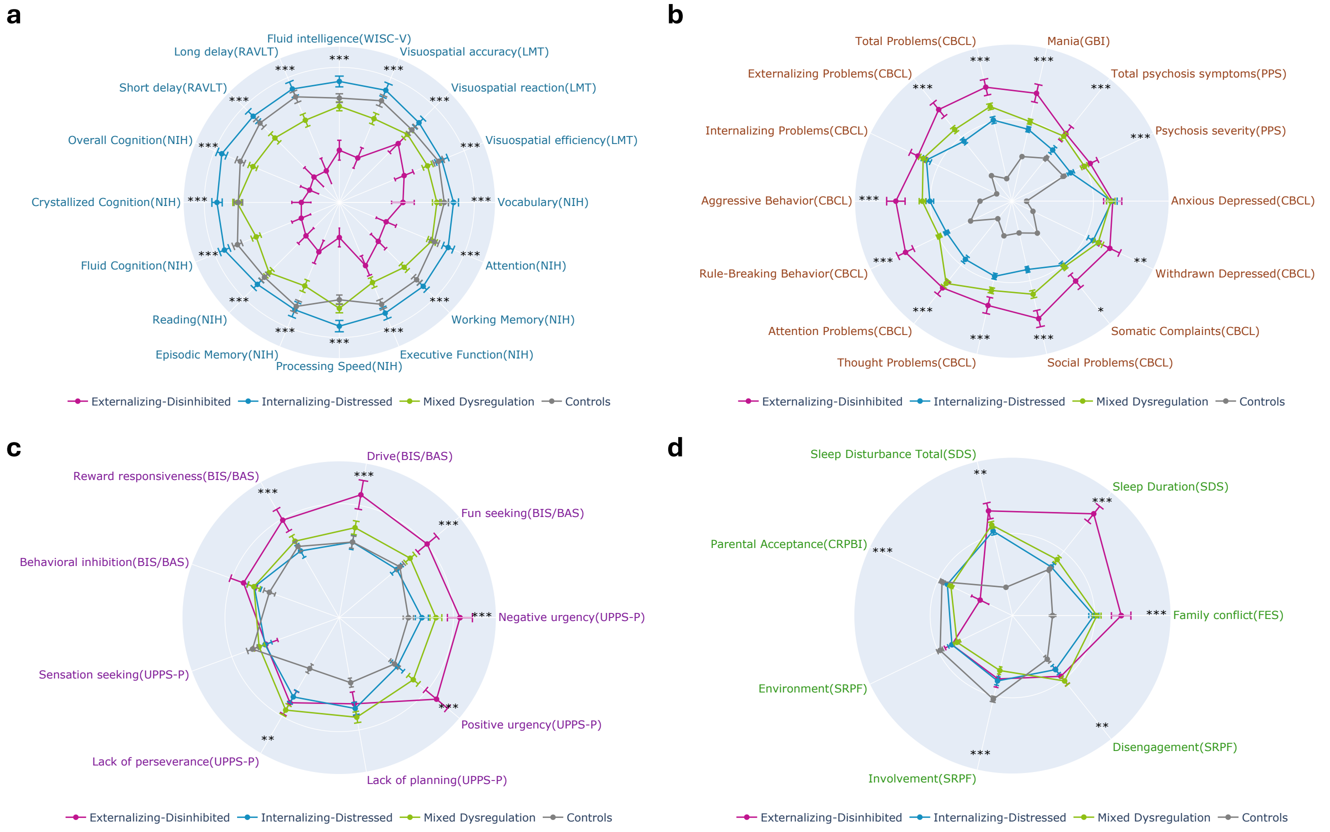

**Figure S16. Correlational structure between cognitive/mental health measures and contextual factors across biotypes and controls.** Text labels display correlation $r$. The colorbar shows FDR-corrected correlation $p_{FDR}$ values. Bubble size reflects absolute Cohen's $d$. Bubble border color indicates FDR-corrected significance of biotype–control differences. **a**, Pearson correlations with covariates including sex, age, family income, parental education, and marital status. For the binary variables sex and marital status, higher values indicate female and married/cohabiting. **b**, Partial correlations with contextual variables such as Sleep Disturbance, Sleep Duration, Family Conflict, School Involvement, Parental Acceptance, and Youth Alcohol Use, controlling for covariates. **c**, Partial correlations with familial psychiatric history variables including Parental Alcohol Use, Drug Use, Antisocial Behavior, Suicide Attempt, Mania, Depression, Paranoia, Nervous Breakdown, and Psychiatric Hospitalization, controlling for covariates.

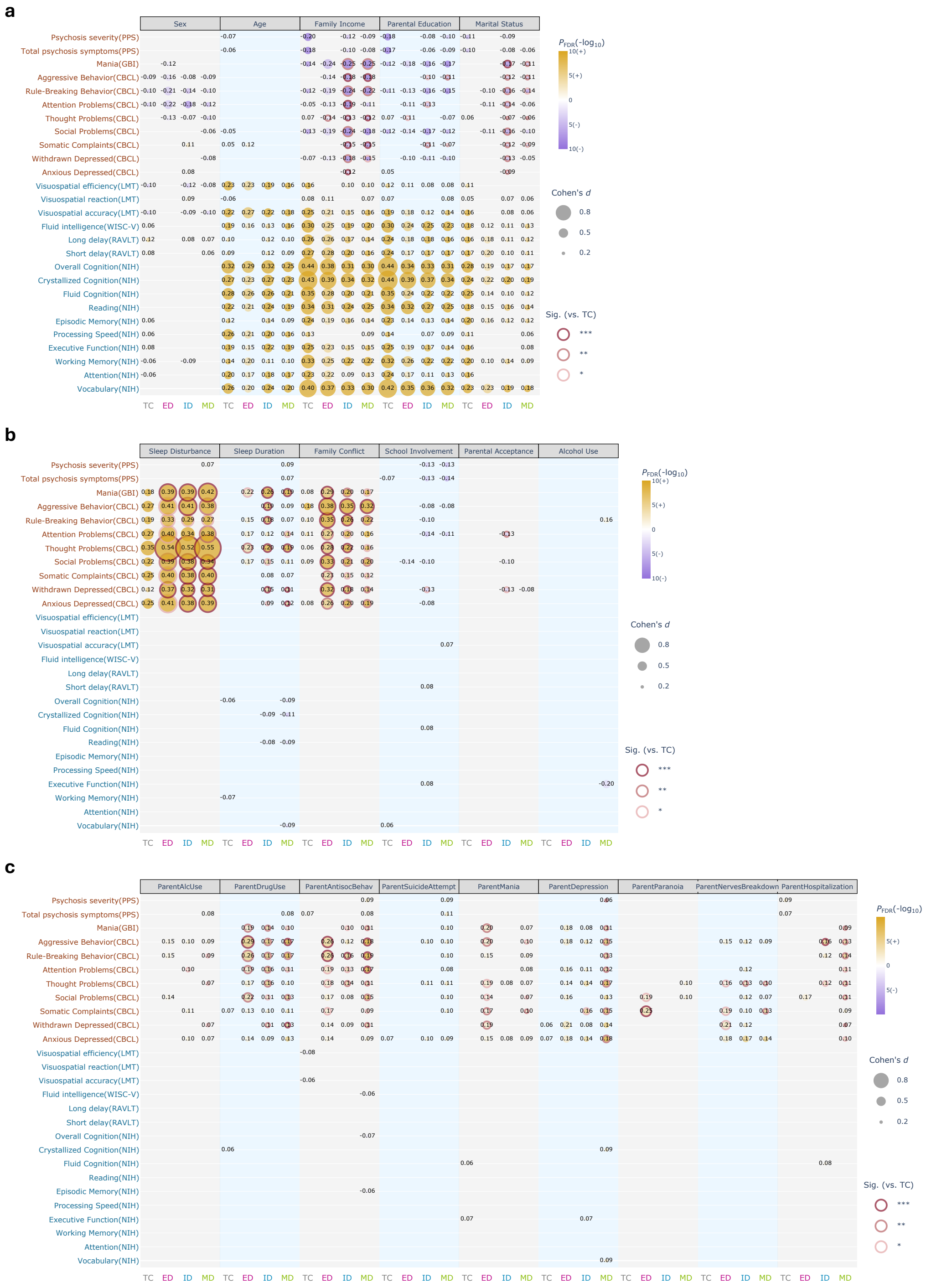

**Figure S17. Group-mean original FC at ROI and network levels.** Group-mean FC matrices for controls and each biotype (ED, ID, MD), computed in the original FC space and averaged across participants after subject-wise demeaning. Colorbars show group-mean connectivity (Fisher $z$ units). **Bottom left of each panel:** ROI–ROI mean connectivity. **Top right of each panel:** Network-level mean connectivity within and between functional networks. Edge thickness and opacity encode connectivity strength.

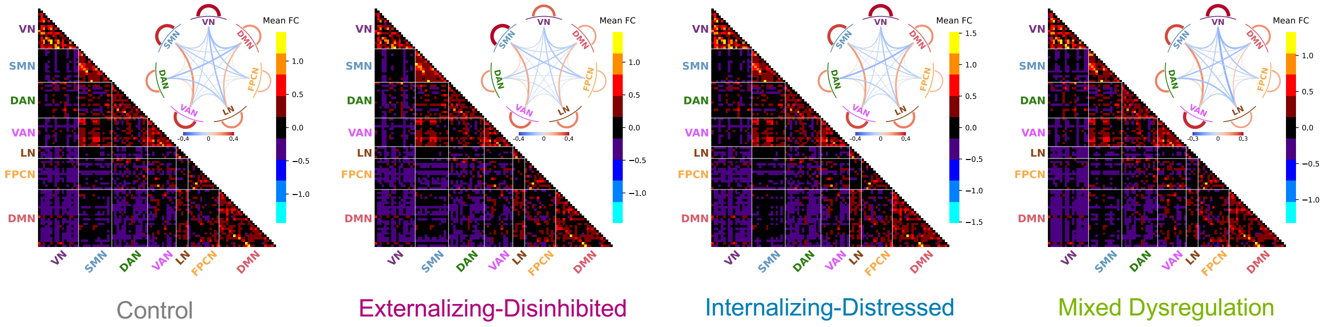

**Figure S18. Pairwise functional connectivity differences between biotypes.** Biotypes were compared in original FC space at both ROI and network levels using two-sample $t$-tests with FDR correction ($q < 0.05$). Colorbars show effect size (Cohen's $d$). Warm colors indicate hyperconnectivity in the first biotype relative to the second, and cool colors indicate hypoconnectivity. For all contrasts (e.g., ID vs. ED), the second biotype served as the reference group, such that positive values denote higher FC in the first biotype and negative values denote lower FC. **Bottom left of each panel:** ROI-level FC differences. **Top right of each panel:** Network-level FC differences computed as mean connectivity within and between functional networks. Edge thickness and opacity encode effect magnitude.

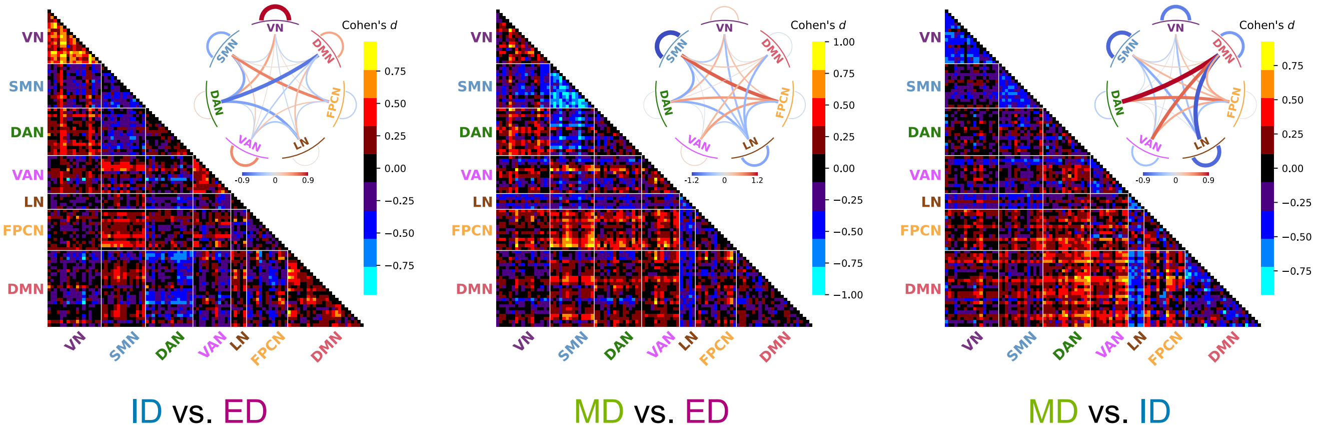

**Figure S19. Longitudinal modeling of sMRI and DTI metrics in ABCD-diagnosed participants.** Linear mixed-effects models were used to characterize age-related changes in sMRI and four DTI metrics across biotypes in children diagnosed with ADHD or AXD at baseline (N=3,508). Statistical significance of fixed effects (sex, parental education, family income, marital status, and intracranial volume) was assessed using Type III Wald $\chi^2$ tests with FDR correction ($q < 0.05$), and effect sizes were quantified using partial eta squared. Only covariates with significant age interactions were retained in the final models. Predicted trajectories represent model-based estimates with 95% confidence intervals; observed data are shown across baseline and follow-up visits. FDR-adjusted $p$-values for age-by-biotype interactions are reported in each panel. **a**, Partial eta squared values for age-interacted covariates in sMRI models. Sex, parental education, family income, and intracranial volume were retained. **b**, Predicted trajectories of sMRI metrics (cortical volume, cortical thickness, cortical surface area, and subcortical volume) for each biotype. **c**, Observed sMRI values across visits. Dashed gray lines represent the mean trajectory of controls. **d**, Partial eta squared values for age-interacted covariates in DTI models. Only sex and intracranial volume were retained. **e**, Predicted trajectories of DTI metrics (fractional anisotropy, mean diffusivity, longitudinal diffusivity, and transverse diffusivity) for each biotype. **f**, Observed DTI values across visits. Dashed gray lines represent the control mean.

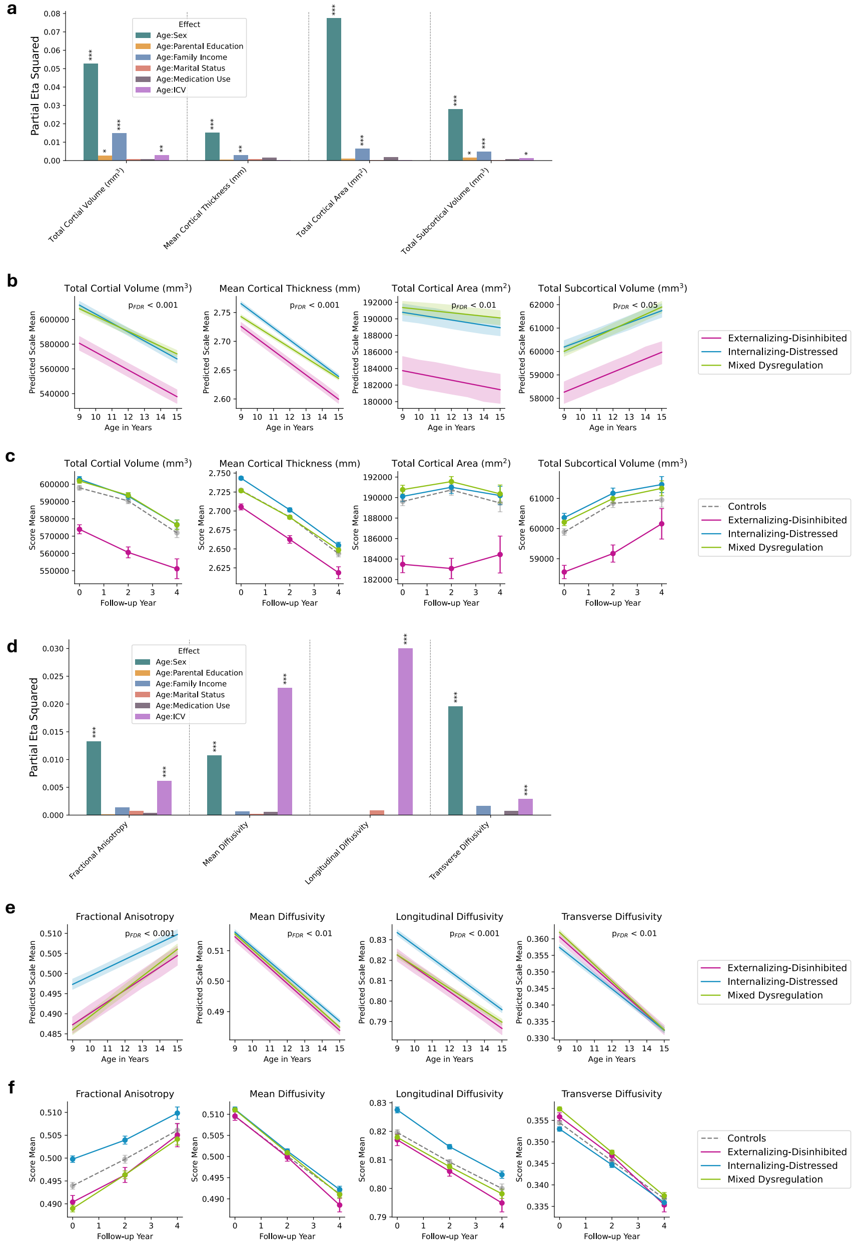

**Figure S20. Biotype-specific regional brain differences across cortical and subcortical ROIs. a**, Desikan–Killiany cortical atlas[94]. **b**, ASEG subcortical atlas[95]. **c–e**, Group differences for each biotype versus controls. Within each panel, regional cortical volume, cortical thickness, cortical surface area, and subcortical volume were compared using linear regression controlling for age, sex, parental education, family income, marital status, and intracranial volume, with FDR correction applied per metric ($q < 0.05$).

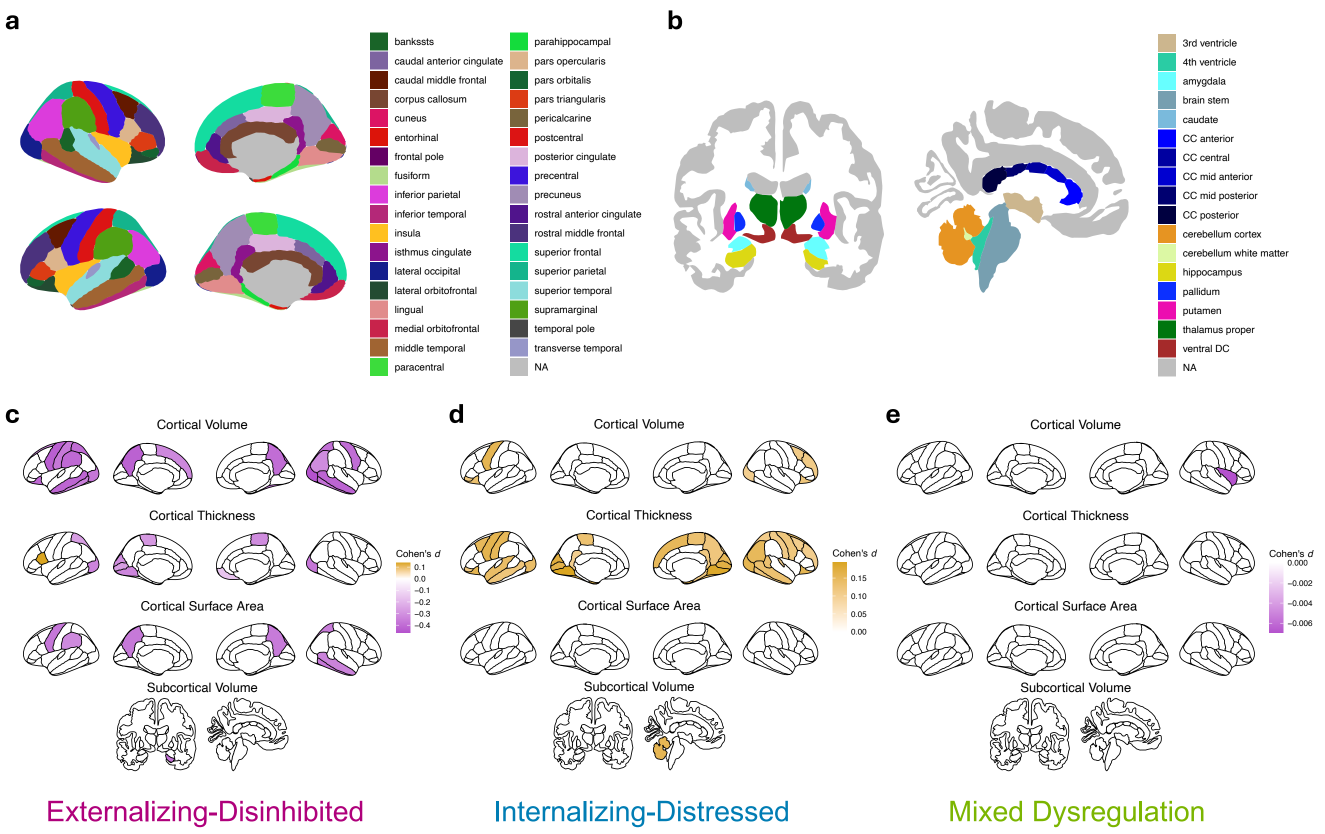

**Figure S21. Biotype-specific white matter differences across major tracts. a**, AtlasTrack template[96] of 35 white matter tracts. **b–d**, Group differences for each biotype versus controls. Within each panel, tract-averaged microstructural properties, including fractional anisotropy, mean diffusivity, longitudinal diffusivity, and transverse diffusivity, were compared using linear regression controlling for age, sex, parental education, family income, marital status, and intracranial volume, with FDR correction applied per metric ($q < 0.05$).

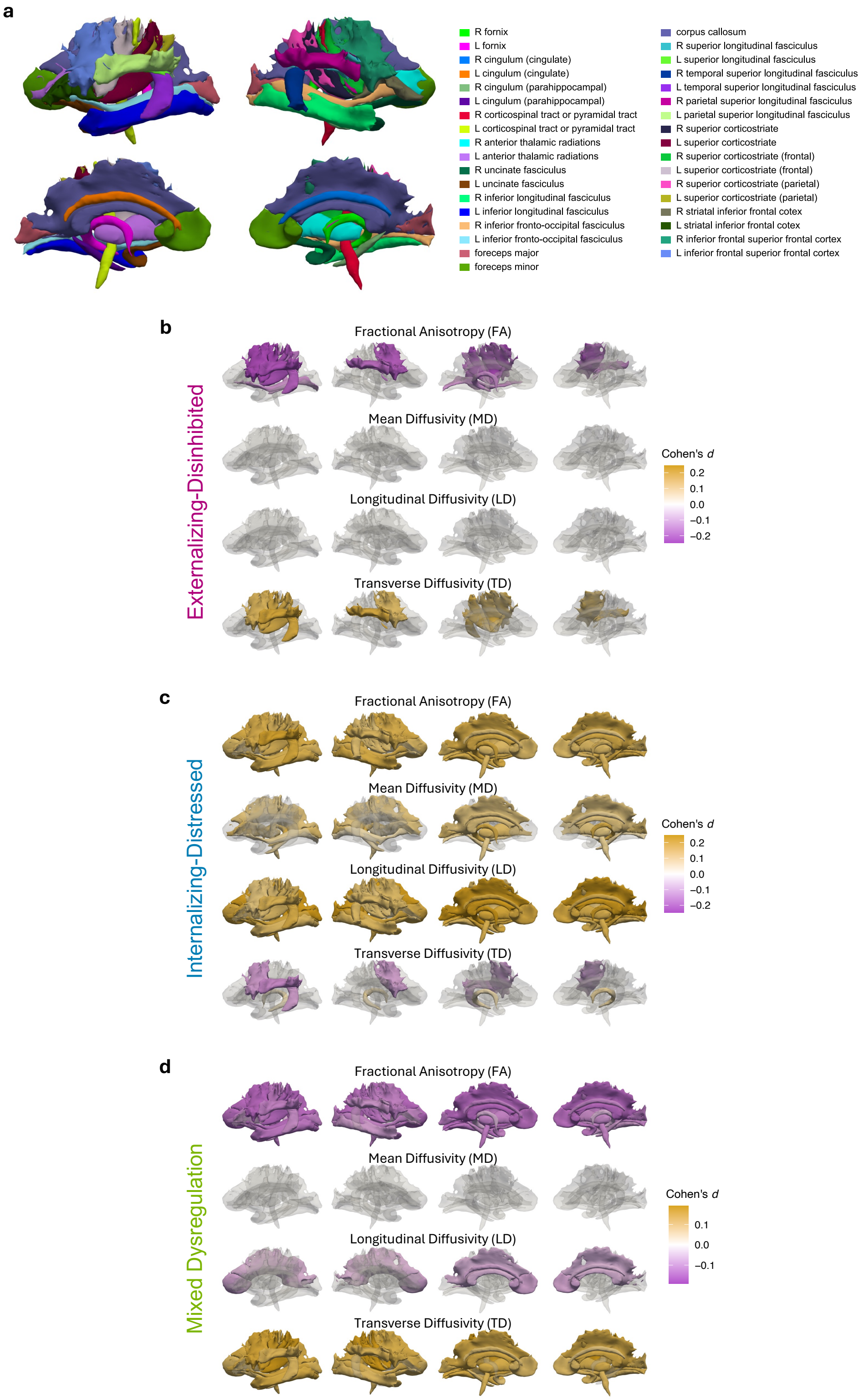

**Figure S22. Longitudinal modeling of CBCL subscales in ABCD-diagnosed participants.** Results from LME models predicting CBCL T scores across ages 9–15 in children diagnosed with ADHD or AXD at baseline (N=3,508). **a**, Partial eta squared values for the interaction of age with covariates (sex, parental education, family income, marital status, and medication use) based on Type III Wald $\chi^2$ tests. Medication use was parent-reported and included commonly prescribed stimulants, anxiolytics, or antidepressants. Biotype-specific medication use rates were: 20.4% (ED), 17.7% (ID), and 25.6% (MD). **b**, Predicted CBCL trajectories for each biotype based on the final LME model. Shaded areas represent 95% confidence intervals. FDR-corrected $p$-values for age-by-biotype interactions are displayed in each panel. Models included sex, parental education, family income, and medication use as significant covariates. **c**, Observed CBCL T scores at baseline and follow-up visits across biotypes. Dashed gray lines indicate the mean trajectory of controls.

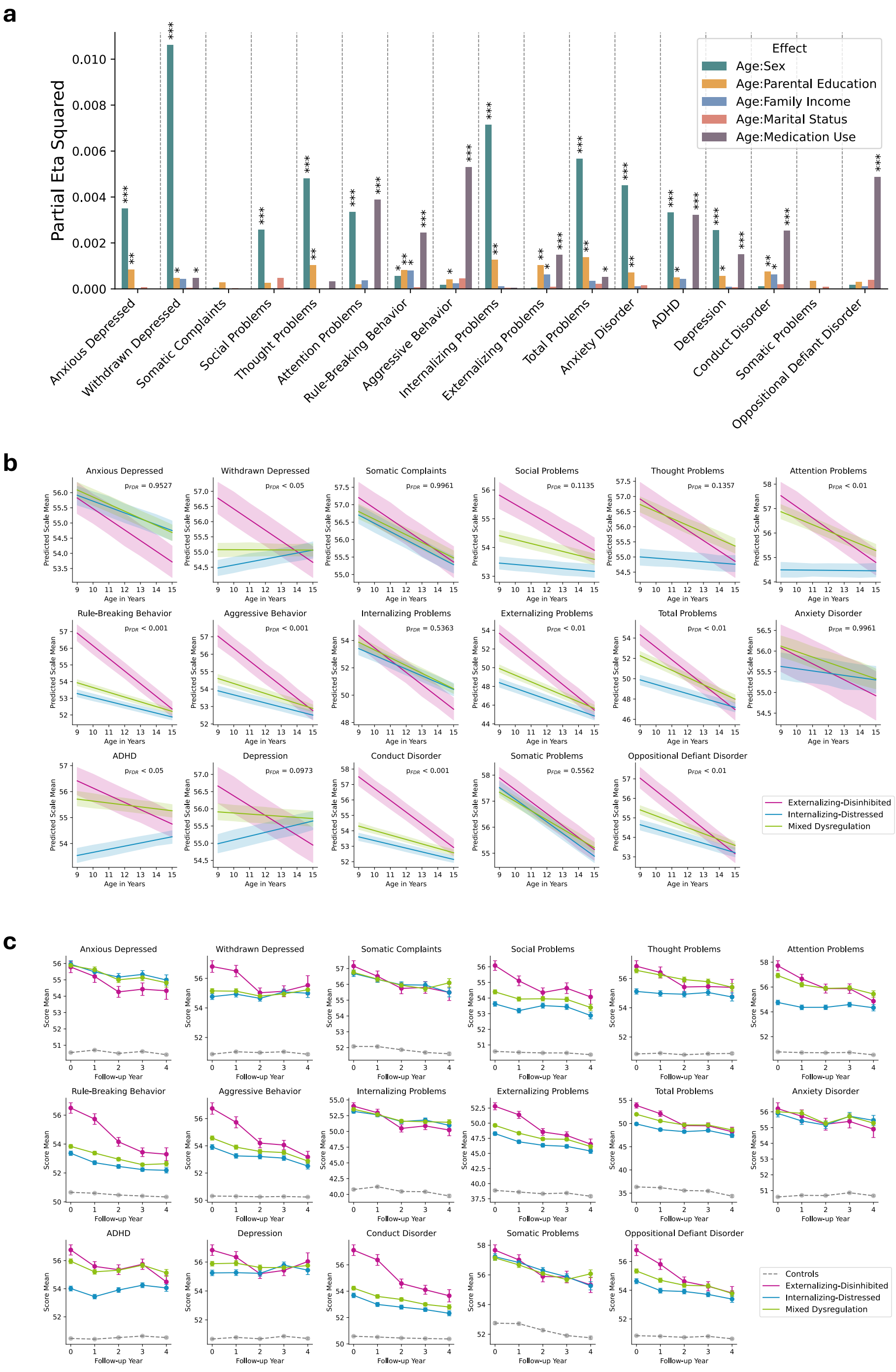

**Figure S23. Statistical validation of biotype prevalence in the independent HBN dataset.** The generalizability of the three-biotype prevalence identified in the ABCD dataset was evaluated in the independent HBN dataset using 1,000 random permutations. In each permutation, the FC features of the HBN dataset were independently shuffled within each feature column across subjects. **a**, Example biotype assignments (posterior probabilities) for a permuted HBN dataset. **b**, Null distribution of prevalence similarity scores. The red dashed line indicates the observed similarity from the original, non-permuted data.

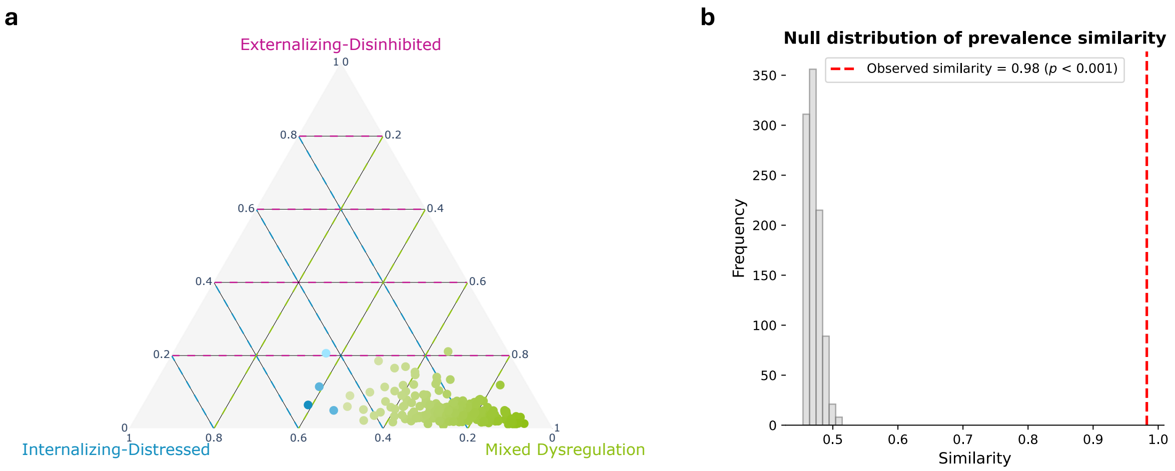

**Figure S24. Statistical validation of biotype-specific clinical and FC characteristics in the independent HBN dataset.** The reproducibility of the clinical and FC characteristics of the ABCD-derived biotypes was evaluated in the independent HBN dataset using 1,000 random permutations. In each permutation, the biotype labels in HBN were randomly shuffled across subjects while retaining the original FC features and derived FC scores. **a**, Null distribution of clinical profile similarity scores. **b**, Null distribution of FC score distributional similarity. **c**, Null distributions of FC pattern similarity for the ED, ID and MD biotypes respectively. In each panel, the red dashed line indicates the observed similarity from the original, non-permuted data.

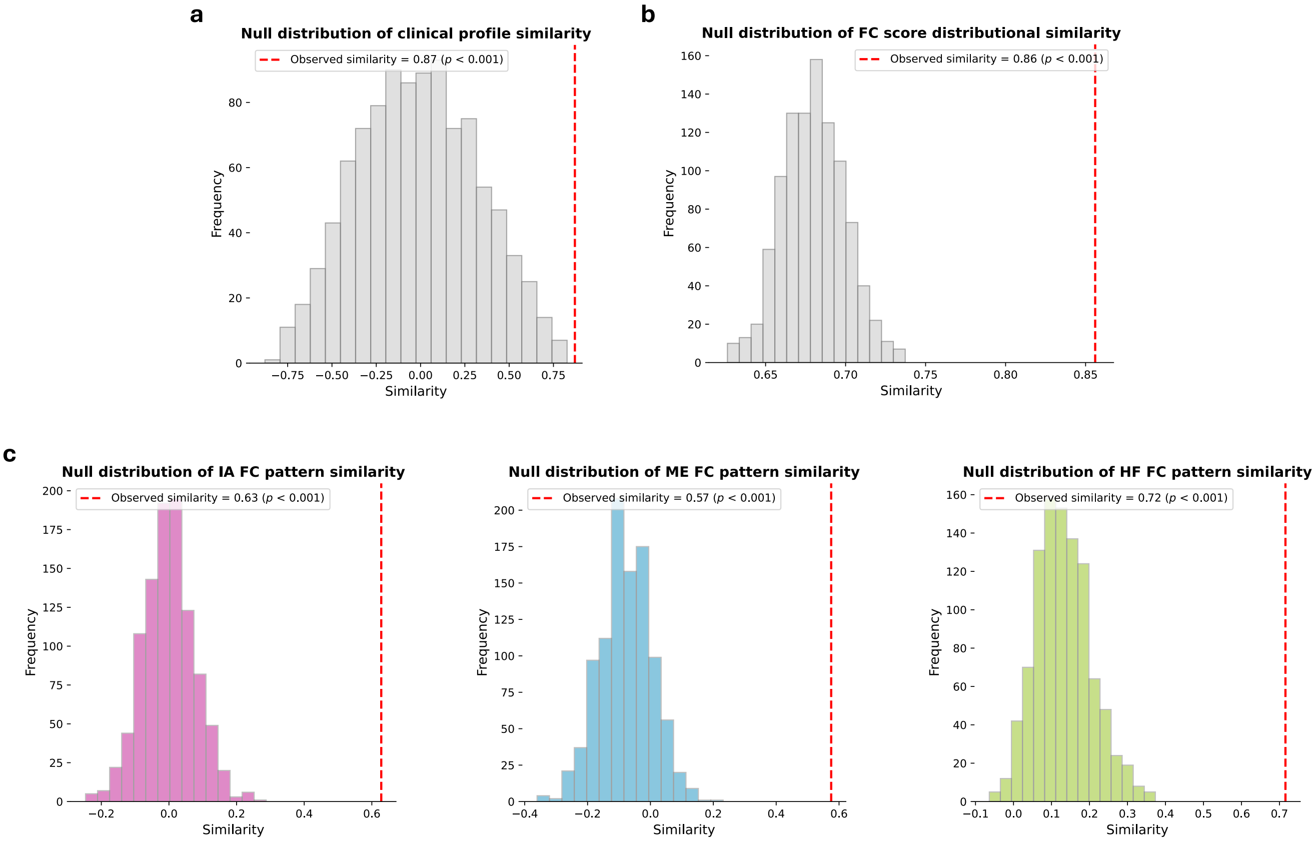

**Figure S25. Brain–behavior dimensions and biotypic expression in the ABCD late-onset cohort.** **a**, Canonical correlations between FC scores and behavior scores for the two identified dimensions at baseline. **b**, Posterior membership probabilities for 693 LO individuals, obtained by applying GMM derived from the diagnosed cohort to baseline FC scores. **c**, DSM diagnostic composition of each late-onset biotype, displaying the number of individuals who later developed ADHD or AXD during follow-up. **d**, Kernel density estimation plots of late-onset biotypes across the two identified brain–behavior dimensions. Contours (25%, 50%, 75%) depict density levels, and marginal boxplots on the top and right summarize score distributions along each dimension.

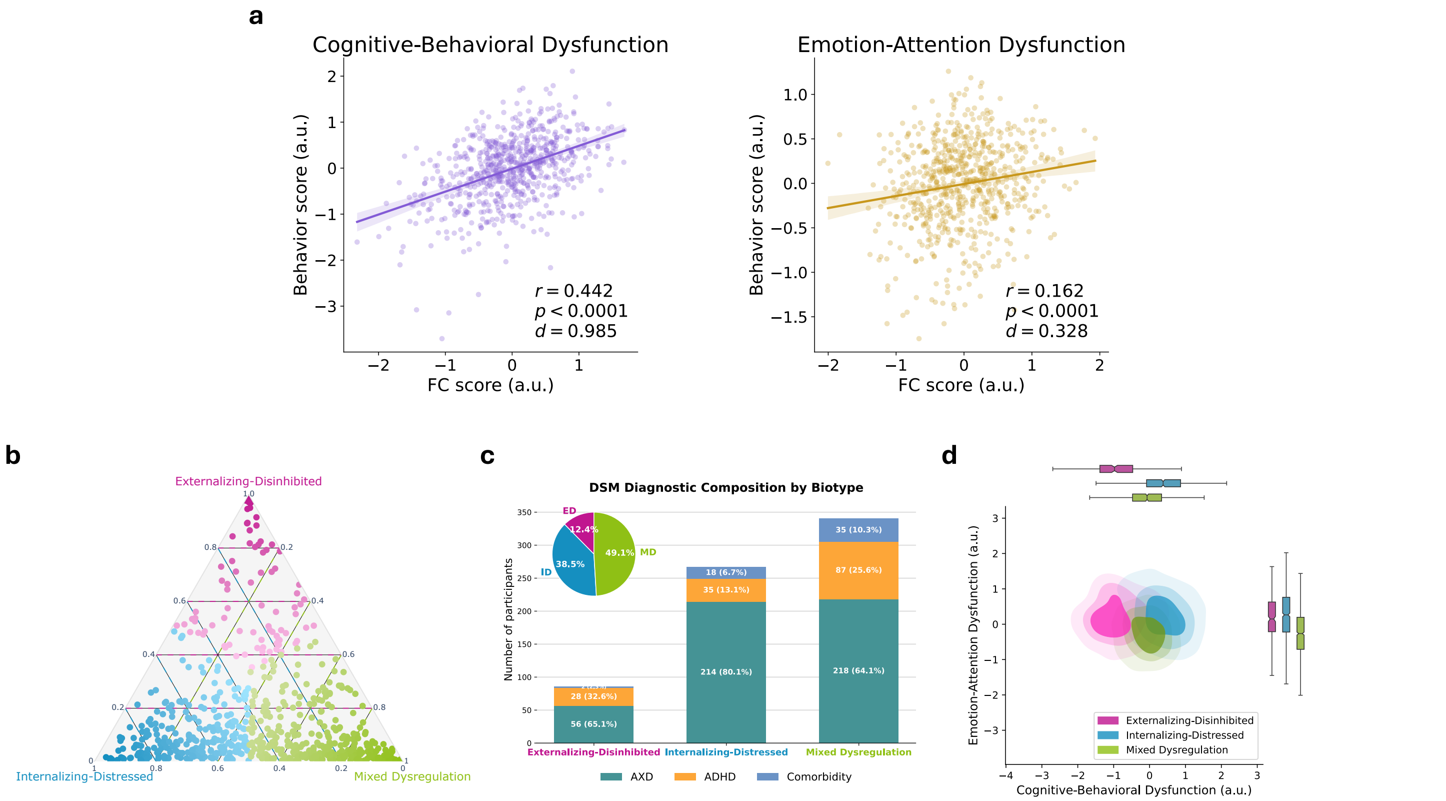

**Figure S26. Group-level comparisons across cognitive, mental health, personality, and contextual domains in ABCD baseline and late-onset participants.** Domain-specific profiles are shown for seven groups: three biotypes identified among children with ADHD or AXD diagnoses at baseline (N=3,508), three biotypes identified among children without a baseline diagnosis but with late-onset ADHD or AXD (N=693), and a control group. **a**, Cognitive performance, including working memory, executive function, processing speed, and fluid intelligence; **b**, Mental health symptoms derived from CBCL subscales; **c**, Personality traits encompassing BIS/BAS and UPPS-P constructs; **d**, Contextual and environmental factors such as sleep patterns, family conflict, and parental psychopathology. Biotypes from baseline-diagnosed individuals and controls are indicated using solid lines; biotypes from the late-onset group are shown with dashed lines. Outliers were removed using a 3-standard-deviation threshold for approximately normally distributed variables and the median absolute deviation method with a threshold of 4 for others. Group differences across the six biotypes (excluding controls) were assessed using the Kruskal–Wallis test, followed by FDR correction ($q < 0.05$) within each domain. Asterisks indicate scales with significant biotype-level differences after correction. Radar positions represent group means; error bars indicate 95% confidence intervals.

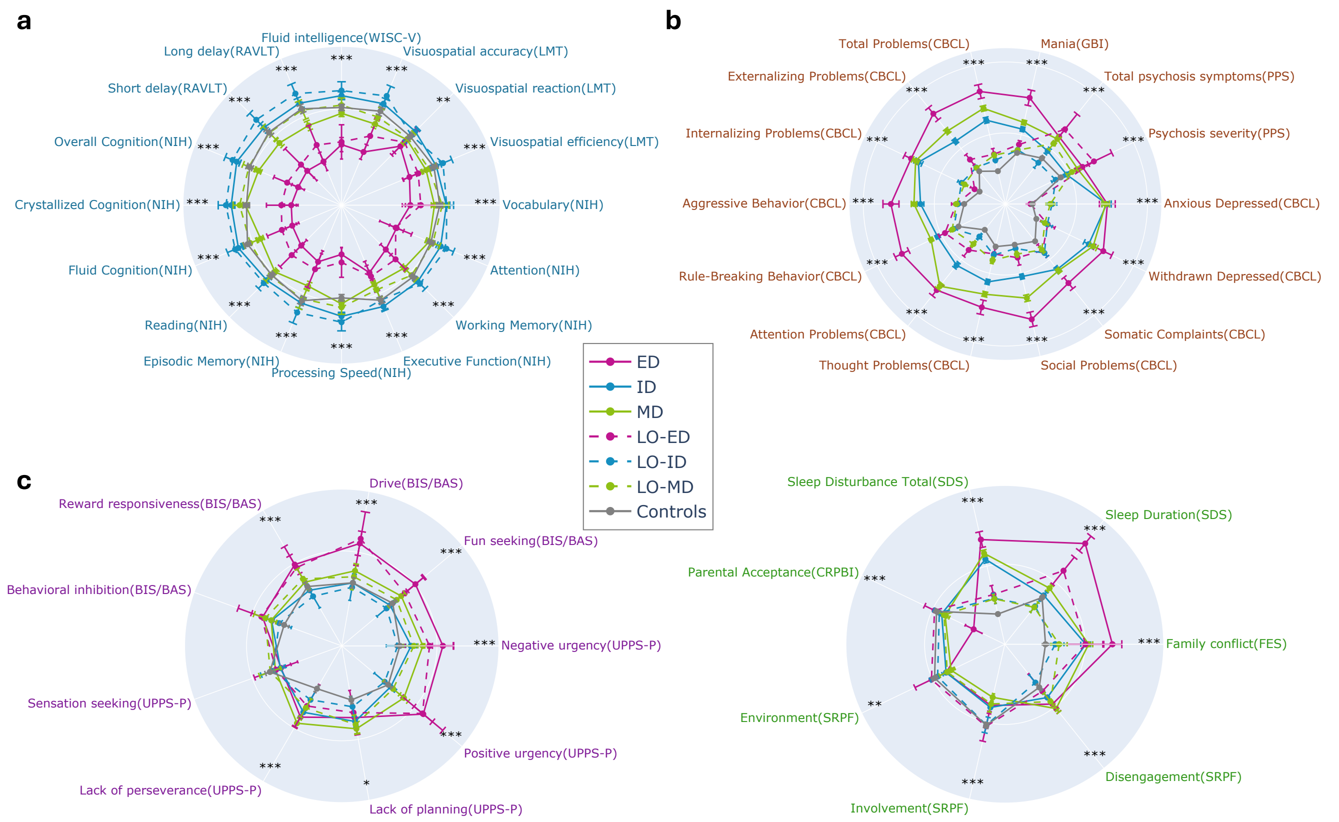

**Figure S27. Structural brain differences across sMRI and DTI modalities among late-onset biotypes and controls.** Pairwise group comparisons for **a** four sMRI metrics (cortical volume, cortical thickness, cortical surface area, and subcortical volume) and **b** four DTI metrics (fractional anisotropy, mean diffusivity, longitudinal diffusivity, and transverse diffusivity). Outliers were excluded within each group using a 3-standard-deviation rule. Linear regression models controlled for age, sex, parental education, family income, marital status, and intracranial volume. Asterisks indicate FDR-corrected significance ($q < 0.05$), Cohen's $d$ effect sizes are shown, and the colors of significance markers and labels denote the group with the higher mean.

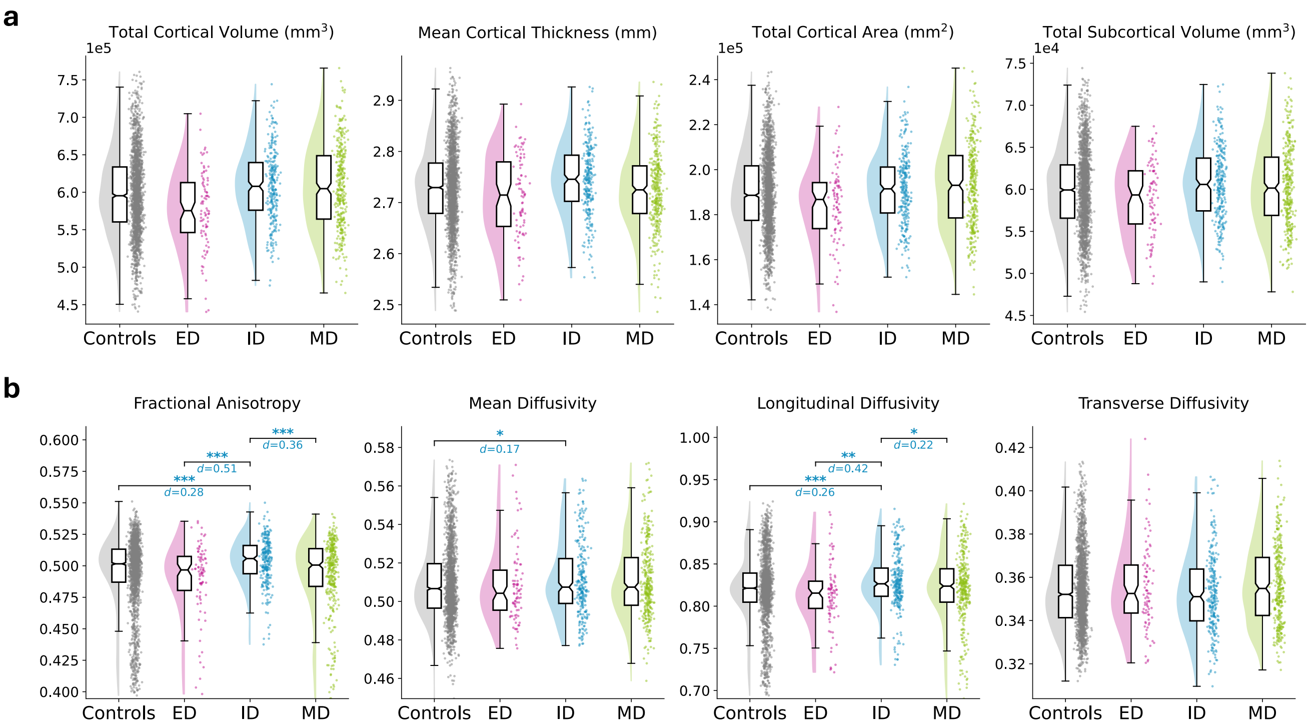

**Figure S28. Longitudinal modeling of CBCL subscales in the ABCD late-onset group.** Results from LME models predicting CBCL T scores across ages 9–15 for children who developed ADHD or AXD diagnoses during follow-up (late-onset group; N=693). **a**, Predicted symptom trajectories based on the LME model, stratified by biotype. Shaded areas represent 95% confidence intervals. FDR-corrected *p*-values for age-by-biotype interaction terms are reported in each panel. Models included sex, parental education, family income, and medication use as covariates. **b**, Observed CBCL T scores across baseline and follow-up visits for each late-onset biotype. Dashed gray lines denote the average trajectory of controls.

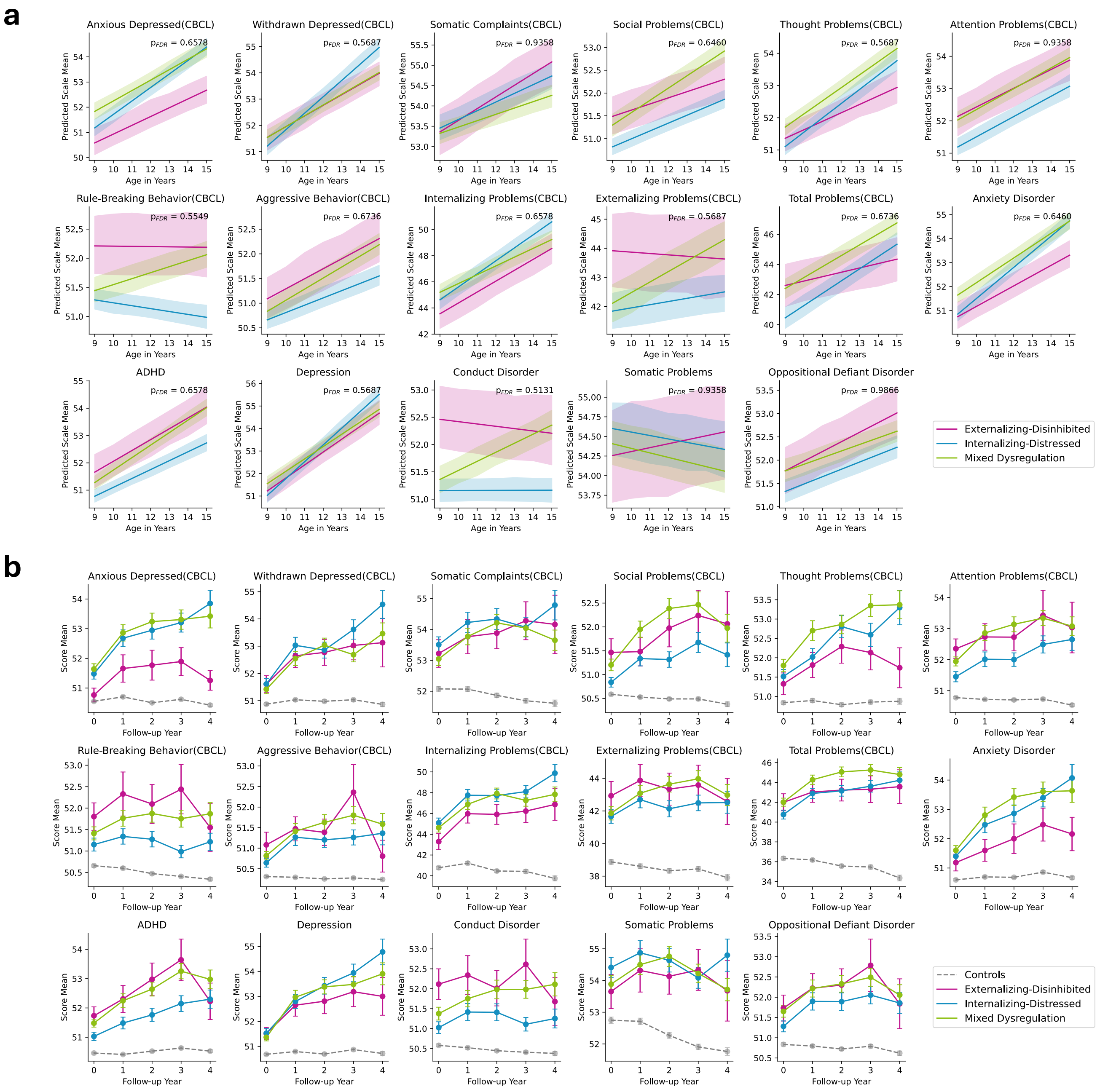

**Figure S29. Longitudinal trajectories of sMRI and DTI metrics in late-onset participants.** Linear mixed-effects models were used to examine age-related changes in brain structure among children who developed ADHD or AXD after baseline (late-onset group; N=693), compared to controls. Predicted trajectories represent model-derived estimates with 95% confidence intervals; observed values are shown across baseline and follow-up visits. FDR-corrected *p*-values for age-by-biotype interactions are reported in each predicted panel. **a**, Predicted trajectories of sMRI metrics (cortical volume, cortical thickness, cortical surface area, and subcortical volume), comparing late-onset and control groups. Models included sex, parental education, family income, and intracranial volume as covariates. **b**, Observed sMRI values across visits; dashed gray lines represent the control mean. **c**, Predicted trajectories of DTI metrics (fractional anisotropy, mean diffusivity, longitudinal diffusivity, and transverse diffusivity), comparing late-onset and control groups. Models included sex and intracranial volume as covariates. **d**, Observed DTI values across visits; dashed gray lines represent the control mean.

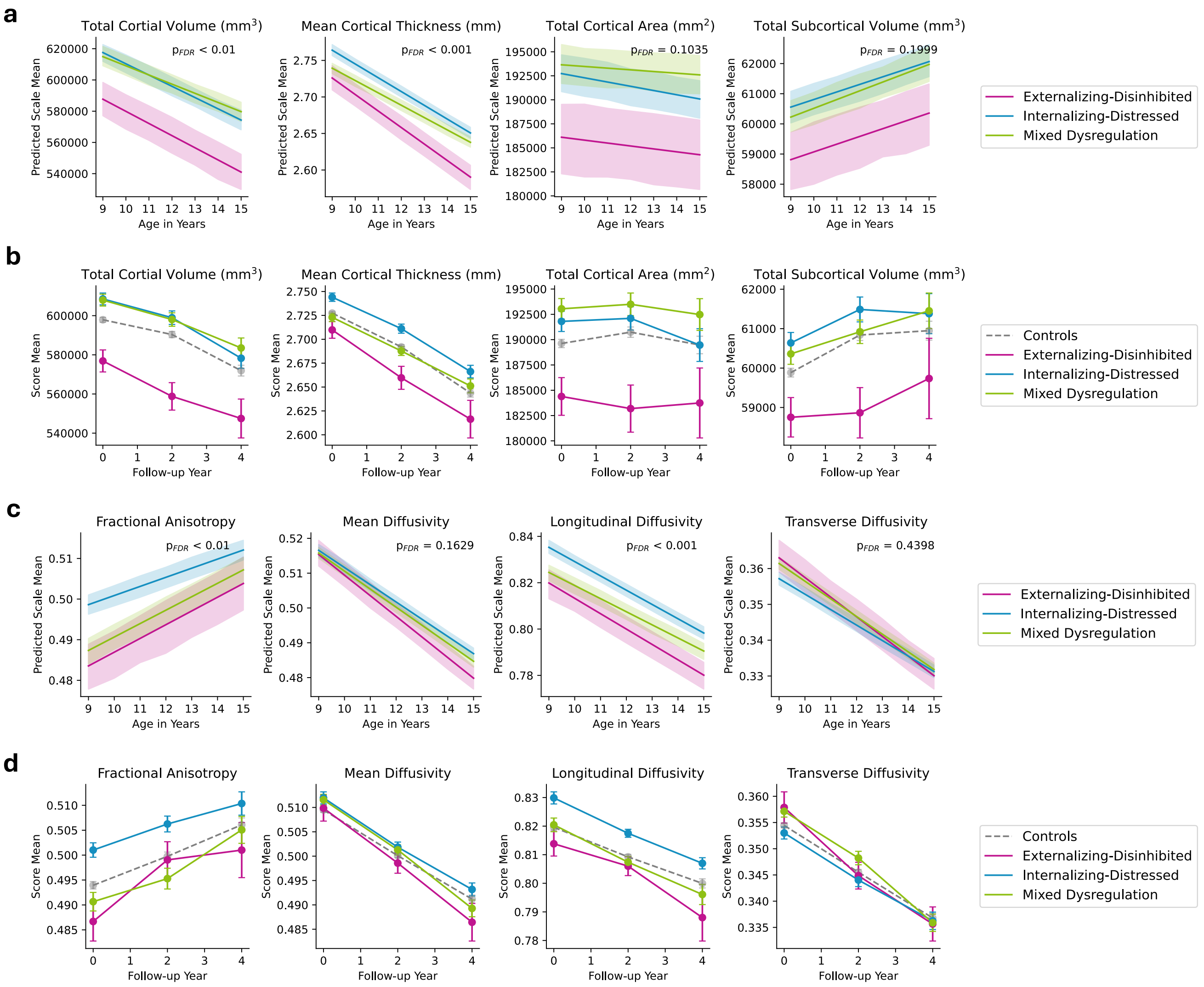

**Figure S30. Neuroimaging differences from controls in DSM-based diagnostic groups.** Individuals with comorbid ADHD and AXD were included in both diagnostic groups. **a**, FC patterns in ADHD and AXD were compared with controls using two-sample $t$-tests at both ROI and network levels, with FDR correction ($q < 0.05$). **b**, Pairwise group comparisons among DSM-based diagnostic groups and controls for four sMRI metrics (cortical volume, cortical thickness, cortical surface area, and subcortical volume). Outliers were excluded within each group using a 3-standard-deviation rule, and linear regression models controlled for age, sex, parental education, family income, marital status, and intracranial volume. No significant group differences were observed after correction. **c**, Pairwise group comparisons among DSM-based diagnostic groups and controls for four DTI metrics (fractional anisotropy, mean diffusivity, longitudinal diffusivity, and transverse diffusivity). Outliers were excluded within each group using a 3-standard-deviation rule, and linear regression models controlled for the same covariates as in **b**. No significant group differences were observed after correction.

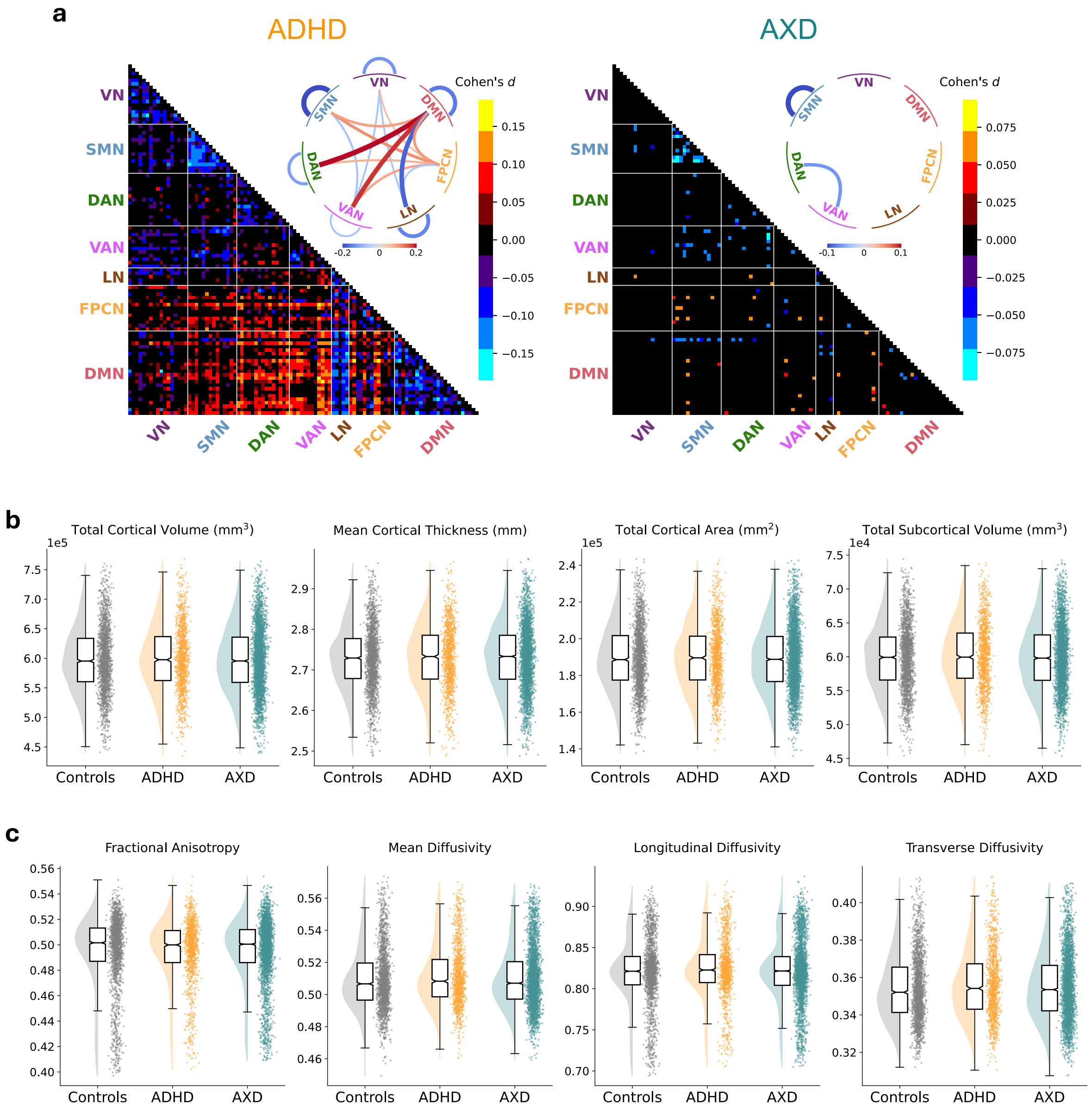

**Figure S31. Comparison between data-driven biotypes and conventional ADHD subtypes in the HBN dataset. a**, HBN individuals diagnosed with ADHD subtypes of hyperactive/impulsive, inattentive, or combined (N=162) across data-driven biotypes and DSM subtypes. **b**, FC patterns in data-driven biotypes and DSM subtypes were compared with controls using two-sample $t$-tests at the ROI level, with FDR correction ($q < 0.05$). **c–d**, Clinical profiles of ADHD individuals grouped by data-driven biotypes (**c**) and DSM subtypes (**d**). Outliers were removed using a 3-standard-deviation threshold for approximately normally distributed variables and the median absolute deviation method with a threshold of 4 for non-normal variables. Group differences were assessed using the Kruskal–Wallis test with FDR correction ($q < 0.05$) across scales. Error bars represent 95% confidence intervals.

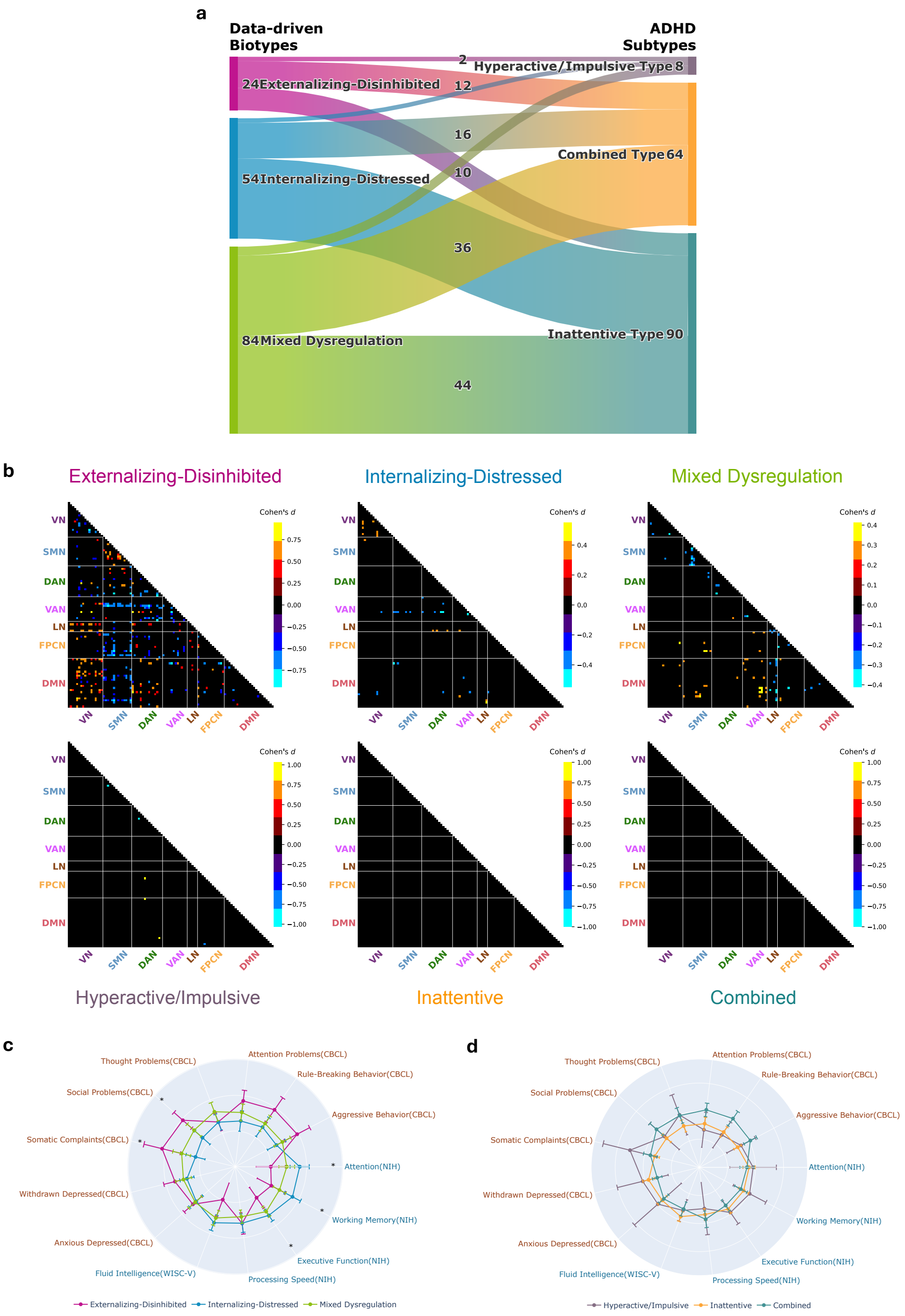

**Figure S32. Clustering model selection for clinical scales.** Bayesian Information Criterion (BIC) and Akaike Information Criterion (AIC) values estimated from 100 repetitions of Gaussian mixture modeling on 80% randomly subsampled data using the 13 standardized clinical scales from diagnosed participants (N=3,508) from the ABCD cohort. Lines represent the mean BIC/AIC across repetitions, and shaded areas indicate the range between minimum and maximum values. Lower values indicate better model fit.

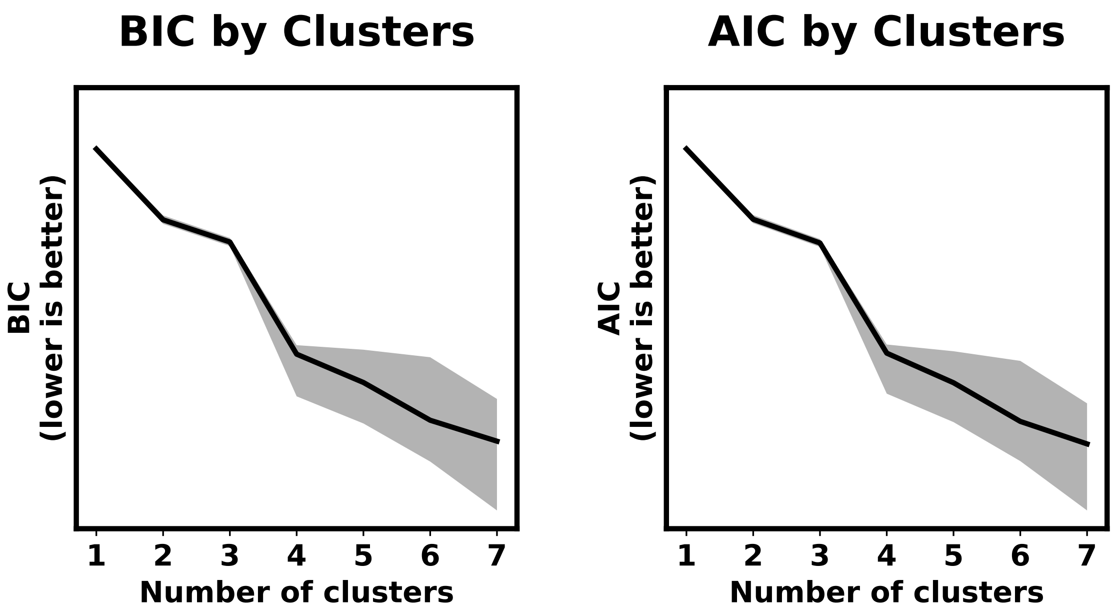

**Figure S33. Clustering model selection and stability evaluation for behavior scores. a**, Bayesian Information Criterion (BIC) and Akaike Information Criterion (AIC) values estimated from 100 repetitions of Gaussian mixture modeling on 80% randomly subsampled data. Lines represent the mean BIC/AIC across repetitions, and shaded areas indicate the range between minimum and maximum values. Lower values indicate better model fit. **b**, Clustering stability scores estimated using 100 repetitions of 5-fold cross-validation with 80% subsampled data. Left panel: Original clustering stability, measured by clustering accuracy (ACC), adjusted Rand index (ARI), and normalized mutual information (NMI), alongside chance-level stability computed by applying the same procedure to randomly shuffled data. Error bars indicate standard deviations across repetitions. Right panel: Adjusted stability scores were calculated as $(\text{Original} - \text{Chance})/(1 - \text{Chance})$ to account for baseline agreement due to chance.

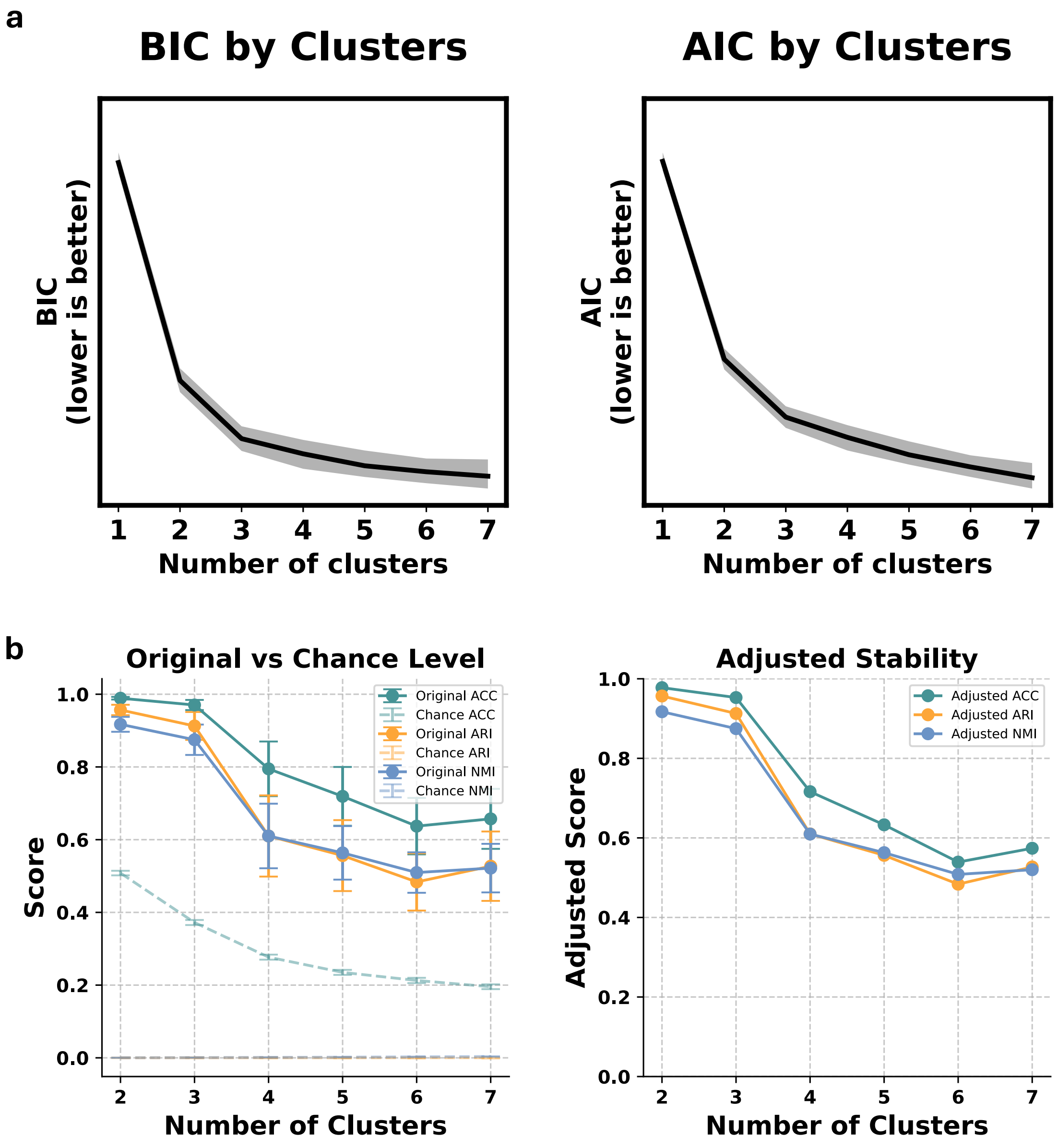

**Figure S34. Biotypes derived from behavior scores in the ABCD (left) and HBN (right) cohorts. a**, Posterior membership probabilities for three biotypes obtained from Gaussian mixture modeling of behavior scores. **b**, DSM diagnostic composition within each biotype. **c**, Clinical profiles across the 13 clinical scales used in model training. **d**, Kernel density estimation plots of biotypes in the two-dimensional canonical behavior score space.

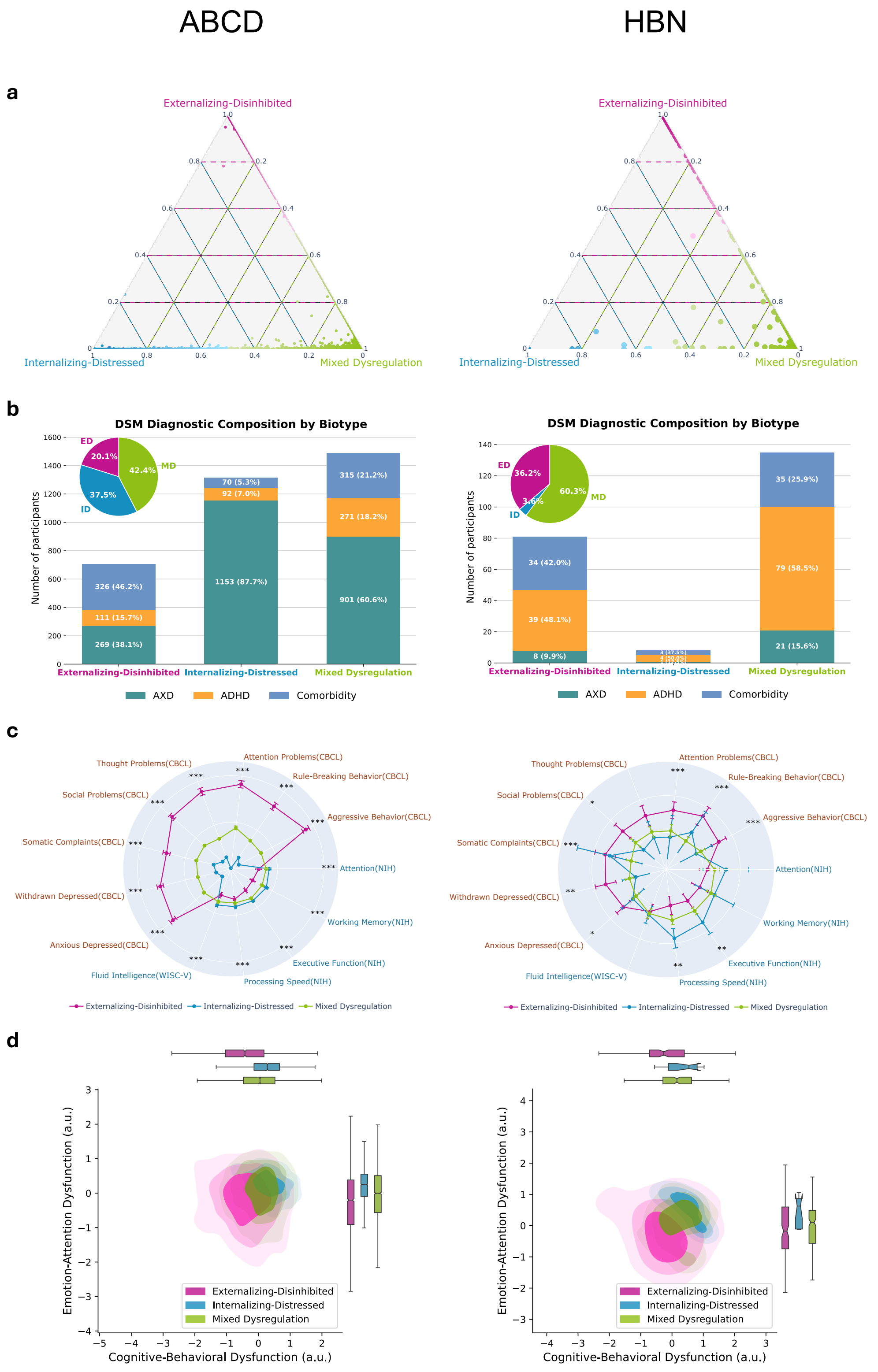

**Figure S35. Atypical functional connectivity patterns in biotypes derived from behavior scores.** For each biotype, FC were compared with the control group using two-sample $t$-tests, followed by FDR correction ($q < 0.05$). Colorbars indicate effect size (Cohen's $d$). Warm colors indicate hyperconnectivity (biotype $>$ control), and cool colors indicate hypoconnectivity (biotype $<$ control). **a**, ABCD cohort. **b**, HBN cohort.

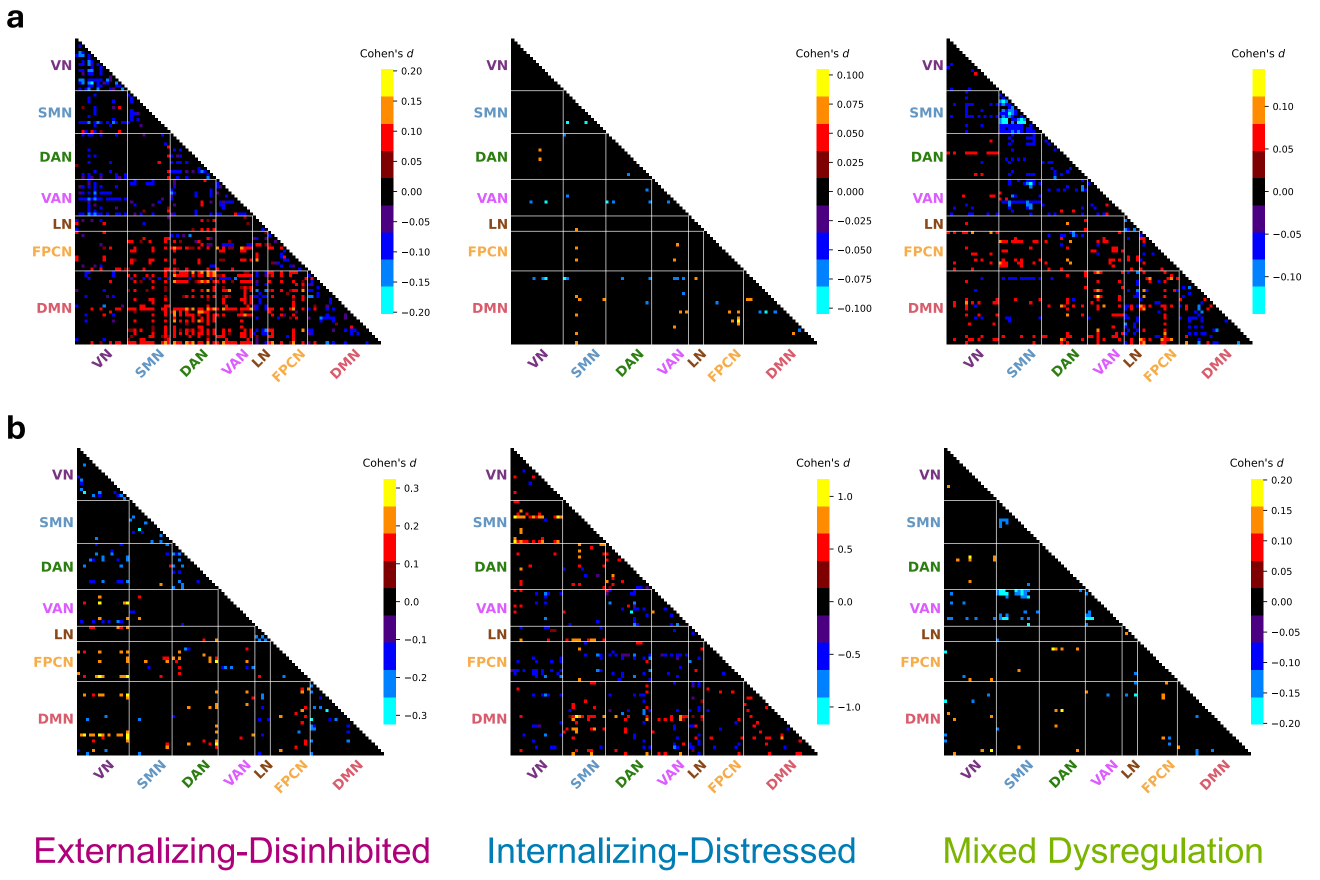

**Figure S36. Data inclusion and exclusion flow for ABCD and HBN cohorts. a**, ABCD participants were selected from the baseline release. After excluding individuals with incomplete rs-fMRI data, missing diagnostic or behavioral information at baseline, or failed MRI preprocessing and quality control, a total of 6,148 children were retained for analysis. Among them, children diagnosed with ADHD or AXD at baseline formed the discovery cohort (N=3,508). A separate late-onset group (N=693) included children with no diagnosis at baseline who developed ADHD or AXD during follow-up. Controls (N=1,947) had no diagnoses at any timepoint and passed all data quality criteria. **b**, HBN participants were filtered based on imaging quality, behavioral data completeness, and age matching to the ABCD cohort. The final diagnostic sample comprised children with ADHD or AXD diagnoses (N=224).

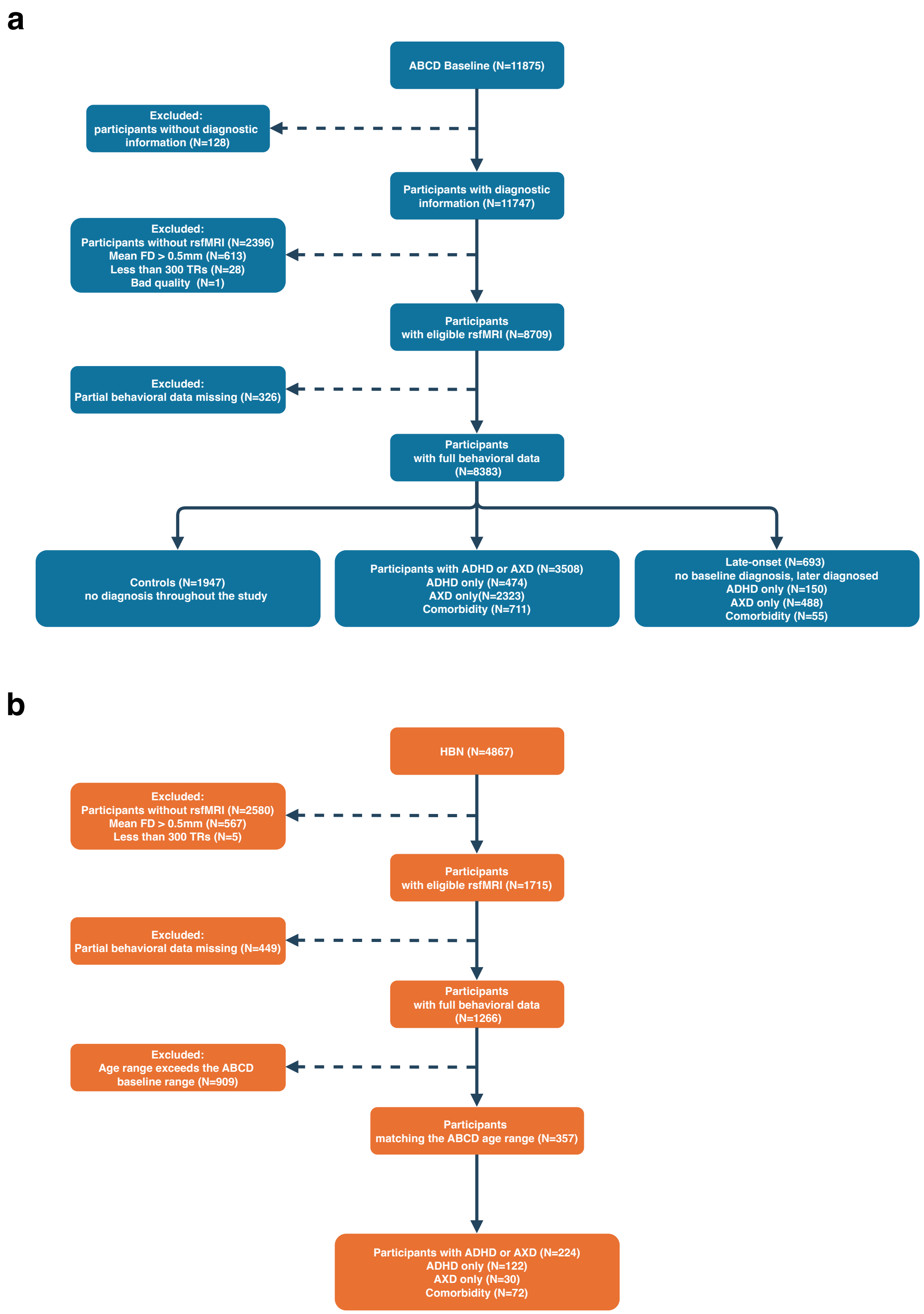

**Figure S37. Correlations across cognitive and mental health scales in ABCD-diagnosed participants.** Pairwise Pearson correlations among 13 clinical scales used in model training, including five cognitive measures (NIH Toolbox and WISC-V) and eight CBCL mental health subscales, computed across diagnosed participants from the ABCD dataset (N=3,508).

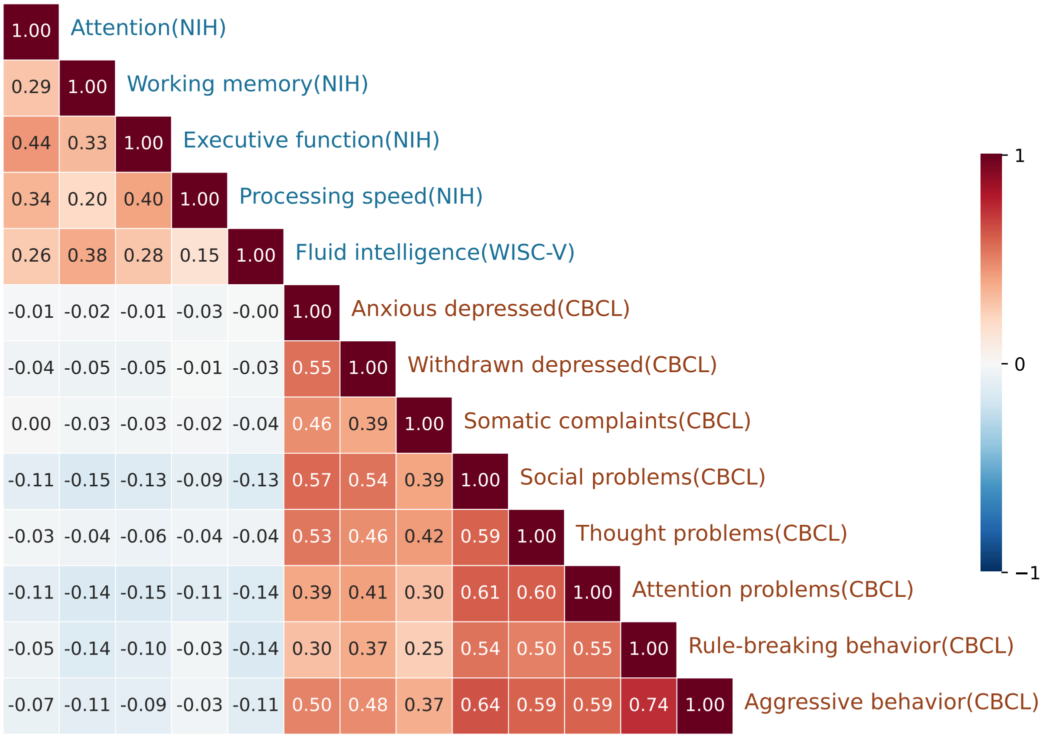

**Figure S38. Training convergence curves of DeCoDE a**, Total loss across training epochs during 10-fold cross-validation of the cVAE. The curve shows fold-averaged values (the shaded band is visually indistinguishable due to minimal variation across folds). **b**, Validation correlation curves for each DGCCA canonical dimension during training. Curves show fold-averaged values; shaded areas represent standard deviation across folds.

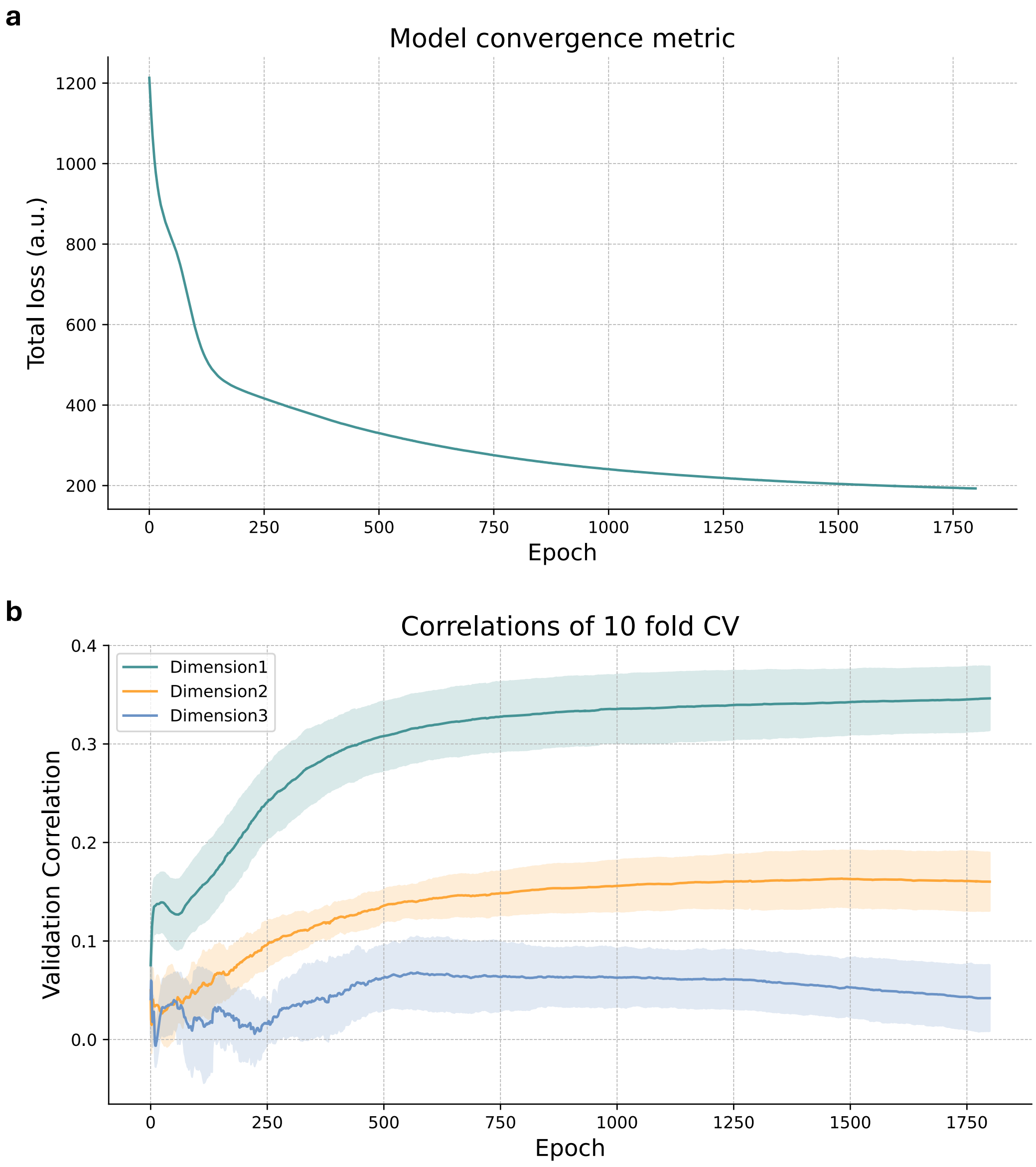

**Table S1. Demographic and clinical characteristics of youth with ADHD or AXD and controls at baseline in the ABCD cohort.**

Demographic and clinical characteristics of controls and ADHD & AXD

| | Controls | Comorbidity | ADHD only | AXD only |
|---|---|---|---|---|
| N | 1947 | 711 (20.3%) | 474 (13.5%) | 2323 (66.2%) |
| Female (%) | 1028 (52.8%) | 264 (37.1%) | 140 (29.5%) | 1227 (52.8%) |
| Age in years (mean ± SD) | 9.95 ± 0.62 | 9.94 ± 0.63 | 9.95 ± 0.60 | 9.95 ± 0.63 |
| Parental education (mean ± SD) | 17.17 ± 2.61 | 17.07 ± 2.30 | 17.17 ± 2.56 | 17.33 ± 2.46 |
| Family income (mean ± SD) | 7.48 ± 2.31 | 6.85 ± 2.44 | 7.22 ± 2.39 | 7.44 ± 2.20 |
| Parents married or cohabiting (%) | 78.4% | 66.1% | 73.9% | 75.7% |
| Pubertal scale (mean ± SD) | 1.71 ± 0.85 | 1.74 ± 0.86 | 1.57 ± 0.78 | 1.80 ± 0.88 |
| BMI (median) | 17.44 | 17.76 | 17.28 | 17.59 |

**Significance marks**: -*** -** -* +* +** +***

**Parental education**: 16=Occupational, Technical, or Vocational; 17=Academic Program; 18=Bachelor's degree; 19=Master's degree; 20=Professional School degree; 21=Doctoral degree.

**Family income**: 5=$25,000 - $34,999; 6=$35,000 - $49,999; 7=$50,000 - $74,999; 8= $75,000 - $99,999; 9=$100,000 - $199,999; 10=$200,000 and greater.

**Table S2. Demographic and clinical characteristics of diagnosed biotypes and controls in the ABCD cohort.**

Demographic and clinical characteristics of controls and diagnosed biotypes

| | Controls | Externalizing-Disinhibited | Internalizing-Distressed | Mixed Dysregulation |
|---|---|---|---|---|
| N | 1947 | 437 (12.5%) | 1257 (35.8%) | 1814 (51.7%) |
| Female (%) | 1028 (52.8%) | 210 (48.1%) | 689 (54.8%) | 732 (40.4%) |
| Age in years (mean ± SD) | 9.95 ± 0.62 | 9.97 ± 0.65 | 10.04 ± 0.62 | 9.88 ± 0.61 |
| Parental education (mean ± SD) | 17.17 ± 2.61 | 15.63 ± 2.68 | 17.60 ± 2.27 | 17.41 ± 2.34 |
| Family income (mean ± SD) | 7.48 ± 2.31 | 5.44 ± 2.59 | 7.70 ± 2.01 | 7.43 ± 2.20 |
| Parents married or cohabiting (%) | 78.4% | 51.7% | 78.6% | 75.1% |
| Pubertal scale (mean ± SD) | 1.71 ± 0.85 | 2.09 ± 0.92 | 1.81 ± 0.87 | 1.65 ± 0.83 |
| BMI (median) | 17.44 | 19.58 | 17.26 | 17.41 |

**Significance marks:** -*** -** -* +* +** +***

**Parental education**: 16=Occupational, Technical, or Vocational; 17=Academic Program; 18=Bachelor's degree; 19=Master's degree; 20=Professional School degree; 21=Doctoral degree.

**Family income**: 5=$25,000 - $34,999; 6=$35,000 - $49,999; 7=$50,000 - $74,999; 8= $75,000 - $99,999; 9=$100,000 - $199,999; 10=$200,000 and greater.

**Table S3. Measures used in the study and score directions.** NIH = National Institutes of Health; RAVLT = Rey Auditory Verbal Learning Test; WISC-V = Wechsler Intelligence Scale for Children, Fifth Edition; LMT = Little Man Task; CBCL = Child Behavior Checklist; GBI = General Behavior Inventory; PPS = Prodromal Psychosis Scale; UPPS-P = Urgency, Premeditation, Perseverance, Sensation Seeking, Positive Urgency Impulsive Behavior Scale; BIS/BAS = Behavioral Inhibition/Behavioral Activation Scales; FES = Family Environment Scale; SRPF = School Risk and Protective Factors; CRPBI = Child Report of Parental Behavior Inventory; SDS = Sleep Disturbance Scale; PDS = Pubertal Development Scale.

| **Measure** | **Source** | **Direction (higher values mean . . . )** |
|---|---|---|
| *Cognition* | | |
| Vocabulary | NIH Toolbox | better performance |
| Attention | NIH Toolbox | better performance |
| Working Memory | NIH Toolbox | better performance |
| Executive Function | NIH Toolbox | better performance |
| Processing Speed | NIH Toolbox | better performance |
| Episodic Memory | NIH Toolbox | better performance |
| Reading | NIH Toolbox | better performance |
| Fluid Cognition | NIH Toolbox | better performance |
| Crystallized Cognition | NIH Toolbox | better performance |
| Overall Cognition | NIH Toolbox | better performance |
| Short delay | RAVLT | better performance |
| Long delay | RAVLT | better performance |
| Fluid intelligence | WISC-V | higher fluid intelligence |
| Visuospatial accuracy | LMT | better performance |
| Visuospatial reaction | LMT | faster reaction |
| Visuospatial efficiency | LMT | higher efficiency |
| *Mental health* | | |
| Anxious/Depressed | CBCL | more problems |
| Withdrawn/Depressed | CBCL | more problems |
| Somatic Complaints | CBCL | more problems |
| Social Problems | CBCL | more problems |
| Thought Problems | CBCL | more problems |
| Attention Problems | CBCL | more problems |
| Rule-Breaking Behavior | CBCL | more problems |
| Aggressive Behavior | CBCL | more problems |
| Internalizing Problems | CBCL | more problems |
| Externalizing Problems | CBCL | more problems |
| Total Problems | CBCL | more problems |
| Mania | GBI | more manic symptoms |
| Total psychosis symptoms | PPS | more psychosis symptoms |
| Psychosis severity | PPS | higher psychosis severity |
| *Personality* | | |
| Negative urgency | UPPS-P | higher negative urgency |
| Positive urgency | UPPS-P | higher positive urgency |
| Lack of planning | UPPS-P | lower premeditation |
| Lack of perseverance | UPPS-P | lower perseverance |
| Sensation seeking | UPPS-P | higher sensation seeking |
| Behavioral inhibition | BIS/BAS | higher behavioral inhibition |
| Reward responsiveness | BIS/BAS | higher reward responsiveness |
| Drive | BIS/BAS | higher drive |
| Fun seeking | BIS/BAS | higher fun seeking |
| *Contextual factors* | | |
| Family conflict | FES | more family conflict |
| Disengagement | SRPF | more disengagement |
| Involvement | SRPF | more involvement |
| Environment | SRPF | more environmental support |
| Parental Acceptance | CRPBI | more parental acceptance |
| Sleep Disturbance Total | SDS | greater sleep disturbance |
| Sleep Duration | SDS | shorter sleep hours |
| Pubertal Stage | PDS | more advanced pubertal stage |

**Table S4. Demographic and clinical characteristics of youth with ADHD or AXD at baseline, those with late-onset diagnoses during follow-up, and controls in the ABCD cohort.**

Demographic and clinical characteristics of controls, diagnosed and late-onset cohorts

| | Controls | Diagnosed | Late-onset |
|---|---|---|---|
| N | 1947 | 3508 | 693 |
| Female (%) | 1028 (52.8%) | 1631 (46.5%) | 361 (52.1%) |
| Age in years (mean ± SD) | 9.95 ± 0.62 | 9.95 ± 0.62 | 9.89 ± 0.63 |
| Parental education (mean ± SD) | 17.17 ± 2.61 | 17.26 ± 2.44 | 17.65 ± 2.22 |
| Family income (mean ± SD) | 7.48 ± 2.31 | 7.29 ± 2.29 | 7.78 ± 2.02 |
| Parents married or cohabiting (%) | 78.4% | 73.5% | 80.2% |
| Pubertal scale (mean ± SD) | 1.71 ± 0.85 | 1.76 ± 0.87 | 1.64 ± 0.83 |
| BMI (median) | 17.44 | 17.57 | 17.49 |

**Significance marks**: -*** -** -* +* +** +***

**Parental education**: 16=Occupational, Technical, or Vocational; 17=Academic Program; 18=Bachelor's degree; 19=Master's degree; 20=Professional School degree; 21=Doctoral degree.

**Family income**: 5=$25,000 - $34,999; 6=$35,000 - $49,999; 7=$50,000 - $74,999; 8= $75,000 - $99,999; 9=$100,000 - $199,999; 10=$200,000 and greater.

**Table S5. Demographic and clinical characteristics of late-onset biotypes and controls in the ABCD cohort.**

Demographic and clinical characteristics of controls and late-onset biotypes

| | Controls | Externalizing-Disinhibited | Internalizing-Distressed | Mixed Dysregulation |
|---|---|---|---|---|
| N | 1947 | 86 (12.4%) | 267 (38.5%) | 340 (49.1%) |
| Female (%) | 1028 (52.8%) | 44 (51.2%) | 154 (57.7%) | 163 (47.9%) |
| Age in years (mean ± SD) | 9.95 ± 0.62 | 9.91 ± 0.67 | 9.97 ± 0.63 | 9.83 ± 0.61 |
| Parental education (mean ± SD) | 17.17 ± 2.61 | 16.20 ± 2.11 | 18.03 ± 2.06 | 17.73 ± 2.23 |
| Family income (mean ± SD) | 7.48 ± 2.31 | 5.90 ± 2.67 | 8.06 ± 1.71 | 8.01 ± 1.83 |
| Parents married or cohabiting (%) | 78.4% | 55.3% | 85.4% | 82.3% |
| Pubertal scale (mean ± SD) | 1.71 ± 0.85 | 1.98 ± 1.01 | 1.62 ± 0.80 | 1.58 ± 0.78 |
| BMI (median) | 17.44 | 19.21 | 17.43 | 17.22 |

**Significance marks:** -*** -** -* +* +** +***

**Parental education**: 16=Occupational, Technical, or Vocational; 17=Academic Program; 18=Bachelor's degree; 19=Master's degree; 20=Professional School degree; 21=Doctoral degree.

**Family income**: 5=$25,000 - $34,999; 6=$35,000 - $49,999; 7=$50,000 - $74,999; 8= $75,000 - $99,999; 9=$100,000 - $199,999; 10=$200,000 and greater.

**Table S6. Cortical parcels from the Schaefer 100-parcel functional atlas annotated with Yeo's 7-network labels.**

| Left Hemisphere | | | Right Hemisphere | | |
|---|---|---|---|---|---|
| **Parcel** | **Region** | **Network** | **Parcel** | **Region** | **Network** |
| 1 | Fusiform | VN | 10 | Fusiform1 | VN |
| 2 | Lingual1 | VN | 11 | Fusiform2 | VN |
| 3 | Lingual2 | VN | 12 | InferiorOccipital1 | VN |
| 4 | MiddleOccipital | VN | 13 | InferiorOccipital2 | VN |
| 5 | Calcarine1 | VN | 14 | Calcarine | VN |
| 6 | Calcarine2 | VN | 15 | Lingual | VN |
| 7 | MiddleTemporal | VN | 16 | MiddleOccipital | VN |
| 8 | SuperiorOccipital | VN | 17 | Cuneus | VN |
| 9 | Cuneus | VN | | | |
| 18 | SuperiorTemporal1 | SMN | 24 | SuperiorTemporal | SMN |
| 19 | Insula | SMN | 25 | Insula | SMN |
| 20 | SuperiorTemporal2 | SMN | 26 | RolandicOperculum | SMN |
| 21 | Postcentral | SMN | 27 | Postcentral | SMN |
| 22 | Precentral | SMN | 28 | Precentral1 | SMN |
| 23 | Paracentral | SMN | 29 | Postcentral1 | SMN |
| | | | 30 | Postcentral2 | SMN |
| | | | 31 | Precentral2 | SMN |
| 32 | InferiorTemporal | DAN | 40 | MiddleTemporal | DAN |
| 33 | InferiorParietal | DAN | 41 | Postcentral | DAN |
| 34 | SuperiorParietal1 | DAN | 42 | InferiorParietal | DAN |
| 35 | PostCentral | DAN | 43 | SuperiorParietal | DAN |
| 36 | Precuneus | DAN | 44 | Precuneus | DAN |
| 37 | SuperiorParietal2 | DAN | 45 | InferiorFrontal | DAN |
| 38 | Precentral | DAN | 46 | SuperiorFrontal | DAN |
| 39 | SuperiorFrontal | DAN | | | |
| 47 | Supramarginal | VAN | 54 | SuperiorTemporal | VAN |
| 48 | PosteriorInsula | VAN | 55 | Supramarginal | VAN |
| 49 | AnteriorInsula | VAN | 56 | Insula | VAN |
| 50 | MiddleFrontal | VAN | 57 | MiddleCingulate | VAN |
| 51 | AnteriorCingulate | VAN | 58 | SupplementaryMotorArea | VAN |
| 52 | MiddleCingulate | VAN | | | |
| 53 | SupplementaryMotorArea | VAN | | | |
| 59 | OrbitalFrontal | LN | 62 | SuperiorOrbitofrontal | LN |
| 60 | TemporalPole | LN | 63 | TemporalPole | LN |
| 61 | InferiorTemporal | LN | | | |
| 64 | InferiorParietal | FPCN | 68 | Supramarginal | FPCN |
| 65 | InferiorFrontal | FPCN | 69 | Angular | FPCN |
| 66 | Precuneus | FPCN | 70 | OrbitalMiddleFrontal | FPCN |
| 67 | MiddleCingulate | FPCN | 71 | InferiorFrontal | FPCN |
| | | | 72 | MiddleFrontal(BA9) | FPCN |
| | | | 73 | MiddleFrontal(BA8) | FPCN |
| | | | 74 | MiddleCingulate | FPCN |
| | | | 75 | AnteriorCingulate | FPCN |
| | | | 76 | Precuneus | FPCN |
| 77 | InferiorTemporal | DMN | 90 | Angular | DMN |
| 78 | MiddleTemporal1 | DMN | 91 | InferiorTemporal | DMN |
| 79 | MiddleTemporal2 | DMN | 92 | SuperiorTemporal | DMN |
| 80 | Angular | DMN | 93 | MiddleTemporal | DMN |
| 81 | InferiorOrbitofrontal | DMN | 94 | OrbitalInferiorFrontal | DMN |
| 82 | InferiorFrontal | DMN | 95 | InferiorFrontal | DMN |
| 83 | MedialOrbitofrontal | DMN | 96 | AnteriorCingulate | DMN |
| 84 | MiddleFrontal | DMN | 97 | SuperiorMedialFrontal | DMN |
| 85 | SuperiorMedialFrontal | DMN | 98 | MiddleFrontal | DMN |
| 86 | CaudalMiddleFrontal | DMN | 99 | Precuneus | DMN |
| 87 | RostralMiddleFrontal | DMN | 100 | PosteriorCingulate | DMN |
| 88 | Precuneus | DMN | | | |
| 89 | PosteriorCingulate | DMN | | | |
